# Reproductive morph specialisation facilitated by a maternal sex-determining region in a fungus gnat (*Bradysia coprophila*)

**DOI:** 10.64898/2026.04.09.717474

**Authors:** Melany Henot, Robert B Baird, Findlay Duncan, Jasmine SY Lee, Laura Ross

## Abstract

Sexual reproduction is a ubiquitous process in eukaryotes, yet mechanisms of sex determination are strikingly diverse. One unique and understudied system is maternal genetic sex determination (*mat*-GSD), in which the genotype of the mother determines offspring sex. In some species, mothers further specialise into genetically determined reproductive morphs: gynogenic females that produce all-female broods, and androgenic females that produce all-male broods. Although this partition echoes the evolution of separate, dimorphic sexes, it remains unclear whether the morphs diverge in traits beyond offspring sex determination, and how this affects the evolutionary dynamics of the mating system. Here we address these questions in the dark-winged fungus gnat *Bradysia coprophila*, in which female morphs are determined by a large X-linked inversion. We show that maternal reproductive morphs diverge significantly in life history traits and gene expression profile, suggesting adaptive specialisation into their reproductive roles. We further evaluate potential drivers for specialisation and test two of them empirically, showing evidence in line with sex-specific maternal provisioning. By drawing explicit parallels with sex chromosome evolution and sexual dimorphism, our results extend core principles of reproductive specialisation to *mat*-GSD systems, and underscore the potential of unusual reproductive systems for extending fundamental evolutionary theory on how selection and genomic architecture interact to shape mating system evolution.

## 1. Introduction

Maternal genetic sex determination (*mat*-GSD) is a unique and understudied sex-determining system, in which zygotic sex is determined by maternal genotype, rather than that of the zygote. It has independently evolved in three speciose groups of Dipteran insects: Sciaridae (dark-winged fungus gnats), Cecidomyiidae (gall midges), and Calliphoridae (blowflies) (Baird et al., 2023a). Outside of Diptera, examples occur in bivalves (Kenchington et al., 2002; Ghiselli et al., 2012), ants (Lagunas-Robles et al., 2021), and *Daphnia* (Ye et al., 2019). In some species, females produce mixed-sex broods (digenic), while in others, they are further distinguished as one of two genetically-determined reproductive morphs with specialised offspring sex ratio—gynogenic females produce exclusively female offspring, and androgenic females exclusively male offspring (Baird et al., 2023a). Such specialisation is analogous to the evolution of separate sexes, but leads to three, rather than two, stable and co-existing reproductive morphs. Yet, the foundational question of whether these morphs have distinct phenotypes remains unexplored—do the morphs differ in anatomy, physiology and behaviour beyond offspring sex determination, or are they otherwise undifferentiated? If divergence exists, is it driven by morph-specific selection and adaptive specialisation, or by constraints and fitness costs associated with the genetic architecture underlying morph identity? Untangling these alternatives is particularly important where morph identity is controlled by large non-recombining regions (supergenes), which can both facilitate the accumulation of morph-beneficial alleles and experience ineffective selection and degeneration.

Unlike *mat*-GSD, the evolution of reproductive specialisation is well-documented in canonical sexual reproduction (Bachtrog et al., 2014). Separation into male and female reproductive morphs, each producing one gamete type, is frequently accompanied by the evolution of secondary sexual traits in behaviour, physiology and life history (Glucksmann, 1974; Andersson, 1994; Mank, 2009; Beukeboom et al., 2015). This allows the sexes to occupy distinct reproductive roles (Schärer et al., 2012; Lehtonen et al., 2016), which can contribute to the stability of the mating system (Charlesworth and Charlesworth, 1978; Hadjivasiliou and Pomiankowski, 2016). At the same time, the scope for sexual dimorphism depends on the underlying sex determination mechanism (Williams and Carroll, 2009). In GSD systems, loci for secondary sexual traits can be linked to the sex determination (SD) locus, by recombination suppression, into a supergene encoding for dimorphic sexes. In fact, selection for sex-specific adaptations is thought to be itself a driver of recombination suppression, leading to heteromorphic sex chromosomes (Rice, 1987; Charlesworth et al., 2005; Flintham and Mullon, 2026). In plants, theory and empirical data show that associations between sex allocation and sexually antagonistic traits can promote specialisation in the transition from hermaphroditism to dioecy (Charlesworth, 2019; Lesaffre et al., 2024). Together, this points to a cyclical relationship between sex determination mechanism and secondary sexual trait evolution—selection on sexual dimorphism can select for tight genetic linkage between the sex-determining locus and secondary traits, while the genetic basis of mating type determination constrains the degree of specialisation that can evolve—shaping the evolution and stability of mating systems. As most prior work has been done on taxa with conventional, zygote-based genetic sex determination (e.g., XY/ZW systems), how specialisation proceeds in alternative sex determination systems remains poorly characterised; to our knowledge, no studies have yet addressed these questions under *mat*-GSD.

In this study, we focus on the monogenic dark-winged fungus gnat *Bradysia coprophila* (Box 1). Across fungus gnats, monogeny appears to have repeatedly evolved from digeny (Baird, 2024), paralleling multiple independent evolutions of divergent sexes across eukaryotes. Among *mat*-GSD species, its genetic basis is best resolved in fungus gnats, enabling tests of whether genomic architecture promotes or hinders morph specialisation. While the offspring sex ratio in digenic species likely has a polygenic basis (Shlyakonova et al., 2026), monogeny seems to be accompanied by repeated evolution of supergenes. In *B. coprophila*, this region has been identified cytological and genomically as a large X-linked inversion (X’), heterozygous in gynogenic females (Metz, 1938; Carson, 1944; Crouse, 1960; Baird et al., 2023b). The morph-limited, permanently-heterozygous, and non-recombining nature of the X’ makes it a prime spot for the accumulation of morph-beneficial alleles. Therefore, we may expect the supergene to facilitate morph-specific divergence. However, recombination suppression and the resulting Hill-Robertson interference leads to degeneration (Hill and Robertson, 1966; Charlesworth and Charlesworth, 2000; Bachtrog et al., 2011), potentially sped up by the low effective population size (N_e_)—one-sixth that of an autosome—of the X’ inversion (Hitchcock et al., 2024). The accumulation of deleterious mutations and repeats is expected to have a negative effect on the fitness of the carrier (Nguyen and Bachtrog, 2021; Teoli et al., 2023). Consistent with this, there is evidence of gene degeneration in the X’ of *B. coprophila* despite its relatively young history (<0.5 Mya) (Baird et al., 2023b), which may predict lower gynogenic female fitness. With conflicting evolutionary forces acting on this region, it is unclear if it facilitates morph-specific divergence or hinders it by being a region of ineffective selection, and how this dynamic contributes to the rapid turnover between monogenic and digenic systems in the family of flies.

Here, using two laboratory populations of *Bradysia coprophila*, we quantify phenotypic divergence between female reproductive morphs, and ask whether it is driven by adaptive specialisation into distinct morphs, or rather by degradation in X’ supergene. We find that reproductive morphs diverge significantly in multiple measures of life history, fitness-related and gene expression traits. Contrary to predictions based on supergene degradation alone, gynogenic females were not less fit than androgenic females. Together with evidence for gynogenic-biased expression, this suggests that morph divergence has an adaptive basis. We tested two potential underlying mechanisms—release from sexually antagonistic selection (SAS) and morph-specific offspring provisioning—and found support for the latter. Our results are consistent with a process of morph specialisation over short evolutionary timeframes, while longer-term supergene degradation could still lead to the high turnover between mating systems in fungus gnats. More broadly, parallels with the evolution of distinct sexes/mating types—origins from undifferentiated ancestors, associated non-recombining regions, and repeated evolution—position this system as a model for reproductive specialisation and the dynamics of its genomic architecture, beyond the conventional context of males and females.

### Box 1: Sex determination mechanism in fungus gnats

In both monogenic and digenic species of dark-winged fungus gnats (Family: Sciaridae), sex determination occurs by elimination of the paternal X chromosome in early embryonic soma (Metz, 1938). All embryos start out with a diploid number of autosomes but three X chromosomes—two of paternal origin (due to meiotic nondisjunction) and one of maternal origin. In the 7-9th cleavage division of early embryogenesis, embryos destined to become female eliminate one of the paternal Xs (producing XX females), while those destined to become male eliminate both paternal Xs (producing XO males) (Phalle and Sullivan, 1996). Since X elimination in the embryonic soma occurs before the onset of zygotic gene activation, the sex of the offspring and number of X chromosomes eliminated is under maternal control.

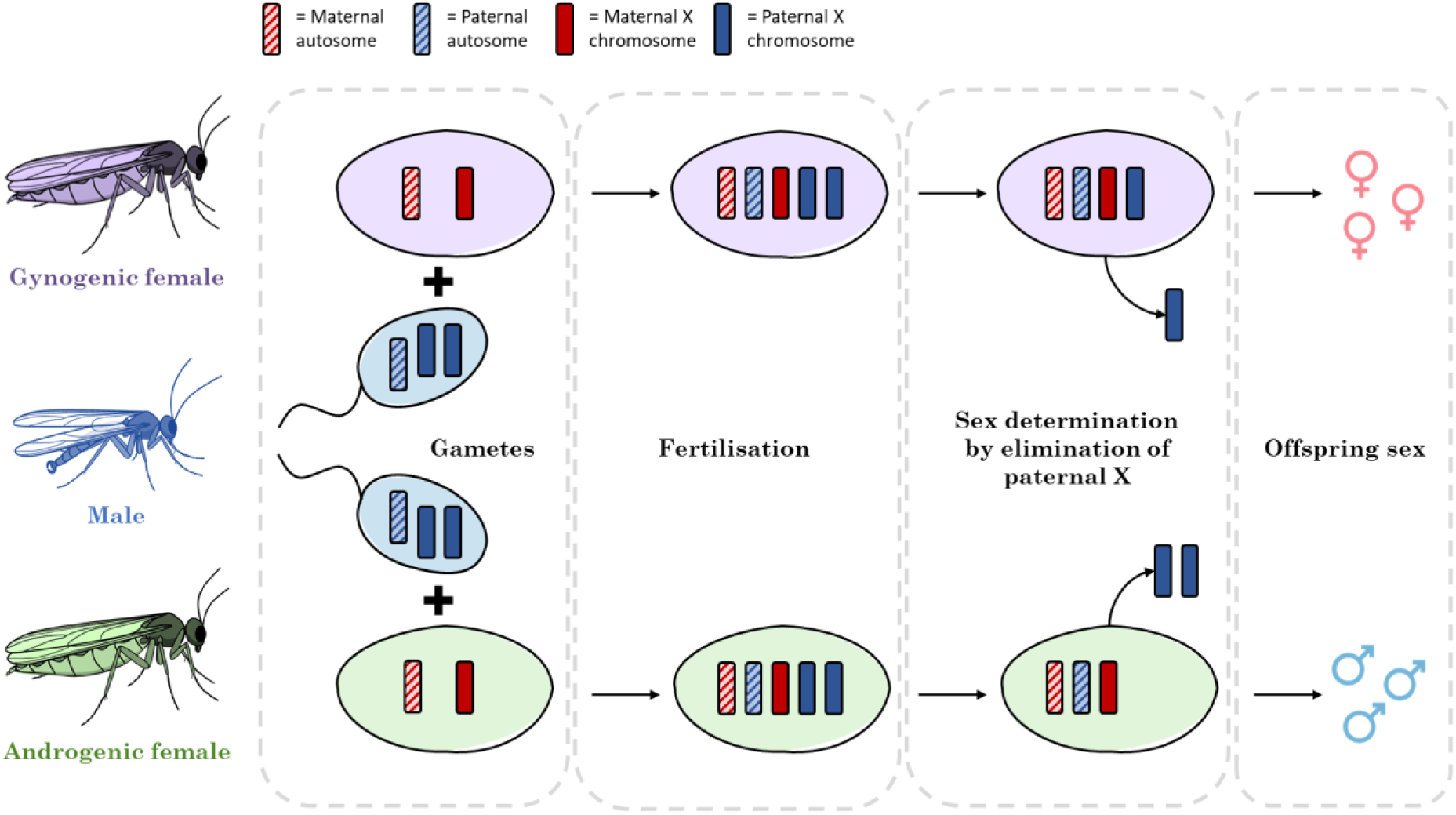

For an extensive overview of fungus gnat sex determination and other aspects of their fascinating biology, see (Gerbi, 1986, 2022).

## 2. Results

### 2.1 Female reproductive morphs differ significantly at the phenotypic level

We compared gynogenic and androgenic females across a number of key life history traits, in two distinct lines of *Bradysia coprophila* (H2 and KM). H2 (Holo2) is a long-established lab line (Metz and Smith, 1931) with a wing marker that distinguishes the otherwise visually identical female reproductive morphs (Gerbi, 2024) (Fig. 7). However, its long laboratory history, which includes exposure to irradiation, may accelerate degeneration of the non-recombining X’ region. To avoid laboratory-line artefacts, we supplemented our study with the KM line, collected from East Lothian, Scotland in February 2023.

Using both lines also permits us to disentangle the effects of heterozygote advantage and sex-specific specialisation. Both lines being relatively inbred, the permanently heterozygous, gynogenic-specific X’ (>50 Mb in a ∼300 Mb genome) elevates genome-wide heterozygosity of gynogenic females and can mask recessive deleterious mutations, potentially making gynogenic females appear fitter without any additional specialisation mutations. To control for this, we generated an outbred line (H2 females x KM males) to normalise genome heterozygosity between morphs.

Across the two inbred and one outbred line, linear/generalised linear mixed-effects models revealed significant divergence between the two reproductive morphs in a number of key life history traits (Fig. 1). The direction of divergence can depend on the line. For developmental rate, gynogenic females developed faster than androgenic females in H2 (*t* = -3.997, *p* = 7.39e^-05^) and H2xKM (*t* = -2.931, *p* = 0.00559) lines, while the opposite was true in the KM line (*t* = 7.155, *p* = 3.07e^-12^) (Fig. 1A). Similarly, gynogenic females were significantly more fecund than androgenic females in the H2 line (*z* = 3.491, *p* = 0.000481), while the opposite was true in the KM line (*z* = -3.550, *p* = 0.000386) (Fig. 1E).

**Figure 1.**
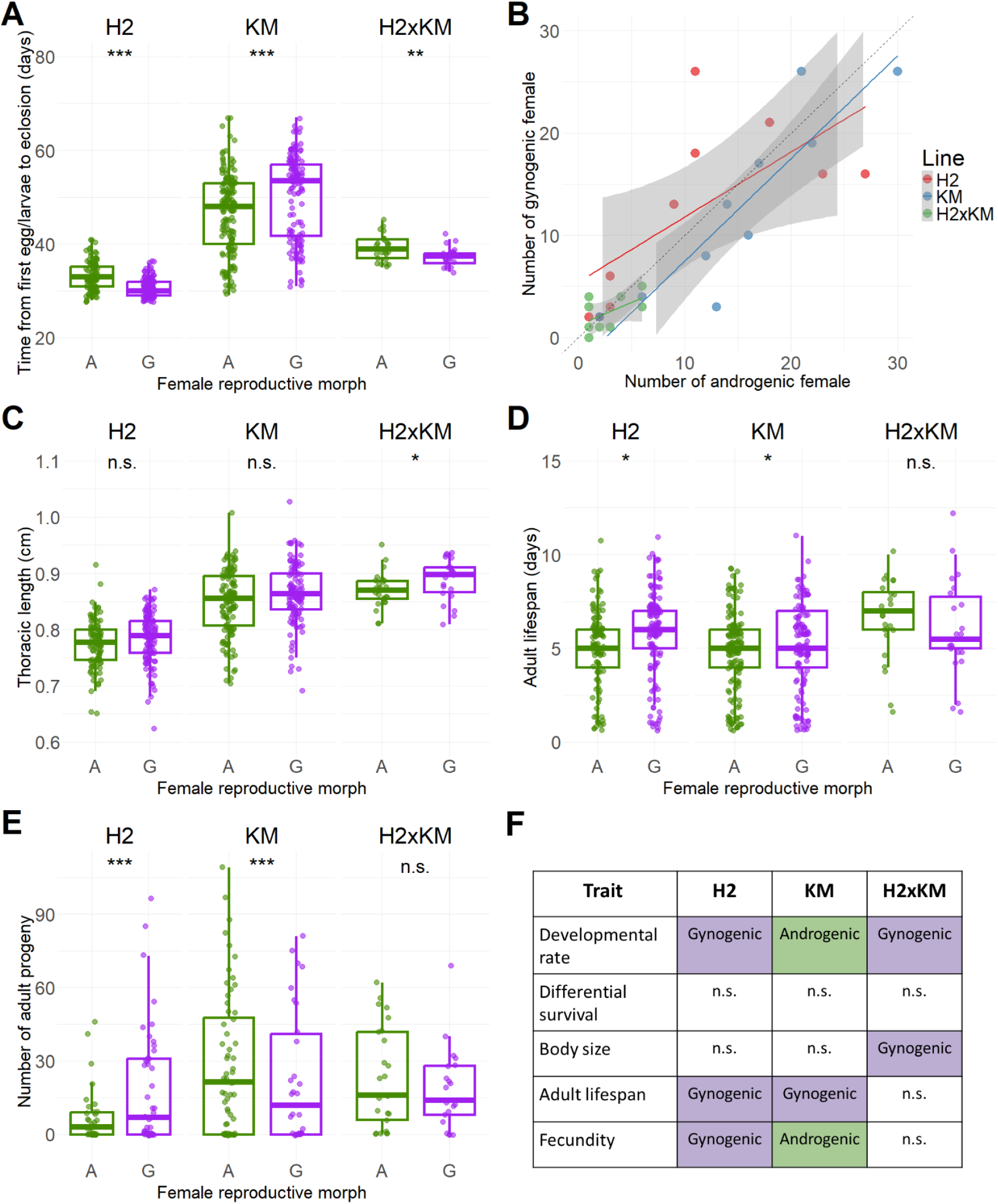
A comparison between gynogenic and androgenic females in two inbred lines (H2 and KM) and one outbred line (H2xKM), in the phenotypic traits of: **(A)** Developmental time, measured as the number of days taken from the hatching of the first larvae in the clutch to individual eclosion as adults in the pure lines, and the number of days from the laying of the first egg in the clutch to individual eclosion as adults in the outbred line. **(B)** Differential survival, measured as the number of adult gynogenic and androgenic females per clutch. **(C)** Adult size, measured by adult thoracic length. **(D)** Adult lifespan, measured as the number of days between eclosion and death. **(E)** Fecundity, measured as the number of adult progeny. **(F)** Summarises whether gynogenic or androgenic females are fitter in the traits measured in each line. For all plots, *** = p < 0.001, ** = p < 0.01, and * = p < 0.05.

Other traits showed greater concordance across lines. In both H2 and KM lines, gynogenic females had significantly longer adult lifespans (*t* = 2.186, *p* = 0.02931) (Fig. 1D). For body size, while no significant difference was found between the morphs in the H2 and KM lines, gynogenic females were significantly larger than their androgenic counterparts in the H2xKM line (*t* = 2.021, *p* = 0.0498) (Fig. 1C). Lastly, no significant differences in survival to adulthood were found in any lines (Fig. 1B).

The pattern of divergence also challenges the null hypothesis of divergence due to a deleterious inversion in gynogenic females, in which gynogenic females are expected to be less fit. On the contrary, androgenic females are fitter than gynogenic females in only two instances of the five traits measured across all three lines (Fig. 1F); gynogenic females are overall just as fit, if not more fit, than androgenic females. As this pattern was maintained in the H2xKM line, this result cannot be fully explained by heterozygote advantage.

### 2.2 Female reproductive morphs diverge significantly in gene expression

To see if the observed phenotypic divergence is accompanied by divergence in gene expression, we compared transcriptomic data at different tissue levels—germline (ovaries), somatic reproductive (ovipositor, accessory glands, spermatheca), and somatic non-reproductive (all other tissue). Since both lines showed phenotypic divergence, only the H2 line, which had more comprehensive genetic resources, was used in this analysis.

#### 2.2.1 Significant gene expression divergence in X-linked genes

Given that maternally acting SD loci in fungus gnats are thought to be X-linked (on the X or X’) (Gerbi, 1986), some morph-specific expression was expected, especially in germline tissue where they should be acting. Supporting this, principal components analysis (PCA) showed samples first clustering by sex (male vs female), and subsequently by female morph identity (androgenic vs gynogenic). Differential gene expression (DGE) analysis of X-linked genes between gynogenic and androgenic females revealed significant differential expression in germline tissue (Fig. 2A, Fig. 2D, Table 1). Unexpectedly, we also see numerous differentially expressed genes (DEGs) in somatic reproductive (Fig. 2B, Fig. 2E, Table 1) and somatic non-reproductive tissues (Fig. 2C, Fig. 2F, Table 1). While differential expression in somatic reproductive tissue (e.g. accessory glands) may contribute to offspring sex determination, this is unlikely for somatic non-reproductive tissue. To clarify whether this dynamic reflects allele-biased expression in X and X’ gametologs, we carried out allele-specific expression analysis in gynogenic females.

**Figure 2.**
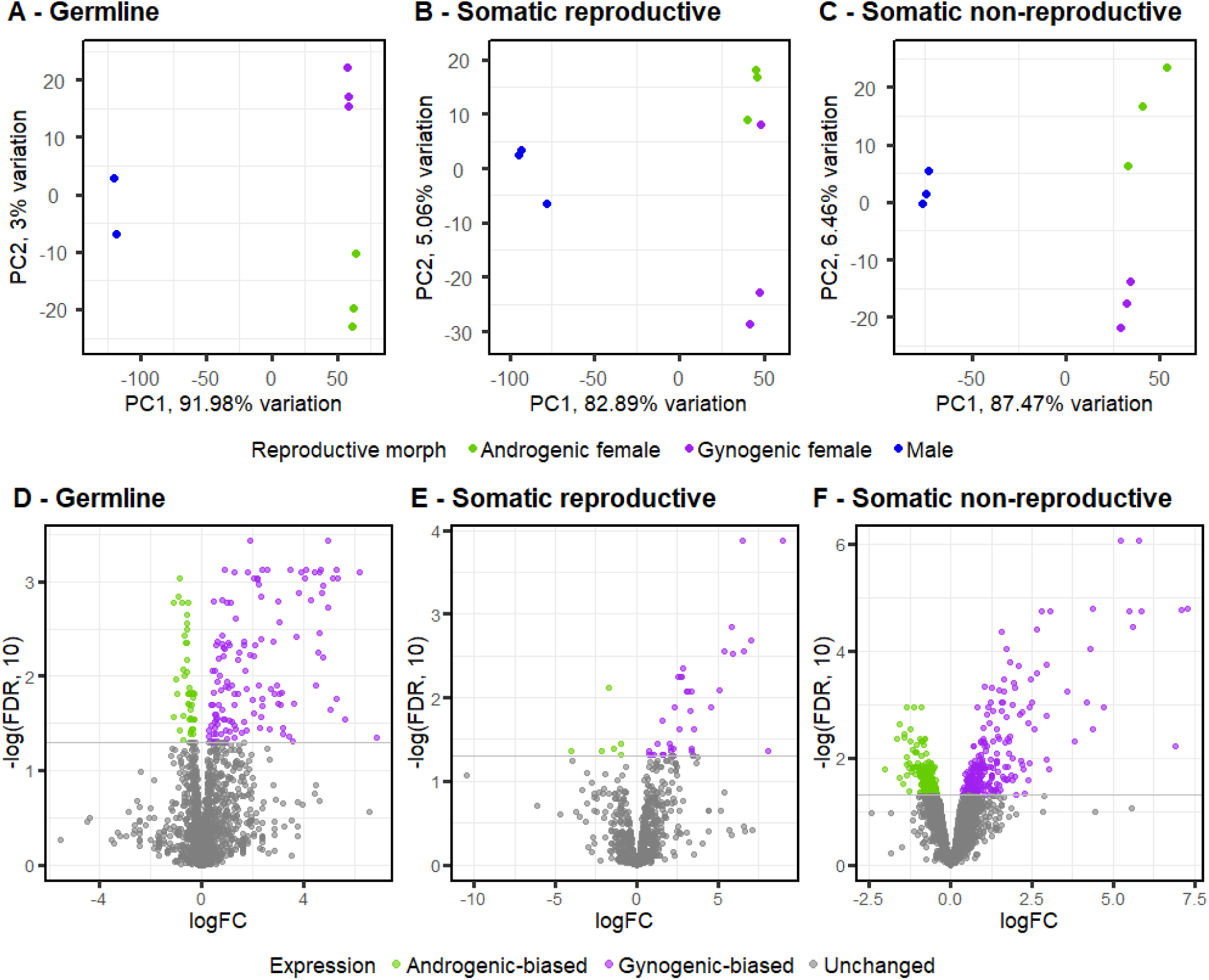
**(A-C)** Principal component analysis of X-linked gene expression in germline, somatic reproductive tissue, and somatic non-reproductive tissue, respectively. Samples cluster first by male (blue) vs female, with subsequent clustering by female reproductive morph (purple for gynogenic, green for androgenic). **(D-F)** Volcano plots demonstrating differential gene expression in germline, reproductive tissue, and somatic non-reproductive tissue, respectively. The x-axis is the log fold-change of gene expression between gynogenic and androgenic females, using androgenic females as a baseline. Values >0 indicate gynogenic-biased expression, while values <0 indicate androgenic-biased expression. The y-axis shows the -log_10_ of the false discovery rate (FDR), a p-value adjusted for multiple testing. The grey horizontal line is drawn at -log_10_(0.05), and genes above that line are coloured (purple for gynogenic and green for androgenic) to signify significance.

**Table 1.** Summary table of the number of X-linked (X or X’) genes with significant differential expression between androgenic and gynogenic females. This is shown separately for germline, somatic reproductive, and somatic non-reproductive tissue.

| Tissue \ X'X expression | Androgenic-biased | Unbiased | Gynogenic-biased |
| --- | --- | --- | --- |
| **Germline** | 44 | 2072 | 135 |
| **Somatic reproductive** | 6 | 2161 | 37 |
| **Somatic non-reproductive** | 211 | 1882 | 243 |

#### 2.2.2 Some X’ alleles are upregulated despite degradation on the inversion

We asked whether the observed morph-biased X-linked expression is linked to allele-specific expression between X and X′ gametologs in gynogenic females. A beta-binomial GLM for read count per X and X’ gametolog pair revealed that most differentially expressed alleles in gynogenic females had X-biased expression. This is in line with known X’ degradation (Baird et al., 2023b), which commonly leads to gene silencing (Mira and Pushker, 2005). However, we also see a significant number of X’-gametolog biased genes across all tissue types (Table 2, Fig. 3A-C). Furthermore, biased expression is associated with differences in molecular evolution rates; mapping to an outgroup, *B. odoriphaga*, which lacks an X’ region, we calculated the ratio of non-synonymous to synonymous nucleotide substitutions (dN/dS), and compared the calculated value between the two gametologs (dN/dS of X gametolog - dN/dS of X’ inversion gametolog). A Welch ANOVA with Games-Howell post-hoc test showed that in somatic reproductive and somatic non-reproductive tissues, the direction of the dN/dS difference between the gametologs is significantly different between the X-biased and X’-biased groups; germline tissue shows the same trend, but is marginally non-significant (Fig. 3D-F). In general, where expression is X’-biased, the X’ gametolog also has more conserved molecular evolution relative to the X gametolog, and vice versa. In other words, when one of the gametologs has more biased expression, the other is under reduced purifying selection.

**Figure 3.**
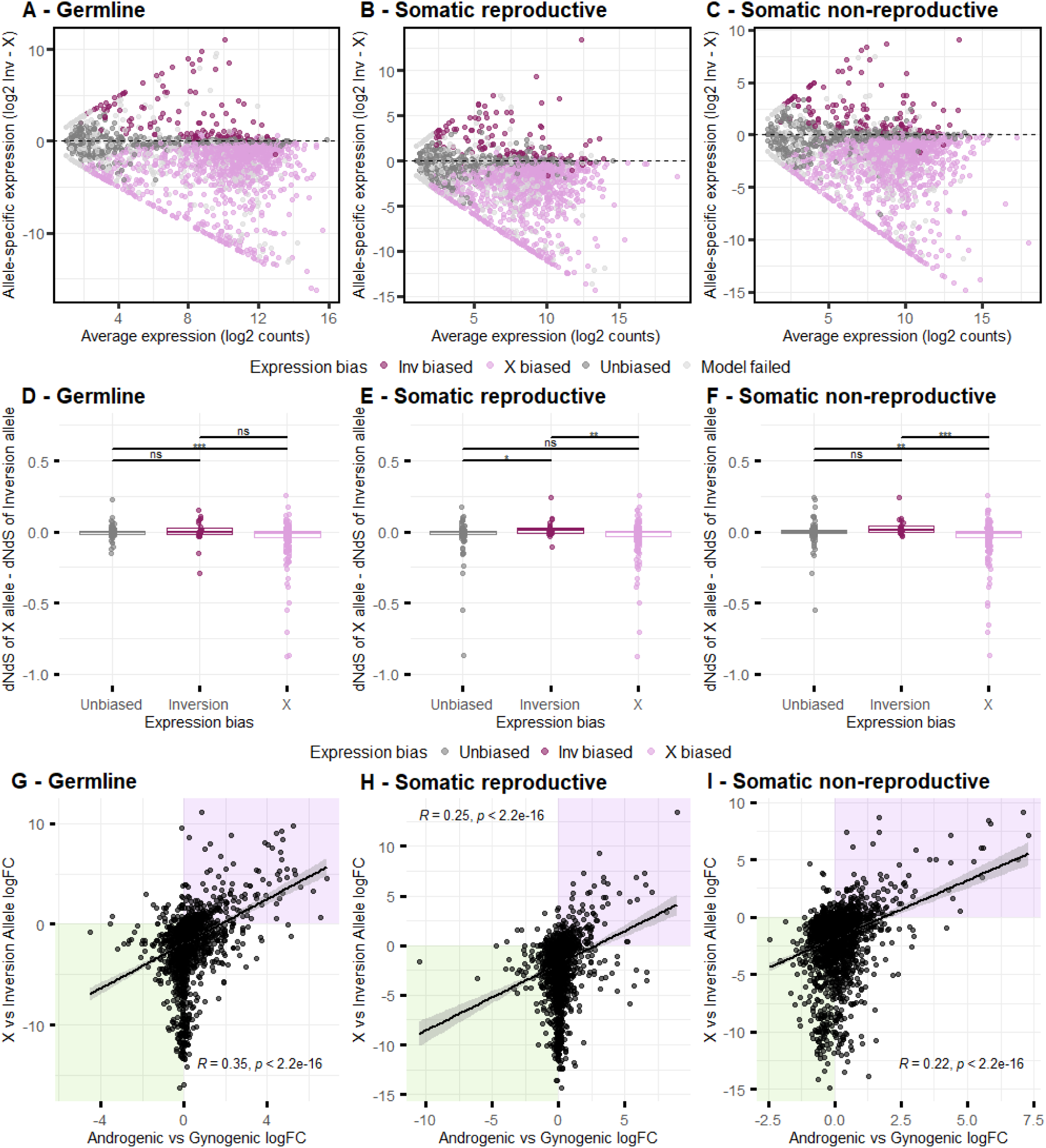
**(A-C)** MA plots looking at allele-biased expression between the X and X’ allele within gynogenic females, in germline, somatic reproductive tissue, and somatic non-reproductive tissue, respectively. The x-axes shows average expression of the gene in three replicates while the y-axes shows the log_2_ of the difference in expression between the two alleles. Values on the y-axes >0 shows X’-biased expression, while <0 shows X-biased expression. Genes with significant allele-biased expression are coloured in according to the direction of bias (maroon for X’, light pink for X). While most X-linked genes in gynogenic females have X-biased expression, there are a number of genes with X’-biased expression in all three tissues. **(D-F)** The difference in ratio of non-synonymous to synonymous substitutions between the X and X’ inversion gametologs, grouped by expression bias. This is presented separately for germline, somatic reproductive, and somatic non-reproductive tissue. Welch ANOVA and Games-Howell post-hoc test shows significant differences in X - X’ dN/dS in X-biased and X’-biased genes in all tissues. In somatic non-reproductive tissue, they are also both significantly different from unbiased genes. Genes with significant allele-biased expression are again coloured in according to the direction of bias (maroon for X’, light pink for X). For all plots, *** = p < 0.001, ** = p < 0.01, and * = p < 0.05. **(G-I)** Scatterplots showing the relationship between differential expression in genes between female reproductive morphs, and allele-biased expression between X and X’ gametologs in gynogenic females. The x-axes show log fold-change of a X-linked gene between androgenic and gynogenic females, with the former as the baseline. The y-axes show the logFC of the X and X’ gametolog of the same X-linked gene within gynogenic females, with X genes as the baseline. The purple-coloured region shows genes with both gynogenic-biased expression when comparing between female morphs, and X’ gametolog-biased expression within X’X gynogenic females. The green region shows genes with both androgenic-biased expression between female morphs, and X gametolog-biased expression within X’X gynogenic females. The result of a Spearman correlation test is shown, as well as a line drawn with the “lm” method.

**Table 2.** Summary table of the number of X-linked genes with allele-biased expression in gynogenic females, in germline, somatic reproductive, and somatic non-reproductive tissue. Unexpressed genes and genes whose models failed to converge are not included.

| Tissue \ X-linked expression | X-biased | Unbiased | X'-biased |
| --- | --- | --- | --- |
| **Germline** | 1220 | 484 | 139 |
| **Somatic reproductive** | 1224 | 596 | 123 |
| **Somatic non-reproductive** | 1362 | 654 | 137 |

Detailed results for X’-biased genes, with functional information, are found in Supplementary Material 3.

To relate this back to morph-specific, X-linked differential expression, we carried out Spearman’s test on the correlation between morph-specific X-linked expression, and allele-biased expression between the X and X’ gametologs. We find a significant association—gynogenic-biased genes are more likely to have X’-biased expression within gynogenic females, and vice versa. This association holds across all tissues (germline: *rho* = 0.35, *p-value* < 2.2^-16^; somatic reproductive: *rho* = 0.25, *p-value* < 2.2^-16^; somatic non-reproductive: *rho* = 0.22, *p-value* < 2.2^-16^), partly driven by a cluster of gynogenic-biased, X’-biased genes (Fig. 3G-I).

X-linked divergence in molecular evolution or expression could either directly encode morph-specific phenotypes, or be trans-acting regulators of autosomal gene expression in a morph-specific manner. We tested the latter by investigating autosomal differential expression between the female morphs.

#### 2.2.3 Significant gene expression divergence in autosomal genes

Since maternal sex-determining genes are X-linked, our null expectation was for little to no DGE in autosomal genes; any autosomal differences would reflect trans-regulatory effects from X/X′ and could underlie the phenotypic divergence reported in Results 2.1.

Although autosomes are found indiscriminately in both morphs, we find that gynogenic and androgenic females have distinct autosomal transcriptomic profiles (Fig. 4A-C), with a significant number of DEGs in germline and somatic non-reproductive tissue (Table 3, Fig. 4D-F). Strikingly, somatic non-reproductive tissue has even more DEGs than germline tissue, despite being unlikely to mediate offspring sex determination. As with X-linked genes, we also observe a bias towards overexpression in gynogenic females even in non-DEGs across all three tissues.

**Figure 4.**
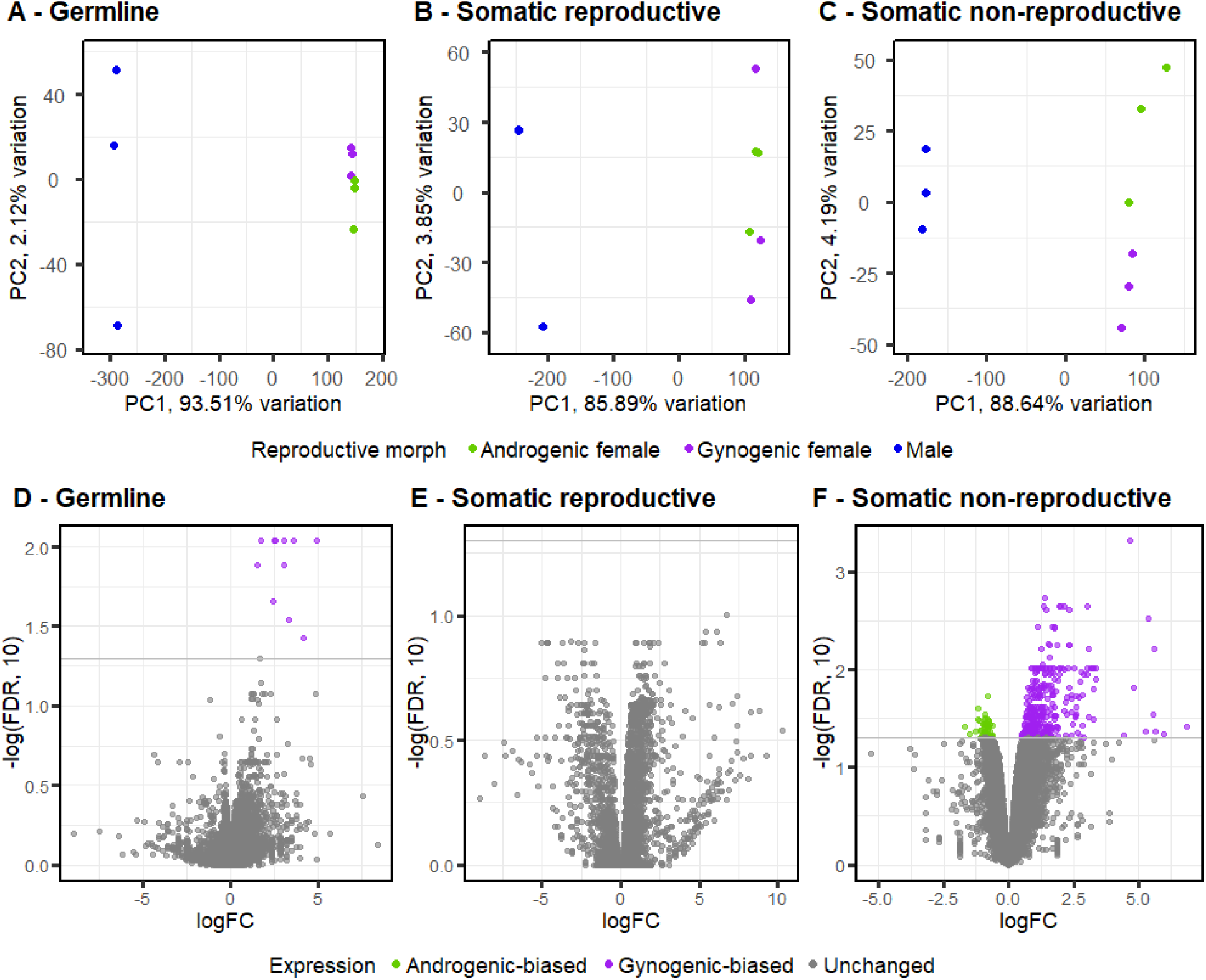
**(A-C)** Principal component analysis of autosomal gene expression in germline, somatic reproductive tissue, and somatic non-reproductive tissue, respectively. Samples show clustering first by male (blue) vs female, with subsequent clustering by female reproductive morph (purple for gynogenic, green for androgenic). **(D-F)** Volcano plots demonstrating differential gene expression in germline, reproductive tissue, and somatic non-reproductive tissue, respectively. The x-axis is the log fold-change of gene expression between gynogenic and androgenic females, using androgenic females as a baseline. Values >0 indicate gynogenic-biased expression, while values <0 indicate androgenic-biased expression. The y-axis shows the -log_10_ of the false discovery rate (FDR), a p-value adjusted for multiple testing. The grey horizontal line is drawn at -log_10_(0.05), and genes above that line are coloured (purple for gynogenic and green for androgenic) to signify significance.

**Table 3.** Summary table of the number of autosomal genes with significant differential expression between androgenic and gynogenic females. This is shown separately for germline, somatic reproductive, and somatic non-reproductive tissue.

| Tissue \ Autosomal expression | Androgenic-biased | Unbiased | Gynogenic-biased |
| --- | --- | --- | --- |
| Germline | 0 | 10481 | 11 |
| Somatic reproductive | 0 | 10816 | 0 |
| Somatic non-reproductive | 58 | 11263 | 335 |

In an attempt to shed light on the functional impact of this, we carried out a gene ontology enrichment analysis using topGO (Alexa and Rahnenführer, 2025), finding significant enrichment in broad categories such as transmembrane transport. Detailed results for DEGs with functional information are found in Supplementary Material 4.

### 2.3 Mechanisms of specialisation

We have shown evidence for divergence in life history traits and gene expression between reproductive morphs, with patterns of allele-specific expression and molecular evolution consistent with adaptive divergence between the two female morphs. To determine the driver of this specialisation, we tested two hypotheses—feminisation of gynogenic female gene expression as a result of release from SAS, and sex-specific provisioning of eggs.

#### 2.3.1 No evidence for release from sexually antagonistic selection

While the X chromosome is found in both males and females, the X’ is exclusively present in gynogenic females. Therefore, while the X may be constrained by differing male and female optimum, the X’ has no such restrictions and may evolve more extreme female-biased expression. To test for “feminisation” in gynogenic gene expression, we fitted a linear model comparing the average expression of each gene for gynogenic females vs male, and androgenic female vs male. While we expected a greater difference between gynogenic female and male gene expression than between androgenic female and male gene expression, we find no significant difference between the contrasts (Fig. 5). This was the case while analysing all genes, only autosomal genes, or only X/X’ genes. Thus, gynogenic females do not exhibit more extreme female gene expression than androgenic females.

**Figure 5.**
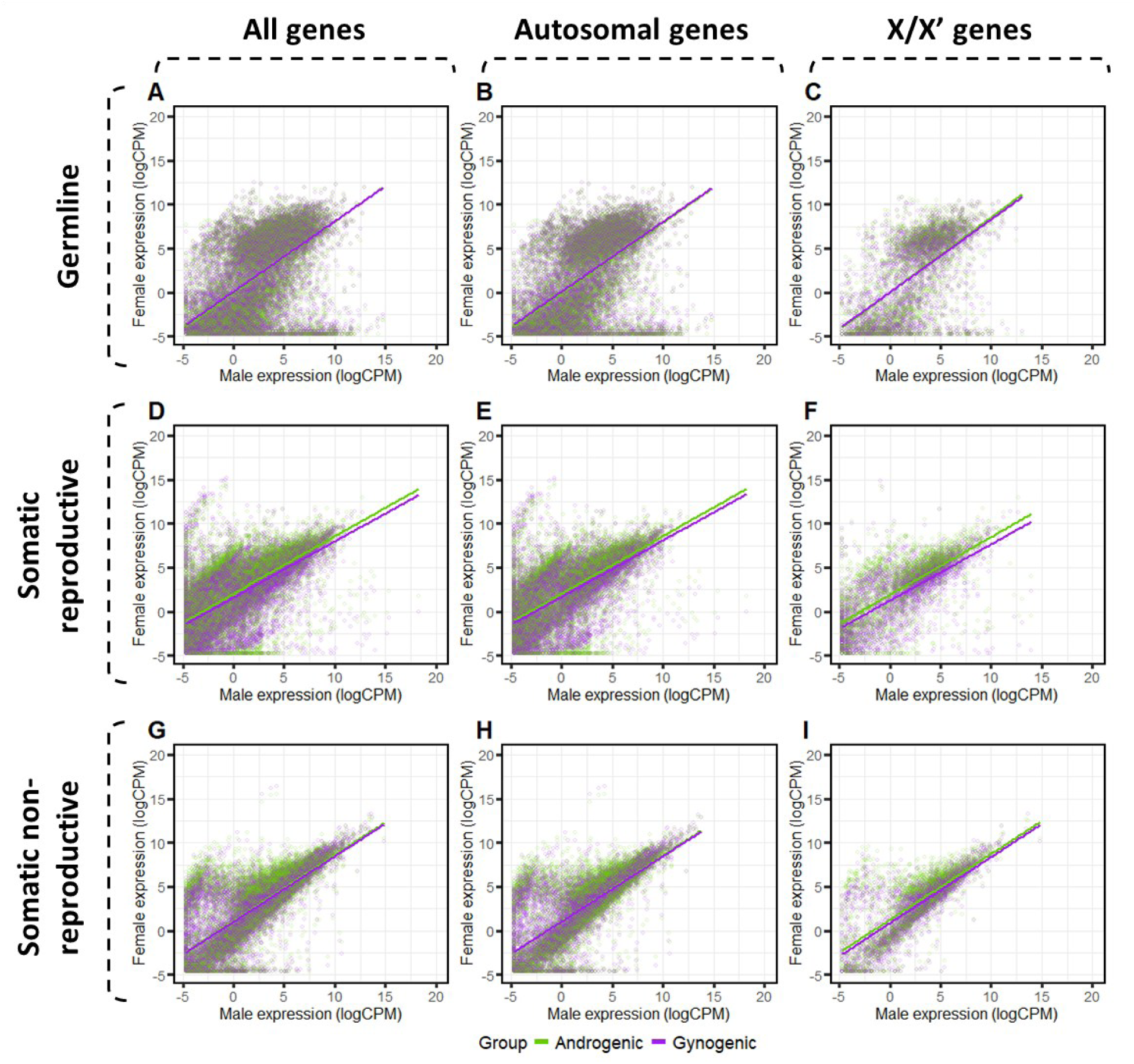
Plots for average male expression against average female expression for all genes **(A, D, G)**, autosomal genes only **(B, E, H)**, and X/X’ genes only **(C, F, I)**; for germline **(A-C)**, somatic reproductive tissue **(D-F)**, and somatic non-reproductive tissue **(G-I)**. Female expression is further divided into androgenic (green) and gynogenic (purple) expression. Regression lines are drawn between male and female expressions separately for the two female morphs. There is no significant difference in the regression lines of the two morphs in any group of genes in any tissue, suggesting that there is no difference in relationship between how expression in the two reproductive morphs compare to male expression.

#### 2.3.2 Reproductive morphs exhibit sex-specific offspring provisioning

In zygotic GSD systems, sex is only determined at fertilisation, leaving little opportunity for offspring to receive sex-specific care in species with minimal post-fertilisation parental care. However, in *mat*-GSD systems, mothers determine offspring sex pre-fertilisation and can provision their eggs accordingly, for example by mRNA deposition. We compared maternally deposited transcripts in female embryos produced by gynogenic females versus male embryos produced by androgenic females.

We find significant sex-biased deposition of X-linked mRNAs (Fig. 6A-B, Table 4). In female embryos, deposition from X vs X’ gametologs varied; although most are X gametolog-biased, there are still a significant number of transcripts that are X’-biased, mirroring results from maternal tissue (Fig 6C, Table 4). Female-biased maternal transcripts are also correlated with an X’-biased expression within female embryos (Fig. 6D). We further find numerous differentially deposited autosomal transcripts (Fig. 6E-F, Table 4). While some of these genes may be involved in maternal sex determination, many—especially autosomal genes—likely reflect broader sex-specific provisioning. Together, these results support sex-specific maternal provisioning as a driver of morph divergence. Further functional information for differentially expressed X’-biased genes and autosomal genes can be found in Supplementary Material 5.

**Figure 6.**
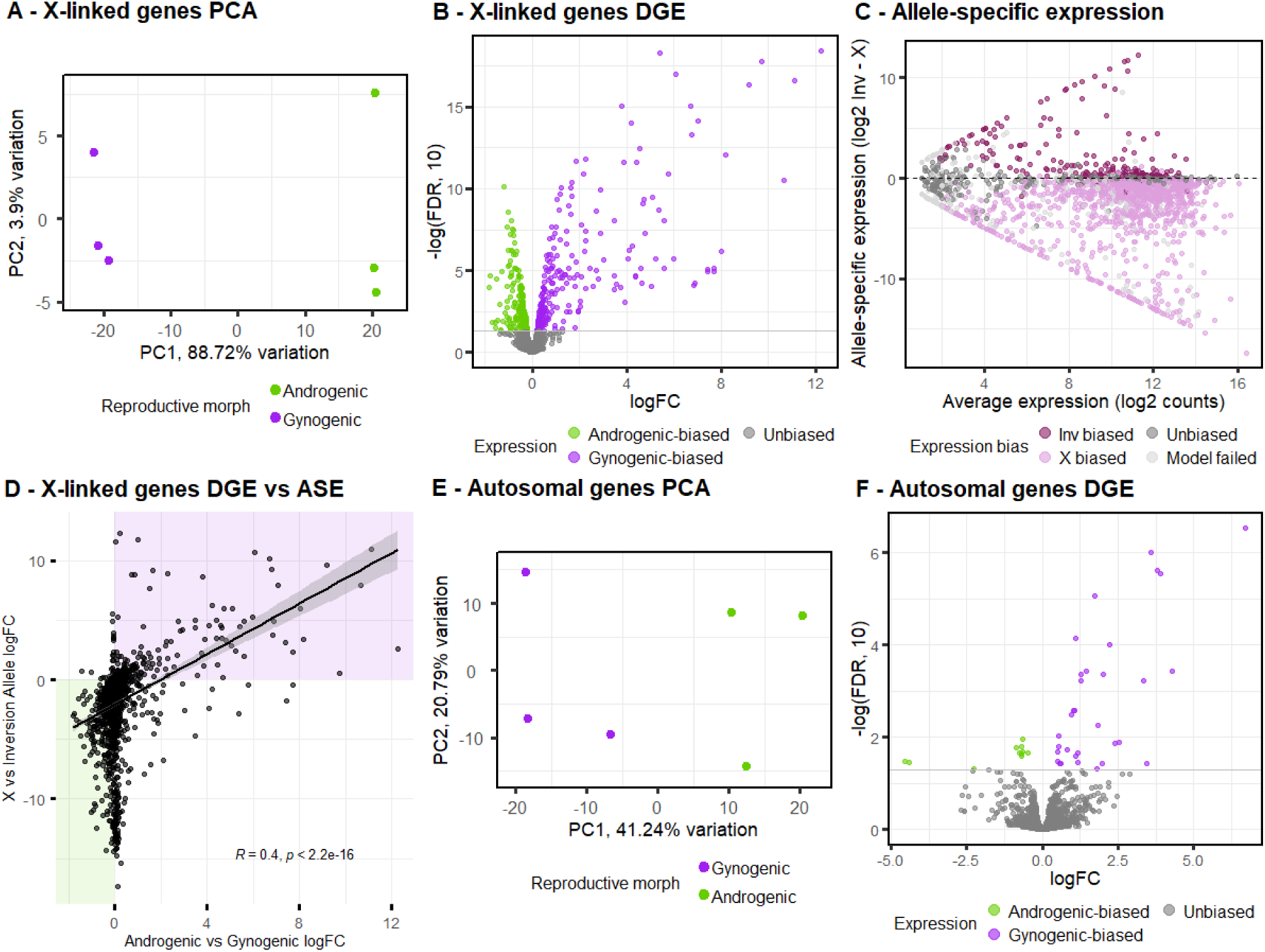
**(A, E)** Principal component analysis of maternal RNA deposited into eggs, for X-linked (X and X’) genes (A) and autosomal genes (E). In both cases, samples show clustering by female reproductive morph. **(B, F)** Volcano plot of maternal RNA deposits, for X-linked genes (B) and autosomal genes (F). The x-axis is the log fold-change of gene expression between gynogenic and androgenic females, using androgenic females as a baseline. Values >0 indicate gynogenic-biased expression, while values <0 indicate androgenic-biased expression. The y-axis shows the -log_10_ of the false discovery rate (FDR), a p-value adjusted for multiple testing. The grey horizontal line is drawn at - log_10_(0.05), and genes above that line are coloured (purple for gynogenic and green for androgenic) to signify significance. **(C)** A MA plot showing allele-biased expression between the X and X’ allele in gynogenic female maternal deposits, using data from eggs pre-determined to develop as females. The x-axes shows average expression of the gene in three replicates while the y-axes shows the log_2_ of the difference in expression between the two alleles. Values on the y-axes >0 shows X’-biased expression, while <0 shows X-biased expression. Genes with significant allele-biased expression are coloured in according to the direction of bias (purple for X’, green for X). **(D)** Scatterplot showing the correlation relationship between differential expression in genes between female reproductive morphs, and allele-biased expression between X and X’ gametologs in gynogenic females, in maternal RNA deposit.

**Table 4.** Summary table of differential gene expression in maternal RNA deposits into eggs, for X-linked and autosomal genes. Also allele-biased expression for X and X’ gametologs within the maternal deposits of gynogenic females.

|  | Androgenic-biased | Unbiased | Gynogenic-biased |
| --- | --- | --- | --- |
| **X-linked maternal deposit** | 208 | 1323 | 230 |
| **Autosomal maternal deposit** | 11 | 7409 | 32 |
|  | **X-biased** | **Unbiased** | **X'-biased** |
| **X'X gametologs** | 1152 | 363 | 194 |

## 3. Discussion

We have shown that in a monogenic species of fungus gnat, female-producing (gynogenic) females and male-producing (androgenic) females diverge phenotypically and transcriptionally in traits/tissues/genes other than offspring sex determination. Phenotypically, gynogenic females are not less fit than androgenic females, despite being the carrier of the non-recombining X’ region. This suggests that it is unlikely for degradation of the X’ to be the only force driving reproductive morph divergence. Supporting this, allele-specific expression analysis between the X and X’ in gynogenic females revealed a significant number of genes with X’-biased expression, which are also those with more conserved X’ gametologs relative to the X gametolog, suggesting that selection is maintaining the expression and function of a non-negligible number of genes on the inversion. The same genes are more likely to be upregulated in gynogenic females, potentially contributing to between-morph specialisation. While some of these genes may be involved in offspring sex determination, it is likely to be only a minority.

One of the more surprising findings is the significant number of autosomal DEGs between the female morphs, which are unlikely to be involved in offspring sex determination, especially in somatic non-reproductive tissues, but are in line with the significant divergence in life history traits observed in the phenotypic assay. DGE in both sex chromosomes and autosomes mirrors what is observed in species with dimorphic sexes (Ellegren and Parsch, 2007), supporting the hypothesis that the female morphs are specialising into their reproductive roles. The autosomal DEGs were not enriched for specific GO terms, suggesting a multi-trait specialisation also typical in sexual dimorphism (Parker et al., 1972; Mank, 2009; Barrett and Hough, 2013). Future work using organ-specific or cell-specific RNA-seq, as well as more lineage-specific functional annotations, could further help clarify the traits involved in morph specialisation. This provides the first formal example of mating type specialisation at the maternal level, extending our existing understanding of reproductive specialisation.

However, the mechanism driving their divergence remains to be elucidated. We will first elaborate on the two hypotheses tested in this study—release from SAS and sex-specific maternal provisioning—before suggesting a third possible mechanism. Although theory suggests that the X’ being released from SAS would result in feminisation of the gynogenic female transcriptome (Rice, 1984; Wright and Mank, 2013), we did not find evidence for this. It is possible that specific genes are indeed evolving under this dynamic, but were not detectable using our methods. However, recent studies of transitions to asexual, female-only reproduction shows gene expression may change only in specific tissues, or can even be masculinised (Veltsos et al., 2017; Parker et al., 2019), suggesting that the relationship between sex-biased expression and sexual antagonism may not be so straightforward. Yet another factor may be at play in fungus gnats: paternal genome elimination (PGE). Under PGE, males eliminate their paternal genome and pass on only their maternal genome. Models suggest that female-beneficial mutations can invade the genome more easily than male-beneficial ones in this context, resulting in an overall more feminised genome (Hitchcock et al., 2022). There may therefore be little scope for further feminisation in gene expression for gynogenic females.

On the other hand, we find evidence for sex-specific maternal provisioning; instead of a handful of offspring sex determination genes, many maternal transcripts are differentially deposited into male vs female embryos. Mothers are known to deposit a range of molecules into their eggs, including components of the sex determination cascade (Cronmiller et al., 1988; Dübendorfer and Hediger, 1998; Verhulst et al., 2010). Under *mat*-GSD, mothers can provision offspring according to their future sex even without post-fertilisation parental care; selection for specialising into producing one sex over the other could drive the evolution of monogeny from digeny. Offspring often benefit in sex-specific ways from maternal condition, egg deposits, and host resources (Charnov et al., 1981; Clutton-Brock et al., 1984; Von Engelhardt et al., 2006). In fact, examples exist for maternal control of sex determination with differential offspring provisioning: increased yolk allocation feminises offspring in the scincid lizard *Bassiana duperreyi* (Radder et al., 2009), sex-specific egg sizes in a monogenic species of parasitic barnacle *Peltogasterella gracilis* (Kajimoto et al., 2024), and sex-specific provisioning in *Encarsia* autoparasitoid wasps (Hunter et al., 1993; Hunter and Godfray, 1995). In fungus gnats, rare offspring of the “wrong” sex can be produced, but are usually sterile (Davidheiser, 1943), consistent with receiving the mismatched sex-specific inputs during development. Quantifying trade-offs in sex-specific provisioning, e.g. comparing fitness of males from female-skewed digenic broods vs male-skewed broods, should clarify how sex-specific provisioning selects for female specialisation. While we show differential RNA deposit in eggs of androgenic vs gynogenic females, extending this to proteins and nutritional molecules will be informative.

A third mechanism—morph-specific mating behaviour—remains untested. Under PGE, female morphs are expected to have divergent mating strategies. Because males receive no fitness returns from sons (paternal genome is eliminated from sperm), selection strongly favours avoiding mating with androgenic females (Bull, 1983). Correspondingly, androgenic females should be selected to hide their identity to secure matings—potentially constraining morph divergence. Secondly, mothers should prefer a high quality male for her daughters but not necessarily for her sons, who will not transmit her mate’s genes (although paternal genes are expressed in the soma). Behavioural experiments in fungus gnats demonstrate that males do not mate preferentially with gynogenic females, partly due to increased choosiness in gynogenic females and higher receptivity in androgenic females (Featherston et al., 2013; Hodson et al., 2024). Females with specialised mating strategy according to offspring sex may do better than those with mixed-sex broods. As a sensory/behavioural adaptation, its transcriptomic basis is likely somatic, consistent with our observations. Comparisons between the fitness of daughters and sons sired by low quality vs high quality male, would clarify the benefits of morph-specific mating behaviour. Importantly, the three mechanisms described above need not be mutually exclusive.

It is clear that PGE profoundly shapes predictions for fungus gnat evolution (Hitchcock et al., 2024). Intriguingly, PGE is also present in Cecidomyiids (gall midges), another group of insects with monogenic reproduction (Ross et al., 2022). The two groups share unusual features in sex determination and inheritance; maternal sex determination via X elimination (Stuart and Hatchett, 1991; Gerbi, 2022), a gynogenic-specific supergene (Benatti et al., 2010), and germline-restricted chromosomes (GRCs)—yet whether female morphs diverge in gall midges is completely unknown. Comparative work between these clades will enable stronger inferences about how these unusual reproductive strategies co-evolve.

Alternatively, monogeny without PGE also occurs (e.g. blowflies (Scott et al., 2014; Andere et al., 2020), parasitic barnacles *Peltogasterella gracilis* (Kajimoto et al., 2024)); testing for female-morphs divergence in those clades will clarify conditions favouring the evolution of mating type specialisation.

The specialisation of female morphs into distinct reproductive roles likely shapes the evolution of monogeny. Drawing on models linking sexually antagonistic selection to sex chromosome evolution (Lesaffre et al., 2024; Flintham and Mullon, 2026), we propose that morph-specific disruptive selection can drive a shift from digeny to monogeny, facilitated by/driving an underlying supergene that links morph-beneficial alleles. However, progressive accumulation of deleterious mutations on the non-recombining supergene results in decreased fitness in gynogenic females and reversion to digeny. A cycle of specialisation, associated with an underlying genetic structure which then degenerates, can explain the high rates of turnover of monogeny and digeny (Box 2). Phenotypic divergence and fitness patterns suggest our study lines are at Step 1 of this cycle. As this model predicts that gynogenic females bearing an older supergene should be less fit than androgenic females, further sampling for inversions of different ages and the associated morph fitnesses would inform the relevance of this model for mating system turnover in fungus gnats.

### Box 2: Hypothesis for turnover of monogeny and digeny in fungus gnats

In this study, we have shown evidence for morph-specific divergence and specialisation in the monogenic *B. coprophila*. This informs a model for the extensive turnover of monogeny and digeny across the tree of fungus gnats.

Step 1. We start with a digenic ancestor (yellow) that produces offspring of both sex, where the sex ratio is a polygenic trait. Maternal traits for producing sons or producing daughters may have divergent optima, causing disruptive selection for biased sex ratio in association with the beneficial maternal trait. This selection pressure results in female morph specialisation and split sex ratio, producing gynogenic females (in purple) and androgenic females (in green). Concurrently, there is also selection for linkage between alleles for extreme offspring sex ratios and the maternal trait beneficial for that sex. The result is a maternally-acting supergene (chromosomal region in purple) that determines offspring sex as well as a suite of other maternal traits.

Step 2. Although recombination suppression allows beneficial traits to be inherited together, it also reduces the efficacy of selection against deleterious mutations. Additionally, the maternal supergene has one-sixth the N_e_ of an autosome, driving deleterious mutation accumulation at a faster rate than the recombining X chromosome (red sites on the gynogenic-specific region). Although initially the deleterious effects may be compensated for by the beneficial impacts of specialisation, in the long term the mutation load borne by gynogenic females is likely to outweigh the benefits. Subsequently, the fitness of gynogenic females decreases relative to that of androgenic females.

Step 3. Finally, as the relative fitness of these two female morphs directly correspond to population sex ratio, unequal fitness of the two morphs may result in local population extinction or reversion back to digeny. We end up at the start of the cycle.

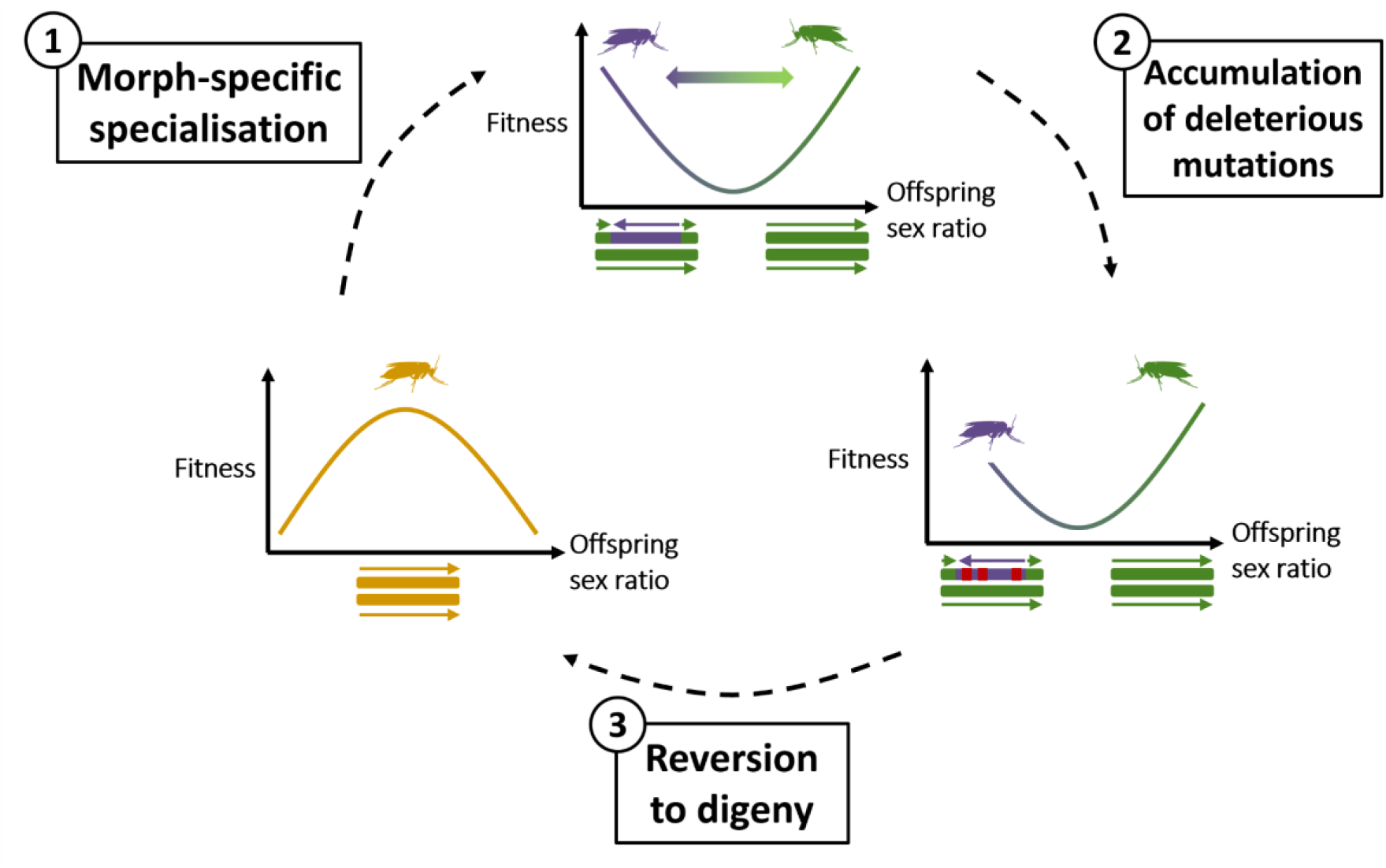

We have characterised and explored the evolutionary driver of reproductive morph divergence in a monogenic species of fungus gnat, associated with a large and recently evolved supergene. Our findings parallel empirical and theoretical work on sexual specialisation and its role in sex chromosome evolution. With the drivers of recombination suppression in sex chromosome evolution still under hot debate, (Charlesworth, 2023; Jay et al., 2024), morph-determining regions in fungus gnats and gall midges offer an analogue for testing predictions. For example, mapping the *mat*-GSD locus and morph specialisation loci, and whether linkage between the two is selected for, can help support the SAS hypothesis. Reproductive morph specialisation is only one part in the intriguing system of fungus gnats; their atypical sex-determining and mating systems provide valuable tests of general principles and conventional assumptions in evolutionary biology.

## 4. Methods

### 4.1 Phenotypic divergence

The two distinct lines of *Bradysia coprophila* used in the assay—the Holo2 (H2) line and the KM line—required different methods of morph-determination. The H2 line has a dominant wavy wing marker on the X’ (Fig. 7A), allowed morphs to be determined at adulthood. In the KM line, polymerase chain reaction (PCR) primers were developed flanking a site where a deletion is present on the X’ but not the X (DEL3018: Fr: 5’-TGCCAGACAGTTGCAGATTG-3’; Rv: 5’-CTCGTTCTCCACTGGTGTTG-3’), yielding two DNA fragments of different lengths in gynogenic females, and only one in androgenic females (Fig. 7B). Husbandry for all lines followed (Gerbi, 2024).

**Figure 7.**
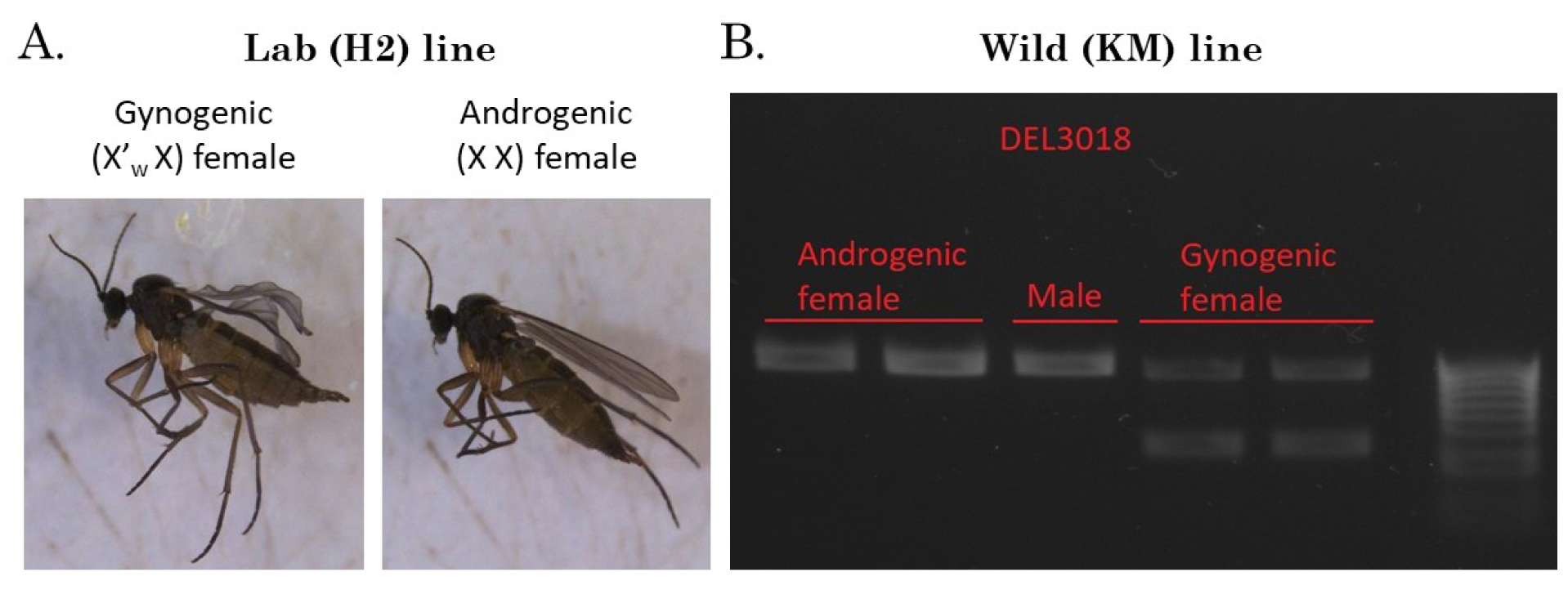
Methods to differentiate between female reproductive morph in the lab and wild line. A.) Holo2 wing markers in a lab stock of *B. coprophila*. Gynogenic (X’X) females carry a dominant wavy marker (W) on the X’ chromosome, which is not present in androgenic (XX) females. Males are phenotypically different from females, most prominently by the presence of a pair of posterior claspers (not pictured). B.) Molecular assay using primer DEL00003018 to differentiate between gynogenic and androgenic females in a wild line with no wing markers. The primer selectively amplifies a region of the X chromosome where the homologous region on the X’ houses a ∼500bp deletion. Therefore, one long fragment is amplified from XX females, while two DNA fragments of different sizes are amplified from X’X females.

The experiment was conducted across three generations, to measure traits in the focal F_1_. Measurements were taken in each F_1_ female for developmental time (days from first egg/larval hatching to eclosion as adults), differential survival (number of adult gynogenic vs androgenic female per clutch), body size (thoracic length), and adult lifespan. A random subset of females were selected and mated to measure fecundity (number of adult progeny). Date-related measurements are accurate to +/-1 day. The outbred line was generated by crossing F0 H2 females with KM males to produce F1 H2xKM individuals.

Otherwise, life history measurements were carried out in the same way. Detail methodology for each trait is found in Supplementary Material 1.

Statistical analyses were run per trait after assessing normality and homogeneity of variance using the DHARMa package in R (Hartig et al., 2024). Developmental time, adult size and adult lifespan were fitted with linear mixed-effect models using the lme4 package in R (Bates et al., 2015), including fixed effects of female morph (androgenic/gynogenic), line (H2/KM), and an interaction between morph and line, with family as a random effect. Adult lifespan was further fitted with a fixed term for mated/unmated. Differential survival was fitted as a binomial GLM with a logit link, testing whether the log-odds of survival between gynogenic and androgenic females deviated from 0. To account for overdispersion and excess zeros from failed matings, the number of adult offspring (fecundity) was fitted with a zero-inflated negative binomial mixed-effects model using the glmmTMB package in R (Brooks et al., 2025), with the same fixed effects as above plus female mating age and thoracic length. For the outbred lines, models were the same except for the omission of line and line:morph interaction terms, since there was only one line. Full models and results can be found in Supplementary material 2.

### 4.2 Gene expression divergence

Gene expression analysis was carried out using only the H2 line. The RNA-seq libraries were produced for (Baird, 2024) and are available on Genbank under accession PRJNA1109384. The reference genome is available under accession PRJNA953429.

#### 4.2.1 Differential gene expression analysis - X-linked and autosomal genes

We analysed only X-linked genes with recognisable, single-copy gametologs between the X and X’ inversion, identified with Orthofinder (Emms and Kelly, 2019). To obtain read counts for each gametolog, we filtered and trimmed the raw RNA-seq reads with FastQC (Andrews, 2010) and fastp (Chen et al., 2018), then mapped the reads to a reference genome containing separate X and X’ scaffolds with STAR (Dobin et al., 2013). Read counts were summarised to the gene level with featureCounts (Liao et al., 2014), counting multimappers fractionally. This means that reads mapping equally well to both gametologs would multimap in both gynogenic and androgenic females (despite certain origin from X in the latter). This is dealt with in this section by summing the read counts across X/X’ gametologs to produce a single per-pair count, which allows for comparison between the female morphs. As a check, mapping androgenic reads to references with vs without the inversion region changed overall mapping rates by <2%, indicating limited mis-mapping.

DGE analysis on the per-pair counts was carried out with edgeR (Chen et al., 2025), using androgenic females as the arbitrary baseline. Genes with positive logFC are therefore gynogenic-biased, and vice versa. The threshold for significance was set at false discovery rate (FDR) < 0.05. PCA plots were generated from logCPM values, with a prior count of 1.

DGE in autosomal genes was analysed with the same process (Results 2.2.3), subsetting read counts for autosomal genes instead before analysis in edgeR. Furthermore, gene ontology enrichment analysis on DEGs were carried out using the topGO package in R (Alexa and Rahnenführer, 2025).

#### 4.2.2 Allele-biased expression analysis

Allele-biased expression was quantified for RNA-seq libraries from gynogenic females, separately for germline, somatic reproductive, and somatic non-reproductive tissue. The process of read trimming and mapping was the same as 4.2.1, but with a higher quality threshold to reduce the proportion of reads with potential sequencing errors. The reads were then mapped to the *B. coprophila* reference genome (with X’ scaffold) using STAR. Alignments with mapping quality < 10 were removed with SAMtools (Danecek et al., 2021), retaining unique mappers as well as primary alignments, but not perfect multimappers and secondary alignments. Since reads are mapped competitively, this means that only reads with informative SNPs are kept, which should be mapped to their gametologs of origin. Read counts were summarised to gene level with featureCounts.

Genes with no reads in any of the three replicates were removed. Genes with complete gametolog-limited expression (>= 10 reads in one gametolog and none in the other) were analysed separately. For pairs with expression in both gametologs, allele-specific expression bias was assessed using a beta-binomial generalised linear model fitted independently per pair, using the glmmTMB package in R. As X’X gametologs may have different gene lengths, due to degradation or lower quality X’ assembly, a fixed offset term was added to the model, which describes the null expectation of read distribution between the two gametologs proportional to gene lengths. This permits testing for differences between the two gametologs beyond what can be explained by length differences. P-values were corrected for multiple testing using the Benjamini-Hochberg method, where adjusted p < 0.05 indicated significant biased expression.

#### 4.2.3 Rate of molecular evolution (dN/dS analysis)

Using parts of the pipeline in (Baird et al., 2025), the ratio of nonsynonymous substitution to synonymous substitution was calculated and compared between X’X gametologs. For each gametolog, synonymous and nonsynonymous site counts were obtained with a custom script *partitions.py* (Mackintosh et al., 2022). Subsequently, we counted the number of synonymous and nonsynonymous mutations by mapping the *B. coprophila* genome to a digenic outgroup, *B. odoriphaga* (Accession: SRR11366020), which lacks a X’ region, with minimap2 (Li, 2018). Variants were called with BCFtools (Danecek et al., 2021), and annotated and filtered with SnpEff and SnpSift (Cingolani et al., 2012a, 2012b). dN/dS was computed and compared for each gametolog pair in R.

### 4.3 Release from sexually antagonistic selection

To test whether gynogenic adaptation reflects release from SAS on the X′, we compared gene expression between gynogenic females, androgenic females, and males. We predicted that gynogenic females would evolve more extreme female-biased gene expression; in other words, the gynogenic–male contrast should diverge further from the 1:1 line than the androgenic–male contrast. We quantified this by regressing female morph expression on male expression and testing the slope differences with an interaction term.

To allow comparison across both autosomal and X-linked genes, RNA-seq reads were mapped to a N-masked version of the reference genome. Using parts of the pipeline in (Baird, 2024), we generated the N-masked genome by 1.) subsetting for the autosomes and X scaffolds, and 2.) N-masking the sites on the X scaffold that is a variant between the X and X’. Sites to be N-masked were determined by mapping a DNA library of a gynogenic female to the subsetted reference genome with BWA (Li and Durbin, 2009). After assigning read groups and marking duplicates with Picard (http://broadinstitute.github.io/picard), variants in the alignment were called and filtered for quality using BCFtools. Finally, variant sites on the X chromosome were N-masked using BEDTools (Quinlan and Hall, 2010).

Trimmed RNA-seq reads were mapped to the N-masked genome using STAR, and summarised to gene level with featureCounts. Genes with low read counts were filtered, and logCPM values were extracted using edgeR. For each gene, we calculated mean expression by sex (gynogenic, androgenic and male), and fitted a lm (base R) to test for slope differences between gynogenic-male and androgenic-male relationships.

### 4.4 Sex-specific offspring provisioning

Under *mat*-GSD and monogeny, females can provision their eggs differently depending on whether they are pre-determined to develop as male or female. We tested one aspect of this by looking at maternally-deposited mRNA in male vs female eggs. Embryonic RNA-seq libraries were produced for (Baird, 2024) and are available under accession PRJNA1220056. Pooled broods of male or female embryos were collected 0-4 hours post-laying, prior to ZGA, to obtain only maternal transcripts (Phalle and Sullivan, 1996; Baird, 2024). We performed DGE for X-linked and autosomal transcripts between male and female embryos as in Methods 4.2.1, and tested allele-biased deposition (X vs X′) in female embryos of gynogenic mothers as in Methods 4.2.2.

All plots were generated in R using the ggplot2 and ggpubr packages (Kassambara, 2025; Wickham et al., 2025). Full walk-through code with package versions for each step of Methods can be found at https://github.com/RossLab/Reproductive_Morph_Specialisation

## Acknowledgements

The authors thank Katy Monteith for assistance with the phenotypic assay, and to members of the Ross Lab for helpful feedback and discussions. This work was funded by a European Research Council starting grant (PGErepro, to L.R.), and M.H. is funded by the NERC E4 DTP.

