## Supplementary files for "Reproductive morph specialisation facilitated by a maternal sex-determining region in a fungus gnat (*Bradysia coprophila*)"

### Supplementary materials

#### Supplementary material 1: Detailed methods on measurement of phenotypic traits

**Table S1.** Measurements of phenotypic traits. Methods of measurement differed slightly between H2 and KM lines because the H2 line carried a X'-linked wing marker that segregated differentially with female types, while the KM line did not. Therefore, in the H2 line female morph could be assigned at eclosion, while in the KM line female morph was assigned posthumously, using morph-specific primers.

| Phenotype | Method of measurement |
|---|---|
| Development rate | The date of egg laying and hatching for each F1 vial was noted.<br>The date of eclosion of each emerging individual was noted and individuals that emerged from the same family on the same day were placed in the same holding vial to await stress recovery trial. |
| Differential survival | The number of gynogenic and androgenic females were counted at adulthood. Deviations from 50-50 should indicate differential survival. |
| Adult lifespan | For each individual in vials with unique IDs, the date of death was noted down. Adult lifespan = Date of death - Date of eclosion. |
| Adult size | Dead individuals were frozen at -70°C. Later, pictures were taken under a microscope with a scale. Thoracic length (from tip/most forward part of the thorax to the most dorsal point where it connected with the abdomen) was measured using ImageJ (Fig. S1). |
| Fecundity | Healthy females between the ages of 3-6 days were chosen at random to be mated, to minimise the effects of extreme ages on fecundity. Progeny were allowed to develop until adulthood, and the number of progeny that survived until adulthood was counted for each mated $F_1$ mother. |

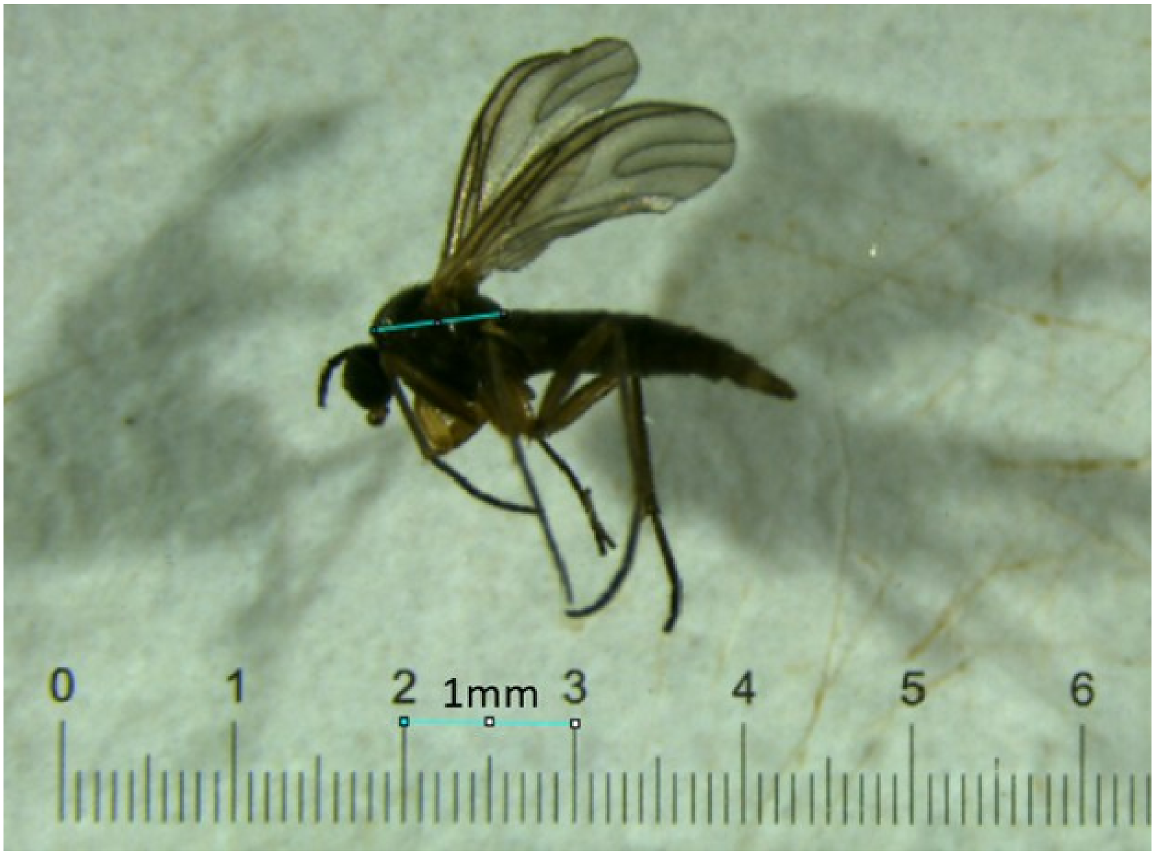


**Figure S1**. Thoracic measurement: the lower line shows a 1mm scale line drawn upon the calibration slide. The line across the thorax was drawn from the highest point at the top of the thorax to the point where the thorax and abdomen joined.

#### Supplementary material 2: Full models and results of phenotypic divergence

**<u>H2 and KM lines</u>**

1. Developmental rate (Days from first larvae hatching to eclosion as adult)

```
Linear mixed model fit by REML. t-tests use Satterthwaite's method
['lmerModLmerTest']
Formula: days_to_eclosion ~ f1_morph + line + f1_morph:line + (1 | family)
   Data: eclosion_all

REML criterion at convergence: 2972.6

Scaled residuals:
    Min      1Q  Median      3Q     Max
-4.5243 -0.5341 -0.0221  0.5225  3.4582

Random effects:
 Groups   Name        Variance Std.Dev.
 family   (Intercept) 30.11    5.487
 Residual             17.31    4.161
Number of obs: 512, groups:  family, 20

Fixed effects:
                 Estimate Std. Error       df t value Pr(>|t|)
(Intercept)       32.4906     1.8110  19.8023  17.941 1.03e-13 ***
f1_morphG         -2.2577     0.5648 491.4609  -3.997 7.39e-05 ***
lineKM            11.9731     2.5384  19.1459   4.717 0.000147 ***
f1_morphG:lineKM   5.4218     0.7578 491.2985   7.155 3.07e-12 ***
---
Signif. codes:  0 ‘***’ 0.001 ‘**’ 0.01 ‘*’ 0.05 ‘.’ 0.1 ‘ ’ 1

Correlation of Fixed Effects:
            (Intr) f1_mrG lineKM
f1_morphG   -0.173
lineKM      -0.713  0.124
f1_mrphG:KM  0.129 -0.745 -0.150
```

2. Differential survival (Number of adult gynogenic and androgenic females)

```
Generalized linear mixed model fit by maximum likelihood (Laplace
Approximation) ['glmerMod']
 Family: binomial  ( logit )
Formula: cbind(count_G, count_A) ~ -1 + Line + (1 | family)
   Data: survival_all
```

```
     AIC      BIC   logLik deviance df.resid
    94.4     97.4    -44.2     88.4       17

Scaled residuals:
    Min      1Q  Median      3Q     Max
-1.8631 -0.3161  0.1304  0.4403  1.5302

Random effects:
 Groups Name        Variance Std.Dev.
 family (Intercept) 0.03679  0.1918
Number of obs: 20, groups:  family, 20

Fixed effects:
       Estimate Std. Error z value Pr(>|z|)
LineH2   0.1474     0.1525   0.967    0.334
LineKM  -0.1992     0.1408  -1.415    0.157

Correlation of Fixed Effects:
       LineH2
LineKM -0.027
```

Notes for interpretation: In this generalised linear model with a binomial family, observing a gynogenic female is defined as "success" while an androgenic female is defined as a "failure". The intercept estimates the log-odds of "success" - i.e. observing a gynogenic female. For both H2 and KM lines, the intercept is not significantly different from 0, which means a log-odds of 1 - i.e. no difference in the odds of observing a gynogenic female vs an androgenic female.

3. Adult lifespan (Number of days from eclosion as adult to death)

Non-hurdle model:

```
Linear mixed model fit by REML. t-tests use Satterthwaite's method
['lmerModLmerTest']
Formula: adult_lifespan_days_mel ~ f1_morph + line + f1_morph:line + mated +
(1 | family)
   Data: lifespan_all

REML criterion at convergence: 2215.9

Scaled residuals:
    Min      1Q  Median      3Q     Max
-2.2656 -0.6358  0.0752  0.5848  4.7228

Random effects:
 Groups   Name        Variance Std.Dev.
 family   (Intercept) 0.5038   0.7098
```

```
 Residual              4.5770   2.1394
Number of obs: 503, groups:  family, 20

Fixed effects:
                   Estimate Std. Error        df t value Pr(>|t|)
(Intercept)         4.29944    0.33535  20.78947  12.821 2.45e-11 ***
f1_morphG           0.63619    0.28970 488.44261   2.196   0.0286 *
lineKM             -0.20450    0.43963  16.81475  -0.465   0.6478
mated1              1.39520    0.20703 489.91381   6.739 4.49e-11 ***
f1_morphG:lineKM  -0.01258    0.39266 485.41012  -0.032   0.9744
---
Signif. codes:  0 '***' 0.001 '**' 0.01 '*' 0.05 '.' 0.1 ' ' 1

Correlation of Fixed Effects:
            (Intr) f1_mrG lineKM mated1
f1_morphG   -0.468
lineKM      -0.717  0.362
mated1      -0.221 -0.023 -0.040
f1_mrphG:KM  0.326 -0.740 -0.444  0.103
```

Hurdle model: In the hurdle model, a binary variable is created to discern individuals with an early death (=< 1 day). A logistic regression is first carried out for the probability of having an early death. A linear model was then carried out for the subset of individuals who did not have an early death.

```
Generalized linear mixed model fit by maximum likelihood (Laplace
Approximation) ['glmerMod']
 Family: binomial  ( logit )
Formula: early_death ~ f1_morph + line + f1_morph:line + mated + (1 |
family)
   Data: lifespan_all

     AIC      BIC   logLik deviance df.resid
   317.2    342.5   -152.6    305.2      497

Scaled residuals:
    Min      1Q  Median      3Q     Max
-0.9392 -0.4291 -0.2956  0.0000  3.2040

Random effects:
 Groups Name        Variance Std.Dev.
 family (Intercept) 1.1      1.049
Number of obs: 503, groups:  family, 20

Fixed effects:
                 Estimate Std. Error z value Pr(>|z|)
(Intercept)       -1.0873     0.4396  -2.473   0.0134 *
f1_morphG         -0.3170     0.3824  -0.829   0.4071
lineH2            -0.1468     0.6712  -0.219   0.8268
```

```
mated1           -21.8836   139.3488  -0.157   0.8752
f1_morphG:lineH2  -0.1267     0.6293  -0.201   0.8404
---
Signif. codes:  0 ‘***’ 0.001 ‘**’ 0.01 ‘*’ 0.05 ‘.’ 0.1 ‘ ’ 1

Correlation of Fixed Effects:
            (Intr) f1_mrG lineH2 mated1
f1_morphG   -0.394
lineH2      -0.656  0.261
mated1      -0.001  0.000  0.001
f1_mrphG:H2  0.240 -0.609 -0.446  0.000

Linear mixed model fit by REML. t-tests use Satterthwaite's method
['lmerModLmerTest']
Formula: adult_lifespan_days_mel ~ f1_morph + line + f1_morph:line + mated +
(1 | family)
   Data: longer_lived

REML criterion at convergence: 1771.5

Scaled residuals:
    Min      1Q  Median      3Q     Max
-2.5846 -0.5901  0.0221  0.4927  5.1649

Random effects:
 Groups   Name        Variance Std.Dev.
 family   (Intercept) 0.1946   0.4412
 Residual             3.0989   1.7604
Number of obs: 442, groups:  family, 20

Fixed effects:
                  Estimate Std. Error        df t value Pr(>|t|)
(Intercept)        5.35931    0.25799  29.49698  20.774  < 2e-16 ***
f1_morphG          0.55053    0.25178 435.48996   2.186  0.02931 *
lineKM            -0.25957    0.32832  24.66007  -0.791  0.43671
mated1             0.50196    0.17781 436.99220   2.823  0.00498 **
f1_morphG:lineKM   0.01863    0.34513 434.04536   0.054  0.95698
---
Signif. codes:  0 ‘***’ 0.001 ‘**’ 0.01 ‘*’ 0.05 ‘.’ 0.1 ‘ ’ 1

Correlation of Fixed Effects:
            (Intr) f1_mrG lineKM mated1
f1_morphG   -0.534
lineKM      -0.707  0.424
mated1      -0.287 -0.016 -0.049
f1_mrphG:KM  0.362 -0.731 -0.524  0.107
```

4. Adult size (Thoracic length)

```
Linear mixed model fit by REML. t-tests use Satterthwaite's method
['lmerModLmerTest']
Formula: thorax_length_mel ~ f1_morph + line + f1_morph:line + (1 | family)
   Data: thoracic_all

REML criterion at convergence: -1471.6

Scaled residuals:
    Min      1Q  Median      3Q     Max
-3.3965 -0.5038  0.0229  0.6298  3.8634

Random effects:
 Groups   Name        Variance Std.Dev.
 family   (Intercept) 0.001164 0.03411
 Residual             0.001790 0.04231
Number of obs: 444, groups:  family, 20

Fixed effects:
                  Estimate Std. Error        df t value Pr(>|t|)
(Intercept)      7.908e-01  1.209e-02 2.061e+01  65.380  < 2e-16 ***
f1_morphG        4.245e-03  6.063e-03 4.247e+02   0.700 0.484232
lineKM           6.555e-02  1.679e-02 1.933e+01   3.904 0.000929 ***
f1_morphG:lineKM 1.145e-02  8.273e-03 4.246e+02   1.384 0.167033
---
Signif. codes:  0 '***' 0.001 '**' 0.01 '*' 0.05 '.' 0.1 ' ' 1

Correlation of Fixed Effects:
            (Intr) f1_mrG lineKM
f1_morphG   -0.281
lineKM      -0.720  0.202
f1_mrphG:KM  0.206 -0.733 -0.253
```

5. Fecundity (Number of adult progeny)

```
Family: nbinom2  ( log )
Formula:          progeny_count ~ f1_morph + line + f1_morph:line +
female_mating_age +      thorax_length_mel + (1 | family)
Zero inflation:                 ~1
Data: progeny_all

      AIC       BIC    logLik -2*log(L)  df.resid
   1098.5    1125.7    -540.3    1080.5       142

Random effects:

Conditional model:
 Groups Name        Variance  Std.Dev.
 family (Intercept) 2.291e-08 0.0001514
```

```
Number of obs: 151, groups:  family, 20

Dispersion parameter for nbinom2 family (): 1.46

Conditional model:
                    Estimate Std. Error z value Pr(>|z|)
(Intercept)          -1.1222     1.7029  -0.659 0.509909
f1_morphG             0.9957     0.2852   3.491 0.000481 ***
lineKM                0.8772     0.2880   3.045 0.002324 **
female_mating_age    -0.1673     0.1122  -1.491 0.135935
thorax_length_mel     5.3544     2.0930   2.558 0.010521 *
f1_morphG:lineKM     -1.2734     0.3587  -3.550 0.000386 ***
---
Signif. codes:  0 '***' 0.001 '**' 0.01 '*' 0.05 '.' 0.1 ' ' 1

Zero-inflation model:
            Estimate Std. Error z value Pr(>|z|)
(Intercept)  -0.8715     0.1879  -4.638 3.52e-06 ***
---
Signif. codes:  0 '***' 0.001 '**' 0.01 '*' 0.05 '.' 0.1 ' ' 1
```

**H2xKM line**

1. Developmental rate (Number of days from eggs laid to eclosion as adult)

```
Linear mixed model fit by REML. t-tests use Satterthwaite's method
['lmerModLmerTest']
Formula: days_to_eclosion ~ f1_morph + (1 | family)
   Data: eclosion_hyb

REML criterion at convergence: 209.7

Scaled residuals:
    Min      1Q  Median      3Q     Max
-1.1403 -0.7603 -0.1932  0.7071  2.1761

Random effects:
 Groups   Name        Variance Std.Dev.
 family   (Intercept) 1.908    1.381
 Residual             4.459    2.112
Number of obs: 47, groups:  family, 9

Fixed effects:
            Estimate Std. Error      df t value Pr(>|t|)
(Intercept)  39.0883     0.6526  9.1288  59.896 3.66e-13 ***
f1_morphG    -1.8862     0.6435 39.3769  -2.931  0.00559 **
---
Signif. codes:  0 '***' 0.001 '**' 0.01 '*' 0.05 '.' 0.1 ' ' 1
```

```
Correlation of Fixed Effects:
          (Intr)
f1_morphG -0.457
```

2. Differential survival (Number of adult gynogenic and androgenic females)

```
Call:
glm(formula = cbind(count_G, count_A) ~ 1, family = binomial(link = "logit"), 
    data = survival_hyb)

Coefficients:
            Estimate Std. Error z value Pr(>|z|)
(Intercept)  -0.1278     0.2923  -0.437    0.662

(Dispersion parameter for binomial family taken to be 1)

    Null deviance: 6.6653  on 8  degrees of freedom
Residual deviance: 6.6653  on 8  degrees of freedom
AIC: 24.987

Number of Fisher Scoring iterations: 3
```

Notes for interpretation: In this generalised linear model with a binomial family, observing a gynogenic female is defined as “success” while an androgenic female is defined as a “failure”. The intercept estimates the log-odds of “success” - i.e. observing a gynogenic female. The intercept is not significantly different from 0, which means a log-odds of 1 - i.e. no difference in the odds of observing a gynogenic female vs an androgenic female.

3. Adult size (Thoracic length)

```
Linear mixed model fit by REML. t-tests use Satterthwaite's method
['lmerModLmerTest']
Formula: thorax_length ~ f1_morph + (1 | family)
   Data: thoracic_hyb

REML criterion at convergence: -170.7

Scaled residuals: 
    Min      1Q  Median      3Q     Max 
-2.6197 -0.6578  0.1514  0.6166  2.4387 

Random effects:
 Groups   Name        Variance  Std.Dev.
 family   (Intercept) 0.0001811 0.01346 
 Residual             0.0010322 0.03213 
Number of obs: 47, groups:  family, 9
```

```
Fixed effects:
             Estimate Std. Error        df t value Pr(>|t|)
(Intercept)  0.868758   0.008177 10.089475 106.247   <2e-16 ***
f1_morphG    0.019513   0.009653 41.210270   2.021   0.0498 *
---
Signif. codes:  0 '***' 0.001 '**' 0.01 '*' 0.05 '.' 0.1 ' ' 1

Correlation of Fixed Effects:
          (Intr)
f1_morphG -0.553
```

4. Adult lifespan (Number of days from eclosion as adult to death)

```
Linear mixed model fit by REML. t-tests use Satterthwaite's method
['lmerModLmerTest']
Formula: adult_lifespan_days ~ f1_morph + (1 | family)
   Data: lifespan_hyb

REML criterion at convergence: 204

Scaled residuals:
    Min      1Q  Median      3Q     Max
-1.8932 -0.6826  0.2478  0.5014  2.2716

Random effects:
 Groups   Name        Variance Std.Dev.
 family   (Intercept) 1.237    1.112
 Residual             4.074    2.018
Number of obs: 47, groups:  family, 9

Fixed effects:
            Estimate Std. Error      df t value Pr(>|t|)
(Intercept)   6.6141     0.5733 15.3511   11.54 5.74e-09 ***
f1_morphG    -0.6238     0.6117 42.3096   -1.02    0.314
---
Signif. codes:  0 '***' 0.001 '**' 0.01 '*' 0.05 '.' 0.1 ' ' 1

Correlation of Fixed Effects:
          (Intr)
f1_morphG -0.497
```

5. Fecundity (Number of adult progeny)

```
Family: nbinom2  ( log )
Formula:          progeny_count ~ f1_morph + female_mating_age +
thorax_length +      (1 | family)
```

```
Zero inflation:                 ~1
Data: progeny_hyb

      AIC       BIC    logLik -2*log(L)  df.resid
    364.5     377.3    -175.2     350.5        39

Random effects:

Conditional model:
 Groups Name        Variance Std.Dev.
 family (Intercept) 0.04444  0.2108
Number of obs: 46, groups:  family, 9

Dispersion parameter for nbinom2 family (): 2.53

Conditional model:
                  Estimate Std. Error z value Pr(>|z|)
(Intercept)         2.7008     3.9062   0.691    0.489
f1_morphG          -0.1996     0.2403  -0.831    0.406
female_mating_age   0.0613     0.4614   0.133    0.894
thorax_length       0.6209     4.2962   0.144    0.885

Zero-inflation model:
            Estimate Std. Error z value Pr(>|z|)
(Intercept)  -1.4280     0.3763  -3.795 0.000147 ***
---
Signif. codes:  0 ‘***’ 0.001 ‘**’ 0.01 ‘*’ 0.05 ‘.’ 0.1 ‘ ’
```

#### Supplementary material 3: Top genes with biased X' allele-biased expression, with functional and GO term annotation

##### Germline

**Table S6.** Functional information for genes found to have significant X' allele-biased expression in germline tissue of gynogenic females. The Gene ID for the inversion allele and the X allele are both given. Log fold change (logFC) and the p-value is given to 4 decimal places or 3 significant figures. Column 5 and 6 contains BLAST results to the entire Non-redundant (NR) Protein database, excluding *Bradysia coprophila*, for the inversion allele and the X allele respectively. Column 7 and 8 contains GO terms from a comprehensive InterProScan to obtain protein domain-level predictions and other protein/peptide/domain motif analyses to predict functional information, for the inversion allele and the X allele respectively. Many hits to the NCBI database were to hypothetical/uncharacterised proteins, and these were left out. Any genes with significant allele-biased expression but have no hits to the NCBI database/hits only to hypothetical proteins, and no GO term results from the InterProScan, are not shown in this table. Genes where the two alleles have NCBI to different proteins, or are predicted to have different GO term annotations, is shown in bold.

| Gene ID (Inv) | Gene ID (X) | logFC (4 d.p.) | p-value (3 s.f.) | NCBI best-match protein (Inv) | NCBI best-match protein (X) | InterProScan GO Term function (Inv) | InterProScan GO Term function (X) |
|---|---|---|---|---|---|---|---|
| **jg20915** | **jg4463** | **5.2656** | / | | | **transmembrane transport** | |
| jg20999 | jg4597 | 5.4304 | 2.84E-57 | | | protein binding | protein binding |
| jg21021 | jg4624 | 0.5155 | 8.42E-04 | | | dephosphorylation; phosphatase activity; protein tyrosine/serine/threonine phosphatase activity; protein dephosphorylation | dephosphorylation; phosphatase activity; protein dephosphorylation; protein tyrosine/serine/threonine phosphatase activity |
| **jg21035** | **jg4637** | **2.5041** | **3.82E-04** | | | **regulation of DNA-templated transcription** | |
| jg21248 | jg4857 | 0.3348 | 1.47E-21 | Poly [ADP-ribose] polymerase 1 [Folsomia candida] | poly [ADP-ribose] polymerase isoform X2 [Folsomia candida] | NAD+ ADP-ribosyltransferase activity | NAD+ ADP-ribosyltransferase activity |
| jg21295 | jg4909 | 2.2214 | 3.08E-07 | | | beta-N-acetylhexosaminidase activity; carbohydrate metabolic process; hydrolase activity, hydrolyzing O-glycosyl compounds | beta-N-acetylhexosaminidase activity; carbohydrate metabolic process; hydrolase activity, hydrolyzing O-glycosyl compounds |

| | | | | | | | |
|---|---|---|---|---|---|---|---|
| jg21363 | jg4982 | 0.7642 | 5.51E-04 | | | MAP kinase activity; ATP binding; protein phosphorylation; protein kinase activity | MAP kinase activity; ATP binding; protein phosphorylation; protein kinase activity |
| **jg21433** | **jg5062** | **2.0419** | **1.13E-91** | | | **enzyme inhibitor activity; ionotropic glutamate receptor activity; serine-type endopeptidase inhibitor activity; ligand-gated monoatomic ion channel activity; cholesterol metabolic process; oxidoreductase activity** | **peptidase inhibitor activity; enzyme inhibitor activity; serine-type endopeptidase inhibitor activity** |
| jg21484 | jg5116 | 0.4024 | 1.20E-04 | | | serine-type endopeptidase activity; proteolysis | serine-type endopeptidase activity; proteolysis |
| jg21488 | jg5121 | 0.2206 | 7.37E-05 | | | damaged DNA binding; DNA repair | damaged DNA binding; DNA repair |
| jg21537 | jg5170 | -0.0598 | 1.51E-02 | | | protein binding | protein binding |
| jg21546 | jg5180 | 0.6159 | 3.80E-34 | | | calcium ion binding; transmembrane transport | transmembrane transport; calcium ion binding |
| **jg21553** | **jg5189** | **0.8734** | **1.10E-106** | | | **pseudouridine synthesis; RNA binding; RNA modification; pseudouridine synthase activity** | **pseudouridine synthesis; RNA binding; RNA modification; pseudouridine synthase activity; tRNA binding; aminoacyl-tRNA ligase activity; ATP binding; tRNA aminoacylation** |
| jg21554 | jg5185 | 1.1563 | 8.23E-52 | | | Hsp90 protein binding; protein binding | protein binding; Hsp90 protein binding |
| jg21568 | jg5199 | 3.9511 | 0 | | | nucleocytoplasmic transport | nucleocytoplasmic transport |
| jg21677 | jg5297 | 0.5021 | 3.46E-16 | | | 3-hydroxyisobutyryl-CoA hydrolase activity | 3-hydroxyisobutyryl-CoA hydrolase activity |
| **jg21709** | **jg5328** | **9.7543** | **/** | | | **nucleic acid binding; zinc ion binding** | |
| jg21732 | jg5346 | 0.4719 | 2.37E-03 | | | actin binding | actin binding |
| **jg21829** | **jg6453** | **3.7542** | **1.14E-12** | | | **protein binding** | **sleep; regulation of synaptic transmission, cholinergic; positive regulation of voltage-gated potassium channel activity** |
| **jg21847** | **jg5469** | **6.0401** | **4.67E-10** | | | **oxidoreductase activity; flavin adenine dinucleotide binding; protein dimerization activity;** | **oxidoreductase activity; flavin adenine dinucleotide binding; protein dimerization activity** |

| | | | | | | | |
|---|---|---|---|---|---|---|---|
| | | | | | | **ubiquitin-protein transferase activity; protein ubiquitination** | |
| jg21921 | jg5531 | 0.7886 | 3.73E-05 | | | protein binding | protein binding |
| jg21922 | jg5532 | 0.2091 | 2.85E-02 | | | protein binding | protein binding |
| jg21948 | jg5562 | 2.0075 | 8.31E-03 | | | carbonate dehydratase activity; zinc ion binding | carbonate dehydratase activity; zinc ion binding |
| jg21960 | jg5577 | 0.6514 | 2.23E-04 | microsomal glutathione S-transferase 1-like isoform X2 [Lucilia cuprina] | microsomal glutathione S-transferase 1-like isoform X2 [Lucilia cuprina] | | |
| **jg21966** | **jg5586** | **2.0899** | **8.79E-65** | | | | **ATP binding; ATP-dependent protein folding chaperone** |
| jg21994 | jg5614 | 0.8853 | 3.07E-16 | | | copper ion binding; copper chaperone activity | copper ion binding; copper chaperone activity |
| jg21995 | jg5615 | 0.6140 | 3.72E-32 | | | NADH dehydrogenase (ubiquinone) activity; ATP synthesis coupled electron transport; oxidoreductase activity; iron-sulfur cluster binding; oxidoreductase activity, acting on NAD(P)H | oxidoreductase activity; NADH dehydrogenase (ubiquinone) activity; ATP synthesis coupled electron transport; iron-sulfur cluster binding; oxidoreductase activity, acting on NAD(P)H |
| jg22056 | jg5675 | 0.9191 | 1.50E-29 | | | protein binding | protein binding |
| jg22233 | jg5863 | 0.4902 | 1.54E-08 | | | hydrolase activity, acting on carbon-nitrogen (but not peptide) bonds; guanine catabolic process; zinc ion binding; guanine deaminase activity; hydrolase activity | hydrolase activity, acting on carbon-nitrogen (but not peptide) bonds; hydrolase activity; guanine catabolic process; zinc ion binding; guanine deaminase activity |
| jg22260 | jg5889 | 1.9860 | 1.75E-71 | | | protein binding | protein binding |
| jg22273 | jg5899 | 8.8303 | 2.00E-08 | | | cilium movement; outer dynein arm assembly | cilium movement; outer dynein arm assembly |
| jg22319 | jg5949 | 4.7218 | / | | | metalloendopeptidase activity; proteolysis; metallopeptidase activity; zinc ion binding | metalloendopeptidase activity; proteolysis; metallopeptidase activity; zinc ion binding |
| jg22336 | jg5969 | 0.5059 | 3.86E-08 | | | FAD binding; oxidoreductase activity, acting on paired donors, with incorporation or reduction of molecular oxygen, NAD(P)H as one | FAD binding; oxidoreductase activity, acting on paired donors, with incorporation or reduction of molecular oxygen, NAD(P)H as one |

| | | | | | | | |
|---|---|---|---|---|---|---|---|
| | | | | | | donor, and incorporation of one atom of oxygen | donor, and incorporation of one atom of oxygen |
| jg22373 | jg6014 | -0.6384 | / | | | catalytic activity; branched-chain-amino-acid transaminase activity; branched-chain amino acid metabolic process | catalytic activity; branched-chain-amino-acid transaminase activity; branched-chain amino acid metabolic process |
| jg22377 | jg6019 | -0.0186 | 4.12E-06 | | | calcium ion binding; mitochondrial calcium ion transmembrane transport | calcium ion binding; mitochondrial calcium ion transmembrane transport |
| jg22464 | jg6098 | 0.3105 | 1.57E-03 | | | proteolysis; metalloexopeptidase activity | proteolysis; metalloexopeptidase activity |
| **jg22465** | **jg6100** | **2.3245** | **8.36E-05** | **myophilin [Drosophila busckii]** | | **protein binding; actin binding; actomyosin structure organization** | |
| jg22477 | jg6111 | 0.8510 | 3.95E-15 | | | metallopeptidase activity; zinc ion binding; proteolysis | proteolysis; metallopeptidase activity; zinc ion binding |
| **jg22505** | **jg6138** | **3.0544** | **3.85E-94** | | | **transaminase activity; pyridoxal phosphate binding; catalytic activity** | |
| jg22507 | jg6139 | 2.5974 | 2.43E-02 | calmodulin-A-like isoform X5 [Hermetia illucens] | calmodulin-A-like isoform X5 [Hermetia illucens] | calcium ion binding | calcium ion binding |
| **jg22540** | **jg6170** | **3.5889** | **7.66E-11** | **odorant receptor 57 [Bradysia odoriphaga]** | **odorant receptor 57 [Bradysia odoriphaga]** | | **olfactory receptor activity; odorant binding; sensory perception of smell** |
| jg22655 | jg6268 | 1.9062 | 7.23E-04 | | | serine-type endopeptidase activity; proteolysis; transmembrane transporter activity; transmembrane transport | transmembrane transporter activity; transmembrane transport |
| jg22722 | jg6335 | 4.6677 | 4.80E-27 | | | serine-type endopeptidase activity; proteolysis | serine-type endopeptidase activity; proteolysis |
| **jg22783** | **jg6391** | **0.5034** | **0.00E+00** | **neuroglian-like isoform X2 [Contarinia nasturtii]** | | | |
| jg22787 | jg6394 | 0.7582 | 4.54E-54 | | | heat shock protein binding; unfolded protein binding; protein folding; Hsp70 protein binding | heat shock protein binding; unfolded protein binding; protein folding; Hsp70 protein binding |
| jg22910 | jg6528 | 5.2727 | 1.30E-07 | | | DNA binding; regulation of DNA-templated transcription | DNA binding; regulation of DNA-templated transcription |
| jg22919 | jg6541 | 0.6208 | 1.16E-07 | | | kinase activity; NAD+ kinase activity | kinase activity; NAD+ kinase activity |

| | | | | | | | |
|---|---|---|---|---|---|---|---|
| jg23000 | jg6620 | 0.4258 | 8.72E-08 | | | metallopeptidase activity; metalloendopeptidase activity; proteolysis | metalloendopeptidase activity; proteolysis; metallopeptidase activity |
| jg23002 | jg6622 | -0.0830 | 3.59E-02 | | | metallopeptidase activity; metalloendopeptidase activity; proteolysis | metallopeptidase activity; metalloendopeptidase activity; proteolysis |
| jg23013 | jg6633 | 0.3784 | 1.30E-05 | cytochrome P450 4V2 isoform X1 [Pipistrellus kuhlii] | cytochrome P450 4V2 isoform X1 [Pipistrellus kuhlii] | iron ion binding; oxidoreductase activity, acting on paired donors, with incorporation or reduction of molecular oxygen; monooxygenase activity; heme binding | iron ion binding; oxidoreductase activity, acting on paired donors, with incorporation or reduction of molecular oxygen; monooxygenase activity; heme binding |
| jg23014 | jg6634 | 0.8587 | 4.44E-02 | | | hydrolase activity, hydrolyzing O-glycosyl compounds; carbohydrate metabolic process | hydrolase activity, hydrolyzing O-glycosyl compounds; carbohydrate metabolic process |
| jg23071 | jg6687 | -0.1769 | 3.07E-02 | | | acetyltransferase activity; N-acetyltransferase activity | acetyltransferase activity; N-acetyltransferase activity |
| jg23106 | jg6715 | 0.1221 | 3.14E-02 | | | protein binding | protein binding |
| jg23157 | jg6760 | 0.2856 | 2.14E-02 | | | ATP binding; protein kinase activity; protein phosphorylation | protein kinase activity; protein phosphorylation; ATP binding |
| jg23165 | jg6770 | 0.7662 | 1.10E-36 | | | hydrolase activity, acting on acid anhydrides, in phosphorus-containing anhydrides; metal ion binding | hydrolase activity, acting on acid anhydrides, in phosphorus-containing anhydrides; metal ion binding |
| jg23166 | jg6771 | 0.3521 | 3.15E-02 | | | calcium ion binding; cell communication; Notch binding; Notch signaling pathway; protein binding; multicellular organism development | cell communication; calcium ion binding; protein binding; Notch binding; Notch signaling pathway; multicellular organism development |
| jg23186 | jg6791 | 3.8061 | 3.63E-04 | | | chitin binding | chitin binding |
| **jg23200** | **jg6804** | **-1.3172** | **2.21E-03** | | | **signal transduction; toll-like receptor signaling pathway; transmembrane signaling receptor activity; immune response** | **signal transduction; protein binding** |
| jg23228 | jg6830 | 8.4346 | 4.53E-10 | | | ATP binding; transmembrane transport; ABC-type transporter activity | ABC-type transporter activity; ATP binding; transmembrane transport |
| **jg23237** | **jg6837** | **1.2319** | **9.46E-13** | | | **serine-type endopeptidase activity; proteolysis** | |

| | | | | | | | |
|---|---|---|---|---|---|---|---|
| **jg23325** | **jg6912** | **0.0664** | **3.13E-03** | | | | **zinc ion binding; Notch signaling pathway; protein ubiquitination** |
| jg23377 | jg6967 | 0.2595 | 1.38E-02 | general transcription factor IIH subunit 5 [Drosophila albomicans] | general transcription factor IIH subunit 5 [Drosophila albomicans] | nucleotide-excision repair; transcription initiation at RNA polymerase II promoter | nucleotide-excision repair; transcription initiation at RNA polymerase II promoter |
| jg23410 | jg7006 | 0.3839 | 8.93E-04 | | | RNA binding; nucleic acid binding | RNA binding; nucleic acid binding |
| jg23418 | jg7013 | 2.1180 | 2.50E-51 | | | catalytic activity | catalytic activity |
| jg23432 | jg7029 | 1.1814 | 2.84E-02 | neurogenic protein mastermind [Hermetia illucens] | neurogenic protein mastermind [Hermetia illucens] | | |
| jg23448 | jg7055 | 2.2391 | 2.23E-37 | | | protein binding; transcription elongation by RNA polymerase II | transcription elongation by RNA polymerase II; protein binding |
| jg23462 | jg7075 | 11.0844 | / | DNA damage-binding protein 1 [Culex pipiens pallens] | PREDICTED: echinoderm microtubule-associated protein-like CG42247 [Bactrocera latifrons] | intracellular signal transduction; nucleic acid binding; protein binding; DNA repair | intracellular signal transduction |
| jg23486 | jg7096 | 2.8441 | 3.32E-180 | | | protein binding | protein binding |
| jg23494 | jg7104 | 0.1752 | 1.13E-02 | | | regulation of translational fidelity; DNA replication; DNA repair | DNA replication; DNA repair; regulation of translational fidelity |
| **jg23498** | **jg7108** | **-0.1199** | **3.95E-38** | | | | **oxidoreductase activity** |
| jg23519 | jg7134 | 0.3839 | 3.26E-04 | | | zinc ion binding; N-acylphosphatidylethanolamine-specific phospholipase D activity | zinc ion binding; N-acylphosphatidylethanolamine-specific phospholipase D activity |
| **jg23528** | **jg7140** | **8.5581** | **1.07E-62** | **diacylglycerol kinase theta isoform X6 [Contarinia nasturtii]** | **diacylglycerol kinase theta isoforms [Contarinia nasturtii]** | **ATP-dependent diacylglycerol kinase activity; signal transduction; protein kinase C-activating G protein-coupled receptor signaling pathway; kinase activity; RNA binding; intracellular protein transport; vesicle-mediated transport; NAD+ kinase activity; clathrin adaptor activity; nucleic acid binding** | **nucleic acid binding; ATP-dependent diacylglycerol kinase activity; signal transduction; RNA binding; NAD+ kinase activity; kinase activity; protein kinase C-activating G protein-coupled receptor signaling pathway** |
| jg23535 | jg7154 | 1.0304 | 7.42E-49 | | | DNA repair | DNA repair |

| | | | | | | | |
|---|---|---|---|---|---|---|---|
| jg23555 | jg7179 | -0.3146 | 1.58E-09 | | | DNA binding; DNA-binding transcription factor activity, RNA polymerase II-specific; regulation of DNA-templated transcription | DNA binding; DNA-binding transcription factor activity, RNA polymerase II-specific; regulation of DNA-templated transcription |
| jg23569 | jg7191 | -1.4378 | 1.23E-15 | | | DNA-binding transcription factor activity; regulation of DNA-templated transcription; sequence-specific DNA binding | DNA-binding transcription factor activity; regulation of DNA-templated transcription; sequence-specific DNA binding; protein binding; intracellular signal transduction |
| jg23574 | jg7197 | 3.2664 | / | | | protein binding | protein binding |
| jg23610 | jg6776 | 7.6011 | 3.81E-09 | deleted in malignant brain tumors 1 protein [Hermetia illucens] | deleted in malignant brain tumors 1 protein [Hermetia illucens] | | |
| jg23690 | jg7309 | 3.7190 | 8.05E-04 | | | monooxygenase activity; iron ion binding; oxidoreductase activity, acting on paired donors, with incorporation or reduction of molecular oxygen; heme binding | monooxygenase activity; iron ion binding; oxidoreductase activity, acting on paired donors, with incorporation or reduction of molecular oxygen; heme binding |
| jg23700 | jg7323 | -0.4635 | 3.17E-13 | | | regulation of DNA-templated transcription | regulation of DNA-templated transcription |
| jg23708 | jg7333 | 3.0031 | 7.14E-19 | | | nucleic acid binding | nucleic acid binding |
| **jg23710** | **jg7335** | **4.0444** | **2.00E-32** | | | **transcription elongation by RNA polymerase II** | |
| **jg23761** | **jg7398** | **0.1866** | **9.69E-03** | | | | **protein binding** |
| jg23774 | jg7413 | 0.9646 | 3.34E-39 | non-structural maintenance of chromosomes element 3 homolog [Hermetia illucens] | non-structural maintenance of chromosomes element 3 homolog [Hermetia illucens] | | |
| jg23775 | jg7414 | 3.5003 | / | | | sensory perception of taste | sensory perception of taste |
| jg23813 | jg7460 | 0.3466 | 2.73E-03 | | | mRNA splicing, via spliceosome; spliceosomal snRNP assembly | mRNA splicing, via spliceosome; spliceosomal snRNP assembly |
| jg23833 | jg7479 | 5.3269 | 4.70E-27 | L-dopachrome tautomerase yellow-f2 [Ceratitis capitata] | L-dopachrome tautomerase yellow-f2 [Ceratitis capitata] | | |
| jg23916 | jg7560 | 0.3608 | 2.92E-05 | | | glucosamine 6-phosphate N-acetyltransferase activity; UDP-N- | acetyltransferase activity; glucosamine 6-phosphate N-acetyltransferase |

| | | | | | | | |
|---|---|---|---|---|---|---|---|
| | | | | | | acetylglucosamine biosynthetic process; acetyltransferase activity | activity; UDP-N-acetylglucosamine biosynthetic process |
| **jg23944** | **jg7590** | **7.8672** | **6.12E-56** | | | **tRNA binding** | |
| jg24003 | jg7641 | 0.0868 | 2.80E-02 | | | hydrolase activity, hydrolyzing O-glycosyl compounds; carbohydrate metabolic process; beta-N-acetylhexosaminidase activity | hydrolase activity, hydrolyzing O-glycosyl compounds; carbohydrate metabolic process; beta-N-acetylhexosaminidase activity |
| **jg24045** | **jg7693** | **7.4357** | **/** | **nucleoporin NUP145 [Lingula anatina]** | | | |
| **jg24120** | **jg7765** | **4.4494** | **0.00E+00** | | **rhomboid-related protein 1 [Folsomia candida]** | **odorant binding** | **serine-type endopeptidase activity; odorant binding** |

**Somatic reproductive tissue**

**Table S7.** Functional information for genes found to have significant X' allele-biased expression in somatic reproductive tissue of gynogenic females. The Gene ID for the inversion allele and the X allele are both given. Log fold change (logFC) and the p-value is given to 4 decimal places or 3 significant figures. Column 5 and 6 contains BLAST results to the entire Non-redundant (NR) Protein database, excluding *Bradysia coprophila*, for the inversion allele and the X allele respectively. Column 7 and 8 contains GO terms from a comprehensive InterProScan to obtain protein domain-level predictions and other protein/peptide/domain motif analyses to predict functional information, for the inversion allele and the X allele respectively. Many hits to the NCBI database were to hypothetical/uncharacterised proteins, and these were left out. Any genes with significant allele-biased expression but have no hits to the NCBI database/hits only to hypothetical proteins, and no GO term results from the InterProScan, are not shown in this table. Genes where the two alleles have NCBI to different proteins, or are predicted to have different GO term annotations, is shown in bold.

| Gene ID (Inv) | Gene ID (X) | logFC (4 d.p.) | p-value (3 s.f.) | NCBI best-match protein (Inv) | NCBI best-match protein (X) | InterProScan GO Term function (Inv) | InterProScan GO Term function (X) |
|---|---|---|---|---|---|---|---|
| jg21021 | jg4624 | 0.4342 | 2.58E-02 | | | dephosphorylation; phosphatase activity; protein tyrosine/serine/threonine phosphatase activity; protein dephosphorylation | dephosphorylation; phosphatase activity; protein dephosphorylation; protein tyrosine/serine/threonine phosphatase activity |
| jg21112 | jg4723 | -0.4088 | 2.10E-06 | | | protein binding | protein binding |

| | | | | | | | |
|---|---|---|---|---|---|---|---|
| jg21137 | jg4747 | 2.1360 | 9.84E-06 | neuroendocrine convertase 2 isoform X2 [Hermetia illucens] | neuroendocrine convertase 2 isoform X2 [Hermetia illucens] | serine-type endopeptidase activity; proteolysis; serine-type peptidase activity | serine-type endopeptidase activity; proteolysis |
| jg21248 | jg4857 | 0.1538 | 2.61E-02 | Poly [ADP-ribose] polymerase 1 [Folsomia candida] | poly [ADP-ribose] polymerase isoform X2 [Folsomia candida] | NAD+ ADP-ribosyltransferase activity | NAD+ ADP-ribosyltransferase activity |
| **jg21349** | **jg4971** | **7.2608** | **/** | | | **mRNA processing; hydrolase activity, acting on ester bonds; hemolysis in another organism** | **hemolysis in another organism** |
| jg21544 | jg5178 | 0.3345 | 8.43E-03 | | | protein deubiquitination; K63-linked deubiquitinase activity; protein K63-linked deubiquitination; metal-dependent deubiquitinase activity; protein binding; metallopeptidase activity | protein deubiquitination; K63-linked deubiquitinase activity; protein K63-linked deubiquitination; metal-dependent deubiquitinase activity; protein binding; metallopeptidase activity |
| **jg21553** | **jg5189** | **0.7982** | **3.41E-36** | | | **pseudouridine synthesis; RNA binding; RNA modification; pseudouridine synthase activity** | **pseudouridine synthesis; RNA binding; RNA modification; pseudouridine synthase activity; tRNA binding; aminoacyl-tRNA ligase activity; ATP binding; tRNA aminoacylation** |
| jg21554 | jg5185 | 0.9621 | 9.63E-03 | | | Hsp90 protein binding; protein binding | protein binding; Hsp90 protein binding |
| jg21703 | jg5321 | 3.7783 | 5.81E-55 | | | glycylpeptide N-tetradecanoyltransferase activity; N-terminal protein myristoylation | glycylpeptide N-tetradecanoyltransferase activity; N-terminal protein myristoylation |
| **jg21709** | **jg5328** | **6.2106** | **/** | | | **nucleic acid binding; zinc ion binding** | |
| jg21780 | jg5400 | 2.8132 | 7.89E-23 | | | peptidase inhibitor activity | peptidase inhibitor activity |
| jg21782 | jg5402 | 0.3699 | 2.79E-07 | | | hydrolase activity | hydrolase activity |
| jg21821 | jg5436 | 0.7822 | 7.61E-10 | | | transmembrane signaling receptor activity; cell surface receptor signaling pathway; G protein-coupled receptor activity; G protein-coupled receptor signaling pathway | G protein-coupled receptor activity; G protein-coupled receptor signaling pathway; transmembrane signaling receptor activity; cell surface receptor signaling pathway |
| **jg21838** | **jg5456** | **1.5329** | **1.62E-02** | **PREDICTED: disks large 1 tumor suppressor protein** | | **protein binding** | **protein binding** |

| | | | | | | | |
|---|---|---|---|---|---|---|---|
| | | | | **isoform X13 [Bactrocera latifrons]** | | | |
| **jg21847** | **jg5469** | **2.5207** | **9.76E-04** | | | **oxidoreductase activity; flavin adenine dinucleotide binding; protein dimerization activity; ubiquitin-protein transferase activity; protein ubiquitination** | **oxidoreductase activity; flavin adenine dinucleotide binding; protein dimerization activity** |
| jg21873 | jg5492 | 13.4073 | / | | | lipid metabolic process | lipid metabolic process |
| jg21921 | jg5531 | 0.6613 | 5.08E-06 | | | protein binding | protein binding |
| **jg21966** | **jg5586** | **-2.0725** | **7.87E-04** | | | | **ATP binding; ATP-dependent protein folding chaperone** |
| jg22052 | jg5672 | 1.4854 | 5.51E-06 | | | iron ion binding; oxidoreductase activity, acting on paired donors, with incorporation or reduction of molecular oxygen; monooxygenase activity; heme binding | iron ion binding; oxidoreductase activity, acting on paired donors, with incorporation or reduction of molecular oxygen; monooxygenase activity; heme binding |
| **jg22071** | **jg5692** | **-1.7133** | **1.27E-03** | | | | **ATP binding; hydrolase activity** |
| jg22157 | jg5788 | 1.9236 | 3.45E-02 | | | protein binding | protein binding |
| jg22158 | jg5789 | 3.4909 | 2.41E-02 | | | GTPase activity; GTP binding | GTPase activity; GTP binding |
| jg22260 | jg5889 | 1.5380 | 5.42E-03 | | | protein binding | protein binding |
| jg22273 | jg5899 | 5.2376 | 7.26E-05 | | | cilium movement; outer dynein arm assembly | cilium movement; outer dynein arm assembly |
| **jg22327** | **jg5960** | **6.4315** | **9.84E-114** | | | **oxidoreductase activity; 1-pyrroline-5-carboxylate dehydrogenase activity; proline catabolic process to glutamate; fatty acid elongase activity; oxidoreductase activity, acting on the aldehyde or oxo group of donors, NAD or NADP as acceptor** | **oxidoreductase activity; 1-pyrroline-5-carboxylate dehydrogenase activity; proline catabolic process to glutamate; oxidoreductase activity, acting on the aldehyde or oxo group of donors, NAD or NADP as acceptor** |
| jg22350 | jg5982 | 0.5212 | 1.14E-07 | | | protein binding | protein binding |
| jg22370 | jg6010 | 0.9602 | 8.54E-03 | | | catalytic activity; amino acid metabolic process; transaminase activity; pyridoxal phosphate binding; biosynthetic process | amino acid metabolic process; transaminase activity; pyridoxal phosphate binding; biosynthetic process; catalytic activity |

| | | | | | | | |
|---|---|---|---|---|---|---|---|
| **jg22465** | **jg6100** | **2.1793** | **1.11E-125** | **myophilin [Drosophila busckii]** | | **protein binding; actin binding; actomyosin structure organization** | |
| jg22499 | jg6131 | 0.6791 | 2.33E-08 | | | transmembrane transporter activity; transmembrane transport | transmembrane transporter activity; transmembrane transport |
| **jg22505** | **jg6138** | **6.9015** | **2.94E-23** | | | **transaminase activity; pyridoxal phosphate binding; catalytic activity** | |
| jg22510 | jg6143 | -0.2211 | 5.85E-66 | | | lipase activity | lipase activity |
| jg22581 | jg6205 | 0.2623 | 7.57E-10 | | | intracellular cholesterol transport | intracellular cholesterol transport |
| **jg22655** | **jg6268** | **1.4940** | **1.86E-02** | | | **serine-type endopeptidase activity; proteolysis; transmembrane transporter activity; transmembrane transport** | **transmembrane transporter activity; transmembrane transport** |
| jg22673 | jg6284 | 3.6068 | 2.12E-13 | | | rRNA processing; U3 snoRNA binding; protein binding | protein binding; rRNA processing; U3 snoRNA binding |
| jg22688 | jg6302 | 0.7972 | 4.00E-04 | | | ADP-ribose diphosphatase activity | ADP-ribose diphosphatase activity |
| jg22722 | jg6335 | 0.3643 | 1.36E-34 | | | serine-type endopeptidase activity; proteolysis | serine-type endopeptidase activity; proteolysis |
| jg22799 | jg6404 | 0.2545 | 4.16E-05 | major royal jelly protein 2-like [Contarinia nasturtii] | major royal jelly protein 2-like [Contarinia nasturtii] | | |
| **jg22814** | **jg6421** | **0.7042** | **3.21E-03** | | **multiple inositol polyphosphate phosphatase 1-like [Contarinia nasturtii]** | | **phosphatase activity** |
| jg22901 | jg6519 | 1.7311 | 1.52E-63 | | | metalloaminopeptidase activity; hydrolase activity | hydrolase activity; metalloaminopeptidase activity |
| jg22910 | jg6528 | 3.3423 | 4.14E-02 | | | DNA binding; regulation of DNA-templated transcription | DNA binding; regulation of DNA-templated transcription |
| jg22914 | jg6534 | 2.5519 | 2.68E-02 | | | protein binding | protein binding |
| jg22970 | jg6588 | 0.6129 | 5.44E-04 | | | hydrolase activity, acting on carbon-nitrogen (but not peptide) bonds, in linear amides; nitrogen compound metabolic process | nitrogen compound metabolic process; hydrolase activity, acting on carbon-nitrogen (but not peptide) bonds, in linear amides |

| | | | | | | | |
|---|---|---|---|---|---|---|---|
| jg23002 | jg6622 | -0.0236 | 2.98E-02 | | | metallopeptidase activity; metalloendopeptidase activity; proteolysis | metallopeptidase activity; metalloendopeptidase activity; proteolysis |
| jg23006 | jg6626 | 4.8121 | 2.18E-07 | | | serine-type endopeptidase activity; proteolysis | serine-type endopeptidase activity; proteolysis |
| jg23024 | jg6641 | 0.1961 | 9.59E-04 | | | protein binding; serine-type endopeptidase activity; proteolysis | protein binding; serine-type endopeptidase activity; proteolysis |
| jg23028 | jg6645 | 1.1675 | 6.19E-07 | | | transmembrane transporter activity; transmembrane transport | transmembrane transporter activity; transmembrane transport |
| jg23040 | jg6657 | 2.6976 | 1.90E-02 | | | oxidoreductase activity; protein binding | oxidoreductase activity; protein binding |
| jg23106 | jg6715 | 0.1188 | 1.48E-03 | | | protein binding | protein binding |
| jg23225 | jg6828 | 3.4087 | 1.60E-08 | | | monooxygenase activity; iron ion binding; oxidoreductase activity, acting on paired donors, with incorporation or reduction of molecular oxygen; heme binding | monooxygenase activity; iron ion binding; oxidoreductase activity, acting on paired donors, with incorporation or reduction of molecular oxygen; heme binding |
| jg23228 | jg6830 | 6.1106 | 6.09E-03 | | | ATP binding; transmembrane transport; ABC-type transporter activity | ABC-type transporter activity; ATP binding; transmembrane transport |
| jg23238 | jg6840 | 0.7046 | 1.50E-25 | | | UDP-glycosyltransferase activity | UDP-glycosyltransferase activity |
| jg23457 | jg7068 | 0.2153 | 2.58E-35 | | | defense response | defense response |
| **jg23462** | **jg7075** | **5.3042** | **7.86E-04** | **DNA damage-binding protein 1 [Culex pipiens pallens]** | **PREDICTED: echinoderm microtubule-associated protein-like CG42247 [Bactrocera latifrons]** | **intracellular signal transduction; nucleic acid binding; protein binding; DNA repair** | **intracellular signal transduction** |
| jg23486 | jg7096 | 0.7802 | 1.41E-02 | | | protein binding | protein binding |
| **jg23498** | **jg7108** | **0.2574** | **4.80E-59** | | | | **oxidoreductase activity** |
| **jg23521** | **jg7136** | **-0.4226** | **6.60E-19** | | **transferrin [Contarinia nasturtii]** | | |
| jg23528 | jg7140 | 2.8933 | 3.39E-07 | diacylglycerol kinase theta isoform X6 [Contarinia nasturtii] | diacylglycerol kinase theta isoforms [Contarinia nasturtii] | ATP-dependent diacylglycerol kinase activity; signal transduction; protein kinase C-activating G protein-coupled receptor signaling pathway; kinase activity; RNA binding; intracellular protein | nucleic acid binding; ATP-dependent diacylglycerol kinase activity; signal transduction; RNA binding; NAD+ kinase activity; kinase activity; protein kinase C- |

| | | | | | | | |
|---|---|---|---|---|---|---|---|
| | | | | | | transport; vesicle-mediated transport; NAD+ kinase activity; clathrin adaptor activity; nucleic acid binding | activating G protein-coupled receptor signaling pathway |
| jg23569 | jg7191 | -1.6163 | 2.73E-02 | | | DNA-binding transcription factor activity; regulation of DNA-templated transcription; sequence-specific DNA binding | DNA-binding transcription factor activity; regulation of DNA-templated transcription; sequence-specific DNA binding; protein binding; intracellular signal transduction |
| jg23583 | jg7207 | 1.8180 | 3.60E-16 | | | G protein-coupled receptor activity; G protein-coupled receptor signaling pathway | G protein-coupled receptor activity; G protein-coupled receptor signaling pathway |
| jg23658 | jg7273 | 3.4227 | / | 3-hydroxyisobutyryl-CoA hydrolase, mitochondrial-like [Anopheles albimanus] | 3-hydroxyisobutyryl-CoA hydrolase [Tropilaelaps mercedesae] | | |
| jg23700 | jg7323 | -0.5929 | 4.27E-06 | | | regulation of DNA-templated transcription | regulation of DNA-templated transcription |
| **jg23710** | **jg7335** | **3.8712** | **1.75E-19** | | | **transcription elongation by RNA polymerase II** | |
| jg23711 | jg7336 | -0.1808 | 2.20E-05 | | | amino acid metabolic process; oxidoreductase activity; oxidoreductase activity, acting on the CH-NH2 group of donors, NAD or NADP as acceptor | amino acid metabolic process; oxidoreductase activity; oxidoreductase activity, acting on the CH-NH2 group of donors, NAD or NADP as acceptor |
| jg23730 | jg7359 | 0.9706 | 7.39E-15 | | | mannosyltransferase activity | mannosyltransferase activity |
| jg23769 | jg7408 | 3.1673 | 4.22E-03 | | | N,N-dimethylaniline monooxygenase activity; flavin adenine dinucleotide binding; NADP binding | flavin adenine dinucleotide binding; NADP binding; N,N-dimethylaniline monooxygenase activity |
| jg23773 | jg7412 | 3.3098 | / | | | carbonate dehydratase activity; zinc ion binding | carbonate dehydratase activity; zinc ion binding |
| jg23775 | jg7414 | 4.6039 | 2.93E-04 | | | sensory perception of taste | sensory perception of taste |
| jg23778 | jg7424 | 2.5253 | 2.50E-05 | | | protein binding | protein binding |
| jg23833 | jg7479 | 3.0057 | 5.34E-12 | L-dopachrome tautomerase yellow-f2 [Ceratitis capitata] | L-dopachrome tautomerase yellow-f2 [Ceratitis capitata] | | |

| | | | | | | | |
|---|---|---|---|---|---|---|---|
| jg23871 | jg7519 | 0.2406 | 2.77E-04 | | | calcium ion binding | calcium ion binding |
| jg23894 | jg7539 | 1.3724 | 2.79E-04 | | | G protein-coupled receptor activity; G protein-coupled receptor signaling pathway | G protein-coupled receptor activity; G protein-coupled receptor signaling pathway |
| jg23919 | jg7562 | -1.0078 | 2.56E-07 | | | DNA-binding transcription factor activity; regulation of DNA-templated transcription; DNA binding; regulation of transcription by RNA polymerase II | DNA-binding transcription factor activity; regulation of DNA-templated transcription; DNA binding; regulation of transcription by RNA polymerase II; protein kinase activity; ATP binding; protein phosphorylation |
| **jg24045** | **jg7693** | **6.2564** | / | **nucleoporin NUP145 [Lingula anatina]** | | | |
| jg24051 | jg7702 | 0.7920 | 2.94E-14 | | | monooxygenase activity; iron ion binding; oxidoreductase activity, acting on paired donors, with incorporation or reduction of molecular oxygen; heme binding | monooxygenase activity; iron ion binding; oxidoreductase activity, acting on paired donors, with incorporation or reduction of molecular oxygen; heme binding |
| jg24102 | jg7749 | 9.3402 | 6.67E-05 | probable G-protein coupled receptor Mth-like 1 [Aedes aegypti] | odorant-binding protein 16 [Bradysia odoriphaga] | odorant binding; G protein-coupled receptor activity; G protein-coupled receptor signaling pathway; transmembrane signaling receptor activity; cell surface receptor signaling pathway | odorant binding |
| jg24120 | jg7765 | 4.4410 | 1.02E-35 | | | odorant binding | serine-type endopeptidase activity; odorant binding |
| jg24325 | jg7985 | 4.7862 | 1.62E-05 | | | protein binding | protein binding |

Somatic non-reproductive tissue

**Table S8.** Functional information for genes found to have significant X' allele-biased expression in somatic non-reproductive tissue of gynogenic females. The Gene ID for the inversion allele and the X allele are both given. Log fold change (logFC) and the p-value is given to 4 decimal places or 3 significant figures. Column 5 and 6 contains BLAST results to the entire Non-redundant (NR) Protein database, excluding *Bradysia coprophila*, for the inversion allele and the X allele respectively. Column 7 and 8 contains GO terms from a comprehensive InterProScan to obtain protein domain-level predictions and other protein/peptide/domain motif analyses to predict functional information, for the inversion allele and the X allele respectively. Many hits to the NCBI database were to hypothetical/uncharacterised proteins, and these were left out. Any genes with significant allele-biased expression but have no hits to the NCBI database/hits only to hypothetical proteins, and no GO term results from the InterProScan, are not shown in this table. Genes where the two alleles have NCBI to different proteins, or are predicted to have different GO term annotations, is shown in bold.

| Gene ID (Inv) | Gene ID (X) | logFC (4 d.p.) | p-value (3 s.f.) | NCBI best-match protein (Inv) | NCBI best-match protein (X) | InterProScan GO Term function (Inv) | InterProScan GO Term function (X) |
|---|---|---|---|---|---|---|---|
| jg20906 | jg4386 | 3.3571 | / | reverse transcriptase [Rhynchosciara americana] | reverse transcriptase [Rhynchosciara americana] | | |
| jg20934 | jg4539 | 0.9647 | 1.85E-04 | | | transmembrane transporter activity; transmembrane transport | transmembrane transporter activity; transmembrane transport |
| jg21112 | jg4723 | 0.0985 | 8.41E-95 | | | protein binding | protein binding |
| jg21137 | jg4747 | 2.2226 | 1.40E-12 | neuroendocrine convertase 2 isoform X2 [Hermetia illucens] | neuroendocrine convertase 2 isoform X2 [Hermetia illucens] | serine-type endopeptidase activity; proteolysis; serine-type peptidase activity | serine-type endopeptidase activity; proteolysis |
| jg21248 | jg4857 | 0.2809 | 1.96E-04 | poly [ADP-ribose] polymerase isoform X1 [Folsomia candida] | poly [ADP-ribose] polymerase isoform X2 [Folsomia candida] | NAD+ ADP-ribosyltransferase activity | NAD+ ADP-ribosyltransferase activity |
| jg21281 | jg4892 | 0.4208 | 6.67E-03 | | | tail-anchored membrane protein insertion into ER membrane | tail-anchored membrane protein insertion into ER membrane |
| **jg21553** | **jg5189** | **0.7952** | **1.12E-49** | | | **pseudouridine synthesis; RNA binding; RNA modification; pseudouridine synthase activity** | **pseudouridine synthesis; RNA binding; RNA modification; pseudouridine synthase activity; tRNA binding; aminoacyl-tRNA ligase activity; ATP binding; tRNA aminoacylation** |
| jg21554 | jg5185 | 1.1078 | 4.41E-02 | | | Hsp90 protein binding; protein binding | protein binding; Hsp90 protein binding |
| jg21576 | jg5203 | 2.3592 | 2.86E-09 | | | glycine hydroxymethyltransferase activity; glycine biosynthetic process from serine; pyridoxal phosphate binding; tetrahydrofolate interconversion; catalytic activity | glycine hydroxymethyltransferase activity; glycine biosynthetic process from serine; pyridoxal phosphate binding; tetrahydrofolate interconversion; catalytic activity |
| jg21703 | jg5321 | 3.7456 | 2.77E-43 | | | glycylpeptide N-tetradecanoyltransferase activity; N-terminal protein myristoylation | glycylpeptide N-tetradecanoyltransferase activity; N-terminal protein myristoylation |
| **jg21709** | **jg5328** | **7.1622** | **/** | | | **nucleic acid binding; zinc ion binding** | |

| | | | | | | | |
|---|---|---|---|---|---|---|---|
| jg21780 | jg5400 | 3.0083 | 3.88E-116 | | | peptidase inhibitor activity | peptidase inhibitor activity |
| jg21782 | jg5402 | 0.9450 | 7.60E-14 | | | hydrolase activity | hydrolase activity |
| jg21822 | jg5437 | -0.0643 | 4.18E-05 | tubulin alpha-1A chain [Aedes aegypti] | tubulin alpha-1A chain [Aedes aegypti]; tubulin alpha-2 chain [Culex quinquefasciatus] | structural constituent of cytoskeleton; GTP binding; microtubule-based process | GTP binding; microtubule-based process; structural constituent of cytoskeleton |
| **jg21847** | **jg5469** | **4.5207** | **5.39E-06** | | | **oxidoreductase activity; flavin adenine dinucleotide binding; protein dimerization activity; ubiquitin-protein transferase activity; protein ubiquitination** | **oxidoreductase activity; flavin adenine dinucleotide binding; protein dimerization activity** |
| jg21873 | jg5492 | 3.3571 | 6.00E-03 | | | lipid metabolic process | lipid metabolic process |
| jg21960 | jg5577 | 0.4105 | 1.79E-03 | microsomal glutathione S-transferase 1-like isoform X2 [Lucilia cuprina] | microsomal glutathione S-transferase 1-like isoform X2 [Lucilia cuprina] | | |
| **jg21966** | **jg5586** | **-2.3735** | **4.33E-08** | | | | **ATP binding; ATP-dependent protein folding chaperone** |
| jg22052 | jg5672 | 1.6214 | 4.52E-30 | | | iron ion binding; oxidoreductase activity, acting on paired donors, with incorporation or reduction of molecular oxygen; monooxygenase activity; heme binding | iron ion binding; oxidoreductase activity, acting on paired donors, with incorporation or reduction of molecular oxygen; monooxygenase activity; heme binding |
| jg22053 | jg5673 | 0.9031 | 1.86E-12 | | | monooxygenase activity; iron ion binding; oxidoreductase activity, acting on paired donors, with incorporation or reduction of molecular oxygen; heme binding | monooxygenase activity; iron ion binding; oxidoreductase activity, acting on paired donors, with incorporation or reduction of molecular oxygen; heme binding |
| jg22135 | jg5761 | 0.7127 | 5.12E-11 | | | transmembrane transporter activity; transmembrane transport | transmembrane transporter activity; transmembrane transport |
| jg22158 | jg5789 | 0.7531 | 2.22E-03 | | | GTPase activity; GTP binding | GTPase activity; GTP binding |
| jg22191 | jg5825 | 2.2818 | 1.46E-04 | | | protein binding | protein binding |
| jg22234 | jg5864 | 1.2335 | 3.60E-02 | | | olfactory receptor activity; odorant binding; sensory perception of smell | olfactory receptor activity; odorant binding; sensory perception of smell |
| jg22260 | jg5889 | 1.9368 | 8.35E-15 | | | protein binding | protein binding |

| | | | | | | | |
|---|---|---|---|---|---|---|---|
| jg22273 | jg5899 | 6.3913 | 1.72E-04 | | | cilium movement; outer dynein arm assembly | cilium movement; outer dynein arm assembly |
| **jg22288** | **jg5916** | **3.0378** | **1.53E-06** | | **cytochrome P450 [Chironomus tentans]** | **monooxygenase activity; iron ion binding; oxidoreductase activity, acting on paired donors, with incorporation or reduction of molecular oxygen; heme binding; calcium ion binding; enzyme regulator activity** | **monooxygenase activity; iron ion binding; oxidoreductase activity, acting on paired donors, with incorporation or reduction of molecular oxygen; heme binding** |
| jg22305 | jg5936 | 2.6566 | 2.99E-02 | | | serine-type endopeptidase activity; proteolysis | serine-type endopeptidase activity; proteolysis |
| jg22316 | jg5946 | 0.0073 | 6.17E-04 | | | sequence-specific DNA binding; DNA-binding transcription factor activity; regulation of transcription by RNA polymerase II; regulation of DNA-templated transcription | sequence-specific DNA binding; DNA-binding transcription factor activity; regulation of DNA-templated transcription; regulation of transcription by RNA polymerase II |
| jg22327 | jg5960 | 5.8793 | 3.23E-170 | | | oxidoreductase activity; 1-pyrroline-5-carboxylate dehydrogenase activity; proline catabolic process to glutamate; fatty acid elongase activity; oxidoreductase activity, acting on the aldehyde or oxo group of donors, NAD or NADP as acceptor | oxidoreductase activity; 1-pyrroline-5-carboxylate dehydrogenase activity; proline catabolic process to glutamate; oxidoreductase activity, acting on the aldehyde or oxo group of donors, NAD or NADP as acceptor |
| jg22350 | jg5982 | 0.2768 | 3.46E-06 | | | protein binding | protein binding |
| jg22369 | jg6009 | 6.0112 | 2.25E-16 | 4-coumarate--CoA ligase 2-like [Contarinia nasturtii] | 4-coumarate--CoA ligase 2-like [Contarinia nasturtii] | | |
| jg22394 | jg6032 | 0.3187 | 2.29E-02 | | | peptidyl-prolyl cis-trans isomerase activity | peptidyl-prolyl cis-trans isomerase activity |
| jg22400 | jg6036 | 1.6000 | 4.77E-02 | | | serine-type endopeptidase activity; proteolysis | serine-type endopeptidase activity; proteolysis |
| **jg22465** | **jg6100** | **2.3278** | **4.37E-246** | **myophilin [Drosophila busckii]** | | **protein binding; actin binding; actomyosin structure organization** | |
| **jg22505** | **jg6138** | **9.1102** | **9.21E-69** | | | **transaminase activity; pyridoxal phosphate binding; catalytic activity** | |

| | | | | | | | |
|---|---|---|---|---|---|---|---|
| jg22510 | jg6143 | -0.2602 | 2.15E-38 | | | lipase activity | lipase activity |
| jg22581 | jg6205 | 0.2460 | 1.09E-06 | | | intracellular cholesterol transport | intracellular cholesterol transport |
| jg22722 | jg6335 | 0.5170 | 2.99E-23 | | | serine-type endopeptidase activity; proteolysis | serine-type endopeptidase activity; proteolysis |
| jg22730 | jg6339 | 0.4490 | 3.81E-03 | | | ligand-gated monoatomic ion channel activity; ionotropic glutamate receptor activity | ligand-gated monoatomic ion channel activity; ionotropic glutamate receptor activity |
| **jg22733** | **jg6343** | **8.4303** | **/** | | | **magnesium ion binding; tRNA modification; tRNA guanylyltransferase activity** | |
| jg22758 | jg6370 | 0.3145 | 1.31E-02 | | | actin cytoskeleton organization; actin filament binding; protein binding | protein binding; actin cytoskeleton organization; actin filament binding |
| **jg22783** | **jg6391** | **0.6285** | **1.64E-170** | **neuroglian-like isoform X2 [Contarinia nasturtii]** | | | |
| **jg22814** | **jg6421** | **0.6296** | **1.22E-02** | | **multiple inositol polyphosphate phosphatase 1-like [Contarinia nasturtii]** | | **phosphatase activity** |
| jg22855 | jg6463 | 3.2664 | / | | | G protein-coupled receptor activity; G protein-coupled receptor signaling pathway | G protein-coupled receptor activity; G protein-coupled receptor signaling pathway |
| jg22901 | jg6519 | 1.1611 | 2.22E-02 | | | metalloaminopeptidase activity; hydrolase activity | hydrolase activity; metalloaminopeptidase activity |
| jg22914 | jg6534 | 2.9282 | / | | | protein binding | protein binding |
| **jg22957** | **jg6573** | **4.7789** | **/** | | | **chitin binding** | |
| jg23002 | jg6622 | 0.2754 | 1.96E-02 | | | metallopeptidase activity; metalloendopeptidase activity; proteolysis | metallopeptidase activity; metalloendopeptidase activity; proteolysis |
| jg23006 | jg6626 | 2.6498 | 4.10E-09 | | | serine-type endopeptidase activity; proteolysis | serine-type endopeptidase activity; proteolysis |
| jg23040 | jg6657 | 1.5145 | 4.31E-04 | | | oxidoreductase activity; protein binding | oxidoreductase activity; protein binding |
| jg23157 | jg6760 | 0.4089 | 5.32E-14 | | | ATP binding; protein kinase activity; protein phosphorylation | protein kinase activity; protein phosphorylation; ATP binding |

| | | | | | | | |
|---|---|---|---|---|---|---|---|
| jg23223 | jg6825 | 0.7383 | 2.72E-07 | | | O-phospho-L-serine:2-oxoglutarate aminotransferase activity; L-serine biosynthetic process; catalytic activity | catalytic activity; O-phospho-L-serine:2-oxoglutarate aminotransferase activity; L-serine biosynthetic process |
| jg23228 | jg6830 | 8.0862 | / | | | ATP binding; transmembrane transport; ABC-type transporter activity | ABC-type transporter activity; ATP binding; transmembrane transport |
| jg23410 | jg7006 | 0.2929 | 1.43E-02 | | | RNA binding; nucleic acid binding | RNA binding; nucleic acid binding |
| jg23422 | jg7019 | 0.2725 | 1.71E-03 | | | inorganic phosphate transmembrane transporter activity; phosphate ion transport | inorganic phosphate transmembrane transporter activity; phosphate ion transport |
| jg23448 | jg7055 | 2.0605 | 3.47E-07 | | | protein binding; transcription elongation by RNA polymerase II | transcription elongation by RNA polymerase II; protein binding |
| jg23457 | jg7068 | 1.4255 | 3.89E-05 | | | defense response | defense response |
| jg23471 | jg7082 | 1.1547 | 2.17E-02 | transcription factor 15-like isoform X3 [Colletes gigas]; basic helix-loop-helix transcription factor scleraxis-like isoform X2 [Coccinella septempunctata] | transcription factor 15-like isoform X3 [Colletes gigas]; basic helix-loop-helix transcription factor scleraxis-like isoform X2 [Coccinella septempunctata] | protein dimerization activity | protein dimerization activity |
| jg23486 | jg7096 | 0.8630 | 3.33E-23 | | | protein binding | protein binding |
| **jg23498** | **jg7108** | **0.2465** | **7.71E-22** | | | | **oxidoreductase activity** |
| **jg23528** | **jg7140** | **0.9157** | **1.64E-12** | **diacylglycerol kinase theta isoform X6 [Contarinia nasturtii]** | **diacylglycerol kinase theta isoforms [Contarinia nasturtii]** | **ATP-dependent diacylglycerol kinase activity; signal transduction; protein kinase C-activating G protein-coupled receptor signaling pathway; kinase activity; RNA binding; intracellular protein transport; vesicle-mediated transport; NAD+ kinase activity; clathrin adaptor activity; nucleic acid binding** | **nucleic acid binding; ATP-dependent diacylglycerol kinase activity; signal transduction; RNA binding; NAD+ kinase activity; kinase activity; protein kinase C-activating G protein-coupled receptor signaling pathway** |
| **jg23569** | **jg7191** | **-1.6490** | **8.83E-03** | | | **DNA-binding transcription factor activity; regulation of DNA-templated transcription; sequence-specific DNA binding** | **DNA-binding transcription factor activity; regulation of DNA-templated transcription; sequence-specific DNA binding;** |

| | | | | | | | |
|---|---|---|---|---|---|---|---|
| | | | | | | | **protein binding; intracellular signal transduction** |
| jg23574 | jg7197 | 1.1910 | 3.52E-02 | | | protein binding | protein binding |
| jg23625 | jg7244 | 1.5666 | 1.32E-101 | | | protein kinase activity; ATP binding; protein phosphorylation | protein kinase activity; ATP binding; protein phosphorylation |
| jg23658 | jg7273 | 3.3629 | / | 3-hydroxyisobutyryl-CoA hydrolase, mitochondrial-like [Anopheles albimanus] | 3-hydroxyisobutyryl-CoA hydrolase [Tropilaelaps mercedesae] | | |
| jg23680 | jg7299 | 0.5423 | 2.63E-02 | odorant-binding protein 33 [Bradysia odoriphaga] | odorant-binding protein 33 [Bradysia odoriphaga] | odorant binding | odorant binding |
| jg23700 | jg7323 | -0.5520 | 1.54E-04 | | | regulation of DNA-templated transcription | regulation of DNA-templated transcription |
| **jg23710** | **jg7335** | **3.7563** | **7.59E-31** | | | **transcription elongation by RNA polymerase II** | |
| jg23730 | jg7359 | 1.3157 | 4.29E-18 | | | mannosyltransferase activity | mannosyltransferase activity |
| jg23769 | jg7408 | 3.3224 | 4.95E-03 | | | N,N-dimethylaniline monooxygenase activity; flavin adenine dinucleotide binding; NADP binding | flavin adenine dinucleotide binding; NADP binding; N,N-dimethylaniline monooxygenase activity |
| jg23770 | jg7409 | 3.6832 | / | | | chitin binding | chitin binding |
| jg23771 | jg7410 | 1.7899 | 1.23E-02 | | | chitin binding | chitin binding |
| jg23774 | jg7413 | 0.6774 | 8.55E-05 | non-structural maintenance of chromosomes element 3 homolog [Hermetia illucens] | non-structural maintenance of chromosomes element 3 homolog [Hermetia illucens] | | |
| jg23775 | jg7414 | 4.9945 | / | | | sensory perception of taste | sensory perception of taste |
| jg23778 | jg7424 | 1.6344 | 6.47E-31 | | | protein binding | protein binding |
| jg23833 | jg7479 | 2.9018 | 2.74E-31 | L-dopachrome tautomerase yellow-f2 [Ceratitis capitata] | L-dopachrome tautomerase yellow-f2 [Ceratitis capitata] | | |
| jg23919 | jg7562 | -0.9735 | 2.79E-09 | | | DNA-binding transcription factor activity; regulation of DNA-templated transcription; DNA | DNA-binding transcription factor activity; regulation of DNA-templated transcription; DNA binding; regulation of transcription |

| | | | | | | | |
|---|---|---|---|---|---|---|---|
| | | | | | | binding; regulation of transcription by RNA polymerase II | by RNA polymerase II; protein kinase activity; ATP binding; protein phosphorylation |
| **jg23944** | **jg7590** | **7.1491** | **4.93E-21** | | | **tRNA binding** | |
| **jg24045** | **jg7693** | **6.0928** | **4.06E-04** | **nucleoporin NUP145 [Lingula anatina]** | | | |
| **jg24102** | **jg7749** | **8.7233** | **6.15E-14** | **probable G-protein coupled receptor Mth-like 1 [Aedes aegypti]** | **odorant-binding protein 16 [Bradysia odoriphaga]** | **odorant binding; G protein-coupled receptor activity; G protein-coupled receptor signaling pathway; transmembrane signaling receptor activity; cell surface receptor signaling pathway** | **odorant binding** |
| **jg24120** | **jg7765** | **4.8069** | **3.39E-85** | | | **odorant binding** | **serine-type endopeptidase activity; odorant binding** |

450

#### Supplementary material 4: Top autosomal differentially expressed genes, with functional annotation and GO term enrichment analysis

##### Germline

**Table S2.** List of autosomal genes found to be significantly differentially expressed between gynogenic and androgenic females in germline tissue, and the chromosome it's found on. Log fold change (logFC) and the false discovery rate (i.e. adjusted p-value) is given to 4 decimal places. Column 5 contains BLAST results for the protein to the entire Non-redundant (NR) Protein database, excluding *Bradysia coprophila*. Column 6 contains GO terms that result from a comprehensive InterProScan to obtain protein domain-level predictions and other protein/peptide/domain motif analyses to predict functional information. Many hits to the NCBI database were to hypothetical/uncharacterised proteins, and these were left out. Any genes there are significantly differentially expressed but have no hits to the NCBI database/hits only to hypothetical proteins, and no GO term results from the InterProScan, are not shown in this table.

| Gene ID | Chromo-some | logFC (4 d.p.) | FDR (4 d.p.) | NCBI best-match protein | InterProScan GO Term function |
|---|---|---|---|---|---|
| jg166 | II | 3.307783 | 0.028845 | piggyBac transposable element-derived protein 3-like [Portunus trituberculatus] | |
| jg2255 | II | 3.089132 | 0.012914 | | oxidoreductase activity, acting on CH-OH group of donors; flavin adenine dinucleotide binding |
| jg2861 | II | 3.027023 | 0.009131 | | monooxygenase activity; iron ion binding; oxidoreductase activity, acting on paired donors, with incorporation or reduction of molecular oxygen; heme binding |
| jg11334 | IV | 4.186951 | 0.037586 | | nucleic acid binding |
| jg11564 | IV | 2.451545 | 0.022028 | | catalytic activity; iron-sulfur cluster binding |
| jg11827 | IV | 2.492456 | 0.009131 | | antiporter activity; xenobiotic transmembrane transporter activity; transmembrane transport |
| jg18699 | III | 1.770699 | 0.009131 | | GTP binding; microtubule-based process; structural constituent of cytoskeleton |
| jg18809 | III | 1.521185 | 0.012914 | | nucleotidyltransferase activity |
| jg19292 | III | 3.595996 | 0.009131 | | RNA binding; adenosine deaminase activity; RNA processing; protein binding |

**Table S3.** Results of GO-term enrichment analysis on significantly differentially expressed genes between gynogenic and androgenic females in germline tissue. The p-value is obtained with a Classic Fisher test.

| GO:ID | Term | Category | No. of genes annotated with this GO term | No. of significantly expressed genes with this GO term | Expected no. of significantly expressed genes with this GO term | p-value |
|---|---|---|---|---|---|---|
| GO:0006396 | RNA processing | Biological Process | 144 | 1 | 0.05 | 0.05 |
| GO:0042910 | xenobiotic transmembrane transporter activity | Molecular Function | 2 | 1 | 0 | 0.0024 |
| GO:0004000 | adenosine deaminase activity | Molecular Function | 4 | 1 | 0 | 0.0049 |

**Somatic non-reproductive tissue**

**Table S4.** List of autosomal genes found to be significantly differentially expressed between gynogenic and androgenic females in somatic non-reproductive tissue, and the chromosome it's found on. Log fold change (logFC) and the false discovery rate (i.e. adjusted p-value) is given to 4 decimal places. Column 5 contains BLAST results for the protein to the entire Non-redundant (NR) Protein database, excluding *Bradysia coprophila*. Column 6 contains GO terms that result from a comprehensive InterProScan to obtain protein domain-level predictions and other protein/peptide/domain motif analyses to predict functional information. Many hits to the NCBI database were to hypothetical/uncharacterised proteins, and these were left out. Any genes there are significantly differentially expressed but have no hits to the NCBI database/hits only to hypothetical proteins, and no GO term results from the InterProScan, are not shown in this table.

| Gene ID | Chromo-some | logFC (4 d.p.) | FDR (4 d.p.) | NCBI best-match protein | InterProScan GO Term function |
|---|---|---|---|---|---|
| jg102 | II | 0.6243 | 0.0436 | | DNA-binding transcription factor activity; regulation of DNA-templated transcription |
| jg166 | II | 3.2766 | 0.0157 | piggyBac transposable element-derived protein 3-like [Portunus trituberculatus] | |
| jg328 | II | 1.0151 | 0.0287 | | acyltransferase activity, transferring groups other than amino-acyl groups |
| jg354 | II | 0.8056 | 0.0161 | | UDP-glycosyltransferase activity |
| jg460 | II | -0.6281 | 0.0488 | | iron ion binding; DNA binding; zinc ion binding; protein binding |

| | | | | | |
|---|---|---|---|---|---|
| jg480 | II | 0.7106 | 0.0194 | | zinc ion binding; catabolic process; hydrolase activity, acting on ester bonds |
| jg526 | II | -0.7101 | 0.0493 | | proteolysis; cysteine-type peptidase activity |
| jg607 | II | 2.2293 | 0.0194 | | transmembrane transporter activity; transmembrane transport |
| jg608 | II | 2.3531 | 0.0155 | | catalytic activity; amino acid metabolic process; carboxy-lyase activity; carbon-carbon lyase activity; carboxylic acid metabolic process; pyridoxal phosphate binding |
| jg609 | II | 2.5952 | 0.0245 | | transmembrane transporter activity; transmembrane transport |
| jg616 | II | 0.5909 | 0.0471 | | metal ion transport; metal ion transmembrane transporter activity |
| jg646 | II | 1.6754 | 0.0037 | | transmembrane transporter activity; transmembrane transport |
| jg740 | II | 2.8875 | 0.0113 | | transmembrane transporter activity; transmembrane transport |
| jg742 | II | 1.3102 | 0.0287 | | transmembrane transporter activity; transmembrane transport |
| jg747 | II | 0.4986 | 0.0454 | | protein kinase activity; ATP binding; protein phosphorylation; phosphorylase kinase activity; calmodulin binding; glycogen biosynthetic process |
| jg882 | II | 0.5598 | 0.0475 | | flavin adenine dinucleotide binding; oxidoreductase activity, acting on CH-OH group of donors |
| jg929 | II | 0.8185 | 0.0377 | | ammonium transmembrane transporter activity; ammonium transmembrane transport |
| jg960 | II | 0.9489 | 0.0489 | galectin [Culex quinquefasciatus] | carbohydrate binding |
| jg1007 | II | 0.8517 | 0.0304 | | sulfotransferase activity |
| jg1099 | II | 1.0894 | 0.0446 | | nucleic acid binding |
| jg1223 | II | 1.4155 | 0.0110 | | glutathione catabolic process; glutathione hydrolase activity |
| jg1301 | II | 0.9743 | 0.0258 | | monooxygenase activity; iron ion binding; oxidoreductase activity, acting on paired donors, with incorporation or reduction of molecular oxygen; heme binding |
| jg1327 | II | -0.9656 | 0.0362 | | protein kinase activity; ATP binding; protein phosphorylation |
| jg1330 | II | 0.8537 | 0.0416 | ATP synthase subunit O, mitochondrial [Drosophila navojoa] | proton motive force-driven ATP synthesis; proton-transporting ATP synthase activity, rotational mechanism |
| jg1332 | II | 0.8790 | 0.0489 | arginine kinase isoform X1 [Anopheles albimanus] | kinase activity; transferase activity, transferring phosphorus-containing groups; creatine kinase activity; phosphocreatine biosynthetic process; catalytic activity |
| jg1339 | II | -0.7942 | 0.0377 | | protein kinase activity; ATP binding; protein phosphorylation |
| jg1358 | II | 0.6431 | 0.0466 | | NADP binding; phosphogluconate dehydrogenase (decarboxylating) activity; pentose-phosphate shunt; oxidoreductase activity |
| jg1361 | II | 0.7776 | 0.0498 | | catalytic activity; flavin adenine dinucleotide binding; helicase activity |
| jg1363 | II | 0.9867 | 0.0155 | | flavin adenine dinucleotide binding; FAD binding; catalytic activity |

| | | | | | |
|---|---|---|---|---|---|
| jg1372 | II | 2.4372 | 0.0466 | | transmembrane transporter activity; transmembrane transport |
| jg1385 | II | 0.6609 | 0.0477 | | transmembrane transporter activity; transmembrane transport |
| jg1459 | II | 0.8785 | 0.0148 | | transmembrane transporter activity; transmembrane transport |
| jg1497 | II | 1.7001 | 0.0223 | | iron ion binding; oxidoreductase activity, acting on paired donors, with incorporation or reduction of molecular oxygen; monooxygenase activity; heme binding |
| jg1535 | II | 0.9547 | 0.0277 | | nitrogen compound metabolic process |
| jg1563 | II | 1.3251 | 0.0467 | | protein binding |
| jg1604 | II | 2.4640 | 0.0359 | | selenium binding |
| jg1692 | II | 1.6352 | 0.0472 | juvenile hormone, partial [Bradysia odoriphaga] | |
| jg1747 | II | -0.7768 | 0.0340 | | protein binding; nucleic acid binding; RNA binding; histone H3K4 methyltransferase activity; histone H3-K4 methylation |
| jg1800 | II | 0.9914 | 0.0374 | Levanase [Orchesella cincta] | hydrolase activity, hydrolyzing O-glycosyl compounds; carbohydrate metabolic process |
| jg1911 | II | -1.1114 | 0.0336 | | protein binding |
| jg2019 | II | 1.1455 | 0.0374 | | antiporter activity; xenobiotic transmembrane transporter activity; transmembrane transport |
| jg2033 | II | 0.6212 | 0.0488 | | monoatomic ion transport; transmembrane transporter activity; chloride:monoatomic cation symporter activity; transmembrane transport |
| jg2225 | II | 0.7300 | 0.0258 | | transmembrane transporter activity; transmembrane transport |
| jg2230 | II | 1.0528 | 0.0144 | | methyltransferase activity; nucleic acid binding; methylation |
| jg2317 | II | 0.9806 | 0.0498 | | hydrolase activity |
| jg2392 | II | -0.6877 | 0.0489 | | protein binding |
| jg2430 | II | 3.0899 | 0.0061 | transmembrane protease serine 9-like [Frankliniella occidentalis] | serine-type endopeptidase activity; proteolysis |
| jg2448 | II | 0.8616 | 0.0377 | CAPA receptor [Aedes aegypti] | G protein-coupled receptor activity; G protein-coupled receptor signaling pathway; neuromedin U receptor activity |
| jg2476 | II | 0.7724 | 0.0303 | | sulfate assimilation; sulfate adenylyltransferase (ATP) activity; adenylylsulfate kinase activity; ATP binding |
| jg2510 | II | 1.4421 | 0.0025 | | transmembrane transporter activity; transmembrane transport; phosphorelay sensor kinase activity; phosphorelay signal transduction system; protein histidine kinase activity; cell wall organization |
| jg2684 | II | 0.7778 | 0.0477 | | inward rectifier potassium channel activity; potassium ion transport |

| | | | | | |
|---|---|---|---|---|---|
| jg2795 | II | 1.0474 | 0.0105 | acetyl-coenzyme A transporter 1 [Contarinia nasturtii] | acetyl-CoA transmembrane transporter activity |
| jg2861 | II | 1.4430 | 0.0309 | | monooxygenase activity; iron ion binding; oxidoreductase activity, acting on paired donors, with incorporation or reduction of molecular oxygen; heme binding |
| jg2887 | II | 0.9867 | 0.0461 | | glucose metabolic process; oxidoreductase activity, acting on CH-OH group of donors; NADP binding; glucose-6-phosphate dehydrogenase activity |
| jg3088 | II | 0.7365 | 0.0415 | | proteolysis; metallopeptidase activity; peptidyl-dipeptidase activity |
| jg3308 | II | 0.6394 | 0.0442 | | regulation of DNA-templated transcription; sequence-specific DNA binding; DNA-binding transcription factor activity; regulation of transcription by RNA polymerase II; zinc ion binding |
| jg3316 | II | 1.8205 | 0.0169 | | transmembrane transporter activity; transmembrane transport |
| jg3530 | II | 1.8700 | 0.0381 | | transsulfuration; pyridoxal phosphate binding; catalytic activity |
| jg3540 | II | 2.8653 | 0.0180 | | nucleic acid binding |
| jg3577 | II | 0.8399 | 0.0245 | | iron ion binding; oxidoreductase activity, acting on paired donors, with incorporation or reduction of molecular oxygen; monooxygenase activity; heme binding |
| jg3656 | II | 1.3776 | 0.0367 | open rectifier potassium channel protein 1 [Culex pipiens pallens] | potassium channel activity; potassium ion transmembrane transport |
| jg3707 | II | 0.8156 | 0.0309 | | transferase activity |
| jg3710 | II | 0.9411 | 0.0461 | | oxidoreductase activity |
| jg3851 | II | -0.8731 | 0.0489 | | protein phosphatase regulator activity |
| jg3891 | II | 1.0122 | 0.0493 | | transmembrane transporter activity; transmembrane transport |
| jg3932 | II | 1.3812 | 0.0144 | | oxidoreductase activity, acting on the CH-CH group of donors; flavin adenine dinucleotide binding; acyl-CoA oxidase activity; fatty acid metabolic process; FAD binding; fatty acid beta-oxidation |
| jg4047 | II | -0.7764 | 0.0475 | | RNA processing; polynucleotide 5'-hydroxyl-kinase activity |
| jg8308 | IV | 3.0943 | 0.0305 | | serine-type endopeptidase inhibitor activity |
| jg8382 | IV | 0.7077 | 0.0275 | | monoatomic ion transport; extracellular ligand-gated monoatomic ion channel activity; transmembrane signaling receptor activity; monoatomic ion channel activity; monoatomic ion transmembrane transport |
| jg8444 | IV | 1.3062 | 0.0245 | | transsulfuration; pyridoxal phosphate binding; catalytic activity |
| jg8462 | IV | 1.0431 | 0.0193 | | transmembrane transporter activity; transmembrane transport |
| jg8464 | IV | 1.0252 | 0.0096 | | transmembrane transporter activity; transmembrane transport |
| jg8466 | IV | 0.8350 | 0.0313 | | metal ion binding; intracellular cholesterol transport |

| | | | | | |
|---|---|---|---|---|---|
| jg8567 | IV | -0.9495 | 0.0436 | mutS protein homolog 4-like [Contarinia nasturtii] | ATP binding; mismatch repair; mismatched DNA binding; ATP-dependent DNA damage sensor activity |
| jg8577 | IV | -0.7761 | 0.0334 | | DNA binding; chromatin remodeling; ATP binding; ATP-dependent chromatin remodeler activity |
| jg8590 | IV | 1.7827 | 0.0037 | | L-threonine ammonia-lyase activity; threonine catabolic process |
| jg8659 | IV | 1.1588 | 0.0245 | | peroxidase activity; response to oxidative stress; heme binding |
| jg8780 | IV | 0.9081 | 0.0129 | | iron ion binding; oxidoreductase activity, acting on paired donors, with incorporation or reduction of molecular oxygen; monooxygenase activity; heme binding |
| jg8784 | IV | 1.0441 | 0.0424 | | NAD biosynthetic process; aspartate dehydrogenase activity; oxidoreductase activity; NADP binding |
| jg8785 | IV | 0.7631 | 0.0423 | | oxidoreductase activity; oxidoreductase activity, acting on the aldehyde or oxo group of donors, NAD or NADP as acceptor |
| jg8787 | IV | 1.0714 | 0.0424 | | 3-hydroxyanthranilate 3,4-dioxygenase activity; iron ion binding |
| jg8974 | IV | 0.8618 | 0.0420 | | metal ion transport; metal ion transmembrane transporter activity; transmembrane transport |
| jg9027 | IV | 1.7841 | 0.0297 | | phosphopantetheine binding; oxidoreductase activity; methyltransferase activity; 3-oxoacyl-[acyl-carrier-protein] synthase activity; fatty acid biosynthetic process; transferase activity; acyltransferase activity; biosynthetic process |
| jg9068 | IV | 0.5766 | 0.0496 | | oxidoreductase activity, acting on the CH-CH group of donors; flavin adenine dinucleotide binding; acyl-CoA dehydrogenase activity |
| jg9069 | IV | 0.9314 | 0.0320 | | oxidoreductase activity, acting on the CH-CH group of donors; flavin adenine dinucleotide binding; acyl-CoA dehydrogenase activity |
| jg9097 | IV | 0.7051 | 0.0458 | | transmembrane transport |
| jg9146 | IV | 1.0220 | 0.0388 | | serine-type endopeptidase activity; proteolysis |
| jg9166 | IV | 1.4459 | 0.0096 | | phosphopantetheine binding; oxidoreductase activity |
| jg9199 | IV | -0.8692 | 0.0498 | | nucleic acid binding; nucleotide binding; cellular metabolic process; DNA replication; DNA repair; 3'-5' DNA helicase activity; ATP binding; helicase activity; DNA recombination |
| jg9210 | IV | -0.8186 | 0.0489 | | protein binding; zinc ion binding |
| jg9293 | IV | 2.5979 | 0.0188 | Cell wall protein IFF6-like isoform X3 [Pieris rapae] | |
| jg9310 | IV | -0.8233 | 0.0411 | Histone H2B [Multiple species] | protein heterodimerization activity; DNA binding; structural constituent of chromatin |

| | | | | | |
|---|---|---|---|---|---|
| jg9317 | IV | -0.8340 | 0.0466 | | DNA binding; structural constituent of chromatin; protein heterodimerization activity |
| jg9320 | IV | -0.9088 | 0.0286 | Histone H2B [Multiple species] | protein heterodimerization activity; DNA binding; structural constituent of chromatin |
| jg9378 | IV | 1.1244 | 0.0499 | | zinc ion binding; hydrolase activity; catalytic activity; cytidine deaminase activity; cytidine deamination |
| jg9381 | IV | 1.3471 | 0.0466 | | zinc ion binding; hydrolase activity; cytidine deaminase activity; cytidine deamination; catalytic activity |
| jg9646 | IV | 2.7612 | 0.0134 | | NAD+ ADP-ribosyltransferase activity |
| jg9713 | IV | 0.8188 | 0.0489 | | protein binding |
| jg9714 | IV | 0.9703 | 0.0435 | | protein binding |
| jg9752 | IV | 1.2450 | 0.0480 | laccase-2 [Hermetia illucens] | copper ion binding; oxidoreductase activity |
| jg9993 | IV | -0.8728 | 0.0468 | | protein import into nucleus |
| jg10026 | IV | -0.8312 | 0.0374 | | DNA binding |
| jg10111 | IV | 1.1986 | 0.0194 | | serine-type endopeptidase activity; proteolysis |
| jg10233 | IV | 1.5591 | 0.0055 | | metallodipeptidase activity |
| jg10256 | IV | 0.5427 | 0.0441 | | 6-phosphofructo-2-kinase activity; ATP binding; fructose metabolic process; catalytic activity; fructose 2,6-bisphosphate metabolic process |
| jg10295 | IV | -0.7799 | 0.0493 | histone H1.2-like [Drosophila novamexicana] | protein heterodimerization activity; DNA binding; structural constituent of chromatin |
| jg10307 | IV | -0.8528 | 0.0477 | | DNA binding; structural constituent of chromatin; protein heterodimerization activity |
| jg10308 | IV | -0.7860 | 0.0497 | histone H1.2-like [Drosophila novamexicana] | DNA binding; structural constituent of chromatin; protein heterodimerization activity |
| jg10425 | IV | 4.8125 | 0.0155 | protein mesh [Folsomia candida] | |
| jg10429 | IV | 0.9247 | 0.0321 | facilitated trehalose transporter Tret1-2 homolog [Culex quinquefasciatus] | transmembrane transporter activity; transmembrane transport |
| jg10465 | IV | -0.7383 | 0.0381 | | protein binding |
| jg10562 | IV | 1.2426 | 0.0144 | glucosylceramidase [Anopheles darlingi] | glucosylceramidase activity; sphingolipid metabolic process |
| jg10658 | IV | 1.1218 | 0.0191 | lamin tail domain-containing protein [Myxococcus hansupus]; Cytochrome oxidase biogenesis protein Surf1, facilitates heme A insertion [Myxococcus hansupus] | |

| | | | | | |
|---|---|---|---|---|---|
| jg10780 | IV | 0.6894 | 0.0462 | | phosphatidylinositol phosphate biosynthetic process; inositol monophosphate 1-phosphatase activity |
| jg10847 | IV | 1.2879 | 0.0096 | | oxidoreductase activity |
| jg10929 | IV | 0.9023 | 0.0454 | | proteolysis; serine-type peptidase activity |
| jg11250 | IV | 0.6156 | 0.0362 | | proton-transporting ATPase activity, rotational mechanism |
| jg11310 | IV | 1.0782 | 0.0245 | | lipid transporter activity; lipid transport |
| jg11494 | IV | 0.9196 | 0.0367 | | protein transport |
| jg11513 | IV | 3.1652 | 0.0096 | | nucleic acid binding |
| jg11564 | IV | 2.0496 | 0.0096 | | catalytic activity; iron-sulfur cluster binding |
| jg11621 | IV | 0.8849 | 0.0158 | | oxidoreductase activity |
| jg11791 | IV | -0.6969 | 0.0489 | | protein binding; helicase activity; metal ion binding |
| jg11827 | IV | 0.8310 | 0.0381 | | antiporter activity; xenobiotic transmembrane transporter activity; transmembrane transport |
| jg11851 | IV | 0.8454 | 0.0381 | | transmembrane transporter activity; transmembrane transport |
| jg11854 | IV | 1.6035 | 0.0381 | | serine-type endopeptidase activity; proteolysis |
| jg11862 | IV | 3.3714 | 0.0126 | | nucleic acid binding |
| jg11867 | IV | 0.8869 | 0.0362 | phospholipase A2 inhibitor-like, partial [Contarinia nasturtii] | protein binding |
| jg11952 | IV | 1.9210 | 0.0393 | open rectifier potassium channel protein 1 [Culex pipiens pallens] | potassium channel activity; potassium ion transmembrane transport |
| jg12110 | IV | 0.6934 | 0.0466 | trans-1,2-dihydrobenzene-1,2-diol dehydrogenase-like [Hermetia illucens] | nucleotide binding |
| jg12152 | IV | 0.8365 | 0.0275 | | transmembrane transporter activity; transmembrane transport |
| jg12196 | IV | 0.6738 | 0.0466 | | G protein-coupled receptor activity; G protein-coupled receptor signaling pathway |
| jg12238 | IV | 0.8314 | 0.0436 | | SRP-dependent cotranslational protein targeting to membrane |
| jg12250 | IV | 1.2655 | 0.0242 | | carbohydrate transport; sugar transmembrane transporter activity; transmembrane transporter activity; transmembrane transport |
| jg12332 | IV | 0.7129 | 0.0303 | | ATP binding; protein kinase activity; protein phosphorylation; protein serine/threonine kinase activity; cGMP-dependent protein kinase activity |
| jg12429 | IV | 3.2894 | 0.0321 | PREDICTED: putative leucine-rich repeat-containing protein DDB_G0290503 [Rhagoletis zephyria] | |
| jg12508 | IV | 0.9940 | 0.0321 | | carbohydrate metabolic process |

| | | | | | |
|---|---|---|---|---|---|
| jg12511 | IV | 0.8968 | 0.0178 | collagen alpha-1(IV) chain isoform X2 [Contarinia nasturtii] | extracellular matrix structural constituent |
| jg12512 | IV | 1.0346 | 0.0169 | | extracellular matrix structural constituent |
| jg12514 | IV | 1.5114 | 0.0424 | UDPglucose 4-epimerase [Penicillium sp. RFL-2021a] | |
| jg12554 | IV | 0.6729 | 0.0222 | protein artichoke isoform X1 [Aedes aegypti] | protein binding |
| jg12582 | IV | 1.5634 | 0.0075 | | catalytic activity; biosynthetic process |
| jg12607 | IV | 1.4427 | 0.0165 | | nucleic acid binding; DNA integration |
| jg12671 | IV | 3.0146 | 0.0023 | | serine-type endopeptidase activity; proteolysis |
| jg12691 | IV | 1.2920 | 0.0088 | | catalytic activity |
| jg12717 | IV | 0.9394 | 0.0490 | | intracellular protein transport |
| jg12812 | IV | -0.8199 | 0.0188 | | protein binding |
| jg12847 | IV | 0.8019 | 0.0210 | | protein binding |
| jg13018 | IV | 3.2487 | 0.0096 | | nucleic acid binding |
| jg13156 | IV | 0.9617 | 0.0436 | | metallopeptidase activity; metalloendopeptidase activity; proteolysis; zinc ion binding; ATP binding; ATP-dependent chromatin remodeler activity |
| jg13178 | IV | 2.1619 | 0.0100 | | DNA-binding transcription factor activity; positive regulation of DNA-templated transcription; regulation of DNA-templated transcription |
| jg13370 | IV | 0.5063 | 0.0480 | | hydrolase activity |
| jg13488 | IV | 1.4735 | 0.0388 | | hydrolase activity, hydrolyzing O-glycosyl compounds; carbohydrate metabolic process; beta-galactosidase activity |
| jg13601 | IV | 1.6034 | 0.0367 | | nucleic acid binding; zinc ion binding |
| jg13663 | IV | -0.8273 | 0.0480 | | nucleic acid binding; nucleotide binding; DNA-directed DNA polymerase activity; DNA binding |
| jg13693 | IV | 2.4234 | 0.0275 | | proteolysis; metallodipeptidase activity |
| jg13847 | IV | 1.0433 | 0.0261 | | monooxygenase activity; iron ion binding; oxidoreductase activity, acting on paired donors, with incorporation or reduction of molecular oxygen; heme binding |
| jg13914 | IV | 1.3386 | 0.0441 | | transmembrane transport |
| jg14132 | IV | 1.0695 | 0.0291 | | acyltransferase activity; 3-oxoacyl-[acyl-carrier-protein] synthase activity; fatty acid biosynthetic process; biosynthetic process; oxidoreductase activity; transferase activity |
| jg14139 | IV | 1.3129 | 0.0096 | | serine-type endopeptidase activity; proteolysis; serine-type peptidase activity |

| | | | | | |
|---|---|---|---|---|---|
| jg14168 | IV | 0.7707 | 0.0417 | | sleep; regulation of synaptic transmission, cholinergic; positive regulation of voltage-gated potassium channel activity |
| jg14206 | IV | 0.7638 | 0.0380 | | chitin binding |
| jg14301 | IV | 0.8741 | 0.0251 | | sulfotransferase activity |
| jg14306 | IV | 0.7782 | 0.0296 | | transmembrane transporter activity; transmembrane transport |
| jg14482 | IV | 1.6059 | 0.0411 | | RNA nuclease activity; regulation of gene expression; mRNA base-pairing translational repressor activity; nucleic acid binding; RNA binding |
| jg14575 | IV | 2.1665 | 0.0266 | | serine-type endopeptidase activity; proteolysis |
| jg14648 | IV | -1.0597 | 0.0424 | | nucleic acid binding; zinc ion binding |
| jg14689 | IV | 1.2375 | 0.0096 | | transmembrane transporter activity; transmembrane transport |
| jg14743 | IV | 0.6721 | 0.0424 | | transmembrane transporter activity; transmembrane transport |
| jg14744 | IV | 1.0230 | 0.0110 | | serine-type endopeptidase activity; proteolysis |
| jg14892 | IV | 1.1771 | 0.0096 | | catalytic activity; carbohydrate metabolic process; carbohydrate binding; hydrolase activity, hydrolyzing O-glycosyl compounds |
| jg14893 | IV | 1.9125 | 0.0129 | | glucosylceramidase activity; sphingolipid metabolic process |
| jg14902 | IV | 1.2407 | 0.0474 | | phospholipid metabolic process |
| jg14964 | III | 0.6945 | 0.0381 | | cell adhesion; tissue regeneration |
| jg15333 | III | 5.6262 | 0.0062 | | DNA binding; protein dimerization activity |
| jg15477 | III | 1.7909 | 0.0126 | Regucalcin-like [Contarinia nasturtii] | |
| jg15739 | III | 1.1731 | 0.0381 | | transcription elongation by RNA polymerase II; histone modification |
| jg15842 | III | 0.9671 | 0.0180 | | monooxygenase activity; iron ion binding; oxidoreductase activity, acting on paired donors, with incorporation or reduction of molecular oxygen; heme binding |
| jg15900 | III | 0.9843 | 0.0381 | | chitin binding |
| jg15984 | III | 0.9160 | 0.0436 | | transposase activity; DNA transposition |
| jg16095 | III | 5.5663 | 0.0287 | | carbohydrate binding |
| jg16214 | III | 0.8986 | 0.0096 | | monooxygenase activity; iron ion binding; oxidoreductase activity, acting on paired donors, with incorporation or reduction of molecular oxygen; heme binding |
| jg16251 | III | 0.8126 | 0.0425 | | transmembrane transporter activity; transmembrane transport |
| jg16252 | III | 0.8554 | 0.0420 | | transmembrane transporter activity; transmembrane transport |
| jg16330 | III | 1.9593 | 0.0112 | Peptide transporter family 1-like isoform X1 [Contarinia nasturtii] | transmembrane transporter activity; transmembrane transport |

| | | | | | |
|---|---|---|---|---|---|
| jg16342 | III | -0.9511 | 0.0441 | | DNA binding |
| jg16446 | III | 1.5842 | 0.0488 | | NAD+-protein-arginine ADP-ribosyltransferase activity |
| jg16549 | III | 5.6398 | 0.0428 | | endopeptidase inhibitor activity |
| jg16652 | III | 1.3167 | 0.0309 | | transmembrane transporter activity; transmembrane transport; carbohydrate transport; sugar transmembrane transporter activity |
| jg16691 | III | 1.1269 | 0.0096 | | monoatomic cation transport; monoatomic cation transmembrane transporter activity; transmembrane transport; zinc ion transport |
| jg16695 | III | -0.7190 | 0.0362 | | protein binding |
| jg16788 | III | 0.9042 | 0.0316 | | N-acetylmuramoyl-L-alanine amidase activity; peptidoglycan catabolic process; monooxygenase activity; iron ion binding; oxidoreductase activity, acting on paired donors, with incorporation or reduction of molecular oxygen; heme binding; zinc ion binding |
| jg16913 | III | 1.1590 | 0.0321 | | monooxygenase activity; iron ion binding; oxidoreductase activity, acting on paired donors, with incorporation or reduction of molecular oxygen; heme binding |
| jg17068 | III | 1.6668 | 0.0393 | | serine-type endopeptidase inhibitor activity |
| jg17125 | III | 1.0802 | 0.0362 | Neuromedin-K receptor [Contarinia nasturtii] | G protein-coupled receptor activity; G protein-coupled receptor signaling pathway; neuropeptide Y receptor activity |
| jg17148 | III | 0.7591 | 0.0155 | | serine-type endopeptidase inhibitor activity |
| jg17186 | III | 1.6240 | 0.0472 | Cytidine deaminase-like protein [Aspergillus sergii] | catalytic activity |
| jg17393 | III | -0.8868 | 0.0366 | | U2 snRNA 3'-end processing |
| jg17556 | III | -0.6065 | 0.0468 | | regulation of transcription by RNA polymerase II; protein binding |
| jg17645 | III | -0.9462 | 0.0362 | E3 ubiquitin-protein ligase Bre1 [Contarinia nasturtii] | ubiquitin-protein transferase activity; histone monoubiquitination; metal ion binding |
| jg17751 | III | 1.6293 | 0.0144 | | transmembrane transporter activity; transmembrane transport |
| jg17955 | III | 0.7213 | 0.0377 | | amino acid catabolic process; carbohydrate catabolic process; L-fuconate dehydratase activity |
| jg17967 | III | 1.0528 | 0.0362 | | ATP binding; ABC-type transporter activity |
| jg18255 | III | 1.2046 | 0.0155 | Fork head domain-containing protein FD3 [Hermetia illucens] | DNA-binding transcription factor activity; regulation of DNA-templated transcription; sequence-specific DNA binding |
| jg18298 | III | -1.1527 | 0.0321 | | protein binding |
| jg18342 | III | 0.6225 | 0.0374 | | lipid metabolic process; phosphoric diester hydrolase activity |
| jg18375 | III | -0.5951 | 0.0377 | | mRNA splicing, via spliceosome; pre-mRNA 3'-splice site binding |

| | | | | | |
|---|---|---|---|---|---|
| jg18434 | III | 5.9566 | 0.0452 | | peroxidase activity; response to oxidative stress; heme binding |
| jg18451 | III | 0.9148 | 0.0381 | | transmembrane transporter activity; transmembrane transport; oligopeptide transport |
| jg18452 | III | 1.2961 | 0.0381 | | oxidoreductase activity |
| jg18465 | III | 1.0669 | 0.0131 | | protein binding |
| jg18592 | III | -1.4978 | 0.0452 | FAD-binding domain-containing protein [Conidiobolus coronatus NRRL 28638] | FAD binding; flavin adenine dinucleotide binding; oxidoreductase activity |
| jg18667 | III | 1.0518 | 0.0296 | Glutathione S-transferase [Bradysia odoriphaga] | glutathione transferase activity |
| jg18699 | III | 1.1782 | 0.0187 | | GTP binding; microtubule-based process; structural constituent of cytoskeleton |
| jg18706 | III | 0.8767 | 0.0096 | | protein binding; aminoacylase activity; amino acid metabolic process; hydrolase activity |
| jg18717 | III | 2.8673 | 0.0374 | Chymotrypsin-2-like [Drosophila subpulchrella] | serine-type endopeptidase activity; proteolysis |
| jg18732 | III | 1.8058 | 0.0424 | | protein binding |
| jg18753 | III | 2.3187 | 0.0057 | | polygalacturonase activity; carbohydrate metabolic process |
| jg18823 | III | 1.6444 | 0.0251 | | proteolysis; metallopeptidase activity; zinc ion binding |
| jg18845 | III | 1.7028 | 0.0096 | | dopamine beta-monooxygenase activity |
| jg18848 | III | 1.2625 | 0.0297 | | transmembrane transporter activity; transmembrane transport |
| jg18857 | III | 1.0395 | 0.0468 | | serine-type endopeptidase activity; proteolysis; cell adhesion; tissue regeneration |
| jg18859 | III | -1.0165 | 0.0424 | | protein binding; RNA binding; regulation of DNA-templated transcription; negative regulation of translation; translation repressor activity |
| jg18924 | III | 1.6719 | 0.0169 | Bombesin receptor subtype-3-like [Folsomia candida] | G protein-coupled receptor activity; G protein-coupled receptor signaling pathway |
| jg18975 | III | 0.5036 | 0.0489 | | proton-transporting ATPase activity, rotational mechanism; proton transmembrane transport; proton transmembrane transporter activity |
| jg19006 | III | -1.0087 | 0.0483 | | DNA replication |
| jg19015 | III | -1.0587 | 0.0498 | | DNA replication; nucleic acid binding; zinc ion binding; helicase activity; DNA recombination; ATP binding |
| jg19090 | III | 2.1729 | 0.0219 | Transglycosylase domain-containing protein [Clostridia bacterium] | |
| jg19292 | III | 2.8193 | 0.0105 | | RNA binding; adenosine deaminase activity; RNA processing; protein binding |
| jg19311 | III | 0.6301 | 0.0411 | | protein binding |
| jg19326 | III | -0.7514 | 0.0381 | | microtubule binding; microtubule anchoring |

| | | | | | |
|---|---|---|---|---|---|
| jg19352 | III | 1.2536 | 0.0297 | | transmembrane transporter activity; transmembrane transport |
| jg19427 | III | 2.0036 | 0.0023 | | transmembrane transporter activity; transmembrane transport |
| jg19485 | III | 0.7666 | 0.0436 | | hydrolase activity |
| jg19606 | III | 1.6262 | 0.0287 | Pyridoxal phosphate-dependent aminotransferase [Wolbachia endosymbiont of Ctenocephalides felis wCfeT] | biosynthetic process; pyridoxal phosphate binding; catalytic activity |
| jg19764 | III | 0.9135 | 0.0480 | | actin binding |
| jg19892 | III | -0.7831 | 0.0436 | | protein binding |
| jg20131 | III | 0.5485 | 0.0498 | | deaminase activity; adenosine deaminase activity; adenosine catabolic process |
| jg20137 | III | 1.2548 | 0.0321 | | protein binding |
| jg20169 | III | 0.9135 | 0.0381 | | FMN binding; oxidoreductase activity |
| jg20178 | III | 6.8677 | 0.0380 | | structural constituent of cell wall |
| jg20211 | III | 1.0184 | 0.0133 | | transmembrane transporter activity; transmembrane transport |
| jg20311 | III | 1.2634 | 0.0096 | | NAD+ ADP-ribosyltransferase activity |
| jg20434 | III | 0.8909 | 0.0442 | | methyltransferase activity |
| jg20558 | III | 0.9287 | 0.0195 | Macrophage-expressed gene 1 protein [Folsomia candida] | |
| jg20604 | III | 1.0294 | 0.0096 | | iron ion binding; oxidoreductase activity, acting on paired donors, with incorporation or reduction of molecular oxygen; monooxygenase activity; heme binding |
| jg20647 | III | 3.2065 | 0.0096 | | transmembrane transporter activity; transmembrane transport |
| jg20659 | III | 1.3540 | 0.0023 | | transmembrane transporter activity; transmembrane transport |
| jg20717 | III | 1.1085 | 0.0436 | | magnesium ion homeostasis |
| jg20825 | III | 0.8879 | 0.0240 | | aminoacylase activity; amino acid metabolic process; hydrolase activity |
| jg20835 | III | 1.0939 | 0.0101 | | symporter activity |
| jg20845 | III | 1.1949 | 0.0362 | | oxidoreductase activity |

**Table S5.** Results of GO-term enrichment analysis on significantly differentially expressed genes between gynogenic and androgenic females in somatic non-reproductive tissue. The p-value is obtained with a Classic Fisher test.

| GO:ID | Term | Category | No. of genes with this GO term | No. of significantly expressed genes | Expected no. of significantly | p-value |
|---|---|---|---|---|---|---|

| | | | | with this GO term | expressed genes with this GO term | |
|---|---|---|---|---|---|---|
| GO:0055085 | transmembrane transport | Biological Process | 357 | 34 | 11.01 | 6.40E-09 |
| GO:0006810 | transport | Biological Process | 695 | 44 | 21.44 | 5.70E-06 |
| GO:0051234 | establishment of localization | Biological Process | 699 | 44 | 21.56 | 6.50E-06 |
| GO:0051179 | localization | Biological Process | 707 | 44 | 21.81 | 8.70E-06 |
| GO:0009092 | homoserine metabolic process | Biological Process | 4 | 2 | 0.12 | 0.0055 |
| GO:0019346 | transsulfuration | Biological Process | 4 | 2 | 0.12 | 0.0055 |
| GO:0050667 | homocysteine metabolic process | Biological Process | 4 | 2 | 0.12 | 0.0055 |
| GO:0006534 | cysteine metabolic process | Biological Process | 8 | 2 | 0.25 | 0.0235 |
| GO:0019318 | hexose metabolic process | Biological Process | 9 | 2 | 0.28 | 0.0296 |
| GO:0000103 | sulfate assimilation | Biological Process | 1 | 1 | 0.03 | 0.0308 |
| GO:0006000 | fructose metabolic process | Biological Process | 1 | 1 | 0.03 | 0.0308 |
| GO:0034453 | microtubule anchoring | Biological Process | 1 | 1 | 0.03 | 0.0308 |
| GO:0034474 | U2 snRNA 3'-end processing | Biological Process | 1 | 1 | 0.03 | 0.0308 |
| GO:0000096 | sulfur amino acid metabolic process | Biological Process | 10 | 2 | 0.31 | 0.0363 |
| GO:0009069 | serine family amino acid metabolic process | Biological Process | 11 | 2 | 0.34 | 0.0434 |
| GO:0098662 | inorganic cation transmembrane transport | Biological Process | 63 | 5 | 1.94 | 0.0447 |
| GO:0006790 | sulfur compound metabolic process | Biological Process | 27 | 3 | 0.83 | 0.0493 |
| GO:0098660 | inorganic ion transmembrane | Biological Process | 65 | 5 | 2 | 0.05 |

| | | | | | | |
|---|---|---|---|---|---|---|
| | transport | | | | | |
| GO:0005581 | collagen trimer | Cellular Component | 2 | 2 | 0.05 | 0.00065 |
| GO:0005618 | cell wall | Cellular Component | 1 | 1 | 0.03 | 0.02557 |
| GO:0005784 | Sec61 translocon complex | Cellular Component | 1 | 1 | 0.03 | 0.02557 |
| GO:0009277 | fungal-type cell wall | Cellular Component | 1 | 1 | 0.03 | 0.02557 |
| GO:0030289 | protein phosphatase 4 complex | Cellular Component | 1 | 1 | 0.03 | 0.02557 |
| GO:0071256 | translocon complex | Cellular Component | 1 | 1 | 0.03 | 0.02557 |
| GO:0022857 | transmembrane transporter activity | Molecular Function | 533 | 48 | 15.24 | 3.90E-12 |
| GO:0005215 | transporter activity | Molecular Function | 560 | 49 | 16.01 | 6.60E-12 |
| GO:0031177 | phosphopantetheine binding | Molecular Function | 2 | 2 | 0.06 | 0.00082 |
| GO:0042910 | xenobiotic transmembrane transporter activity | Molecular Function | 2 | 2 | 0.06 | 0.00082 |
| GO:0072341 | modified amino acid binding | Molecular Function | 2 | 2 | 0.06 | 0.00082 |
| GO:0005201 | extracellular matrix structural constituent | Molecular Function | 3 | 2 | 0.09 | 0.0024 |
| GO:0005506 | iron ion binding | Molecular Function | 166 | 12 | 4.74 | 0.00307 |
| GO:0019842 | vitamin binding | Molecular Function | 38 | 5 | 1.09 | 0.00434 |
| GO:0004348 | glucosylceramidase activity | Molecular Function | 5 | 2 | 0.14 | 0.00771 |
| GO:0003950 | NAD+ ADP-ribosyltransferase activity | Molecular Function | 6 | 2 | 0.17 | 0.01134 |
| GO:0022890 | inorganic cation transmembrane transporter activity | Molecular Function | 129 | 9 | 3.69 | 0.01202 |
| GO:0004867 | serine-type endopeptidase | Molecular Function | 18 | 3 | 0.51 | 0.01379 |

| | | | | | | |
|---|---|---|---|---|---|---|
| | inhibitor activity | | | | | |
| GO:0016763 | pentosyltransferase activity | Molecular Function | 18 | 3 | 0.51 | 0.01379 |
| GO:0070573 | metallodipeptidase activity | Molecular Function | 7 | 2 | 0.2 | 0.01558 |
| GO:0008324 | monoatomic cation transmembrane transporter activity | Molecular Function | 136 | 9 | 3.89 | 0.0165 |
| GO:0016705 | oxidoreductase activity, acting on paired donors, with incorporation or reduction of molecular oxygen | Molecular Function | 184 | 11 | 5.26 | 0.01727 |
| GO:0046982 | protein heterodimerization activity | Molecular Function | 21 | 3 | 0.6 | 0.0211 |
| GO:0016491 | oxidoreductase activity | Molecular Function | 591 | 26 | 16.89 | 0.02123 |
| GO:0015318 | inorganic molecular entity transmembrane transporter activity | Molecular Function | 144 | 9 | 4.12 | 0.02298 |
| GO:0015297 | antiporter activity | Molecular Function | 9 | 2 | 0.26 | 0.02572 |
| GO:0019239 | deaminase activity | Molecular Function | 9 | 2 | 0.26 | 0.02572 |
| GO:0022804 | active transmembrane transporter activity | Molecular Function | 59 | 5 | 1.69 | 0.0266 |
| GO:0016810 | hydrolase activity, acting on carbon-nitrogen (but not peptide) bonds | Molecular Function | 40 | 4 | 1.14 | 0.02687 |
| GO:0015078 | proton transmembrane transporter activity | Molecular Function | 23 | 3 | 0.66 | 0.02695 |
| GO:0015291 | secondary active transmembrane transporter activity | Molecular Function | 23 | 3 | 0.66 | 0.02695 |
| GO:0000334 | 3-hydroxyanthranilate 3,4-dioxygenase activity | Molecular Function | 1 | 1 | 0.03 | 0.02858 |

| | | | | | | |
|---|---|---|---|---|---|---|
| GO:0003873 | 6-phosphofructo-2-kinase activity | Molecular Function | 1 | 1 | 0.03 | 0.02858 |
| GO:0004779 | sulfate adenylyltransferase activity | Molecular Function | 1 | 1 | 0.03 | 0.02858 |
| GO:0004781 | sulfate adenylyltransferase (ATP) activity | Molecular Function | 1 | 1 | 0.03 | 0.02858 |
| GO:0005199 | structural constituent of cell wall | Molecular Function | 1 | 1 | 0.03 | 0.02858 |
| GO:0030628 | pre-mRNA 3'-splice site binding | Molecular Function | 1 | 1 | 0.03 | 0.02858 |
| GO:0033735 | aspartate dehydrogenase activity | Molecular Function | 1 | 1 | 0.03 | 0.02858 |
| GO:0036002 | pre-mRNA binding | Molecular Function | 1 | 1 | 0.03 | 0.02858 |
| GO:0030170 | pyridoxal phosphate binding | Molecular Function | 24 | 3 | 0.69 | 0.03016 |
| GO:0070279 | vitamin B6 binding | Molecular Function | 24 | 3 | 0.69 | 0.03016 |
| GO:0016805 | dipeptidase activity | Molecular Function | 10 | 2 | 0.29 | 0.03155 |
| GO:0030527 | structural constituent of chromatin | Molecular Function | 25 | 3 | 0.71 | 0.03356 |
| GO:0033218 | amide binding | Molecular Function | 11 | 2 | 0.31 | 0.03784 |
| GO:0004866 | endopeptidase inhibitor activity | Molecular Function | 28 | 3 | 0.8 | 0.0449 |
| GO:0061135 | endopeptidase regulator activity | Molecular Function | 28 | 3 | 0.8 | 0.0449 |

#### Supplementary material 5: For maternal deposit into eggs, top autosomal differentially expressed genes and top genes with biased X' allele-biased expression, with functional annotation and GO term enrichment analysis

**Table S9.** Functional information for genes found to have significant X' allele-biased expression in the maternal deposit of female eggs. The Gene ID for the inversion allele and the X allele are both given. Log fold change (logFC) and the p-value is given to 4 decimal places or 3 significant figures. Column 5 and 6 contains BLAST results to the entire Non-redundant (NR) Protein database, excluding *Bradysia coprophila*, for the inversion allele and the X allele respectively. Column 7 and 8 contains GO terms that result from a comprehensive InterProScan to obtain protein domain-level predictions and other protein/peptide/domain motif analyses to predict functional information, for the inversion allele and the X allele respectively. Many hits to the NCBI database were to hypothetical/uncharacterised proteins, and these were left out. Any genes with significant allele-biased expression but have no hits to the NCBI database/hits only to hypothetical proteins, and no GO term results from the InterProScan, are not shown in this table. Genes where the two alleles have NCBI to different proteins, or are predicted to have different GO term annotations, is shown in bold.

| Gene ID (Inv) | Gene ID (X) | logFC (4 d.p.) | p-value (4 d.p. or 3 s.f.) | NCBI best-match protein (Inv) | NCBI best-match protein (X) | InterProScan GO Term function (Inv) | InterProScan GO Term function (X) |
|---|---|---|---|---|---|---|---|
| **jg20915** | **jg4463** | **4.8690** | / | | | **transmembrane transport** | |
| jg20937 | jg4541 | 0.2673 | 4.08E-07 | | | protein kinase activity; ATP binding; protein phosphorylation | protein kinase activity; ATP binding; protein phosphorylation |
| jg20999 | jg4597 | 6.0140 | 2.51E-06 | | | protein binding | protein binding |
| jg21021 | jg4624 | 0.4445 | 7.20E-10 | | | dephosphorylation; phosphatase activity; protein tyrosine/serine/threonine phosphatase activity; protein dephosphorylation | dephosphorylation; phosphatase activity; protein dephosphorylation; protein tyrosine/serine/threonine phosphatase activity |
| jg21104 | jg4717 | 0.1899 | 1.10E-02 | | | DNA-binding transcription factor activity; regulation of DNA-templated transcription; signal transduction; DNA binding; growth hormone receptor signaling pathway via JAK-STAT | DNA-binding transcription factor activity; regulation of DNA-templated transcription; signal transduction; DNA binding; growth hormone receptor signaling pathway via JAK-STAT |
| jg21154 | jg4760 | 0.0986 | 5.30E-03 | | | holocytochrome-c synthase activity | holocytochrome-c synthase activity |
| **jg21234** | **jg4841** | **3.3378** | **5.26E-03** | | | **cytoskeletal motor activity; ATP binding** | |

| | | | | | | | |
|---|---|---|---|---|---|---|---|
| jg21248 | jg4857 | 0.3023 | 1.24E-09 | Poly [ADP-ribose] polymerase 1 [Folsomia candida] | poly [ADP-ribose] polymerase isoform X2 [Folsomia candida] | NAD+ ADP-ribosyltransferase activity | NAD+ ADP-ribosyltransferase activity |
| jg21281 | jg4892 | 1.3587 | 5.27E-30 | | | tail-anchored membrane protein insertion into ER membrane | tail-anchored membrane protein insertion into ER membrane |
| **jg21349** | **jg4971** | **8.8846** | **9.80E-36** | | | **mRNA processing; hydrolase activity, acting on ester bonds; hemolysis in another organism** | **hemolysis in another organism** |
| jg21361 | jg4980 | 3.5039 | 2.67E-06 | | | hydrolase activity, hydrolyzing O-glycosyl compounds; carbohydrate metabolic process | hydrolase activity, hydrolyzing O-glycosyl compounds; carbohydrate metabolic process |
| jg21363 | jg4982 | 0.9638 | 4.39E-07 | | | MAP kinase activity; ATP binding; protein phosphorylation; protein kinase activity | MAP kinase activity; ATP binding; protein phosphorylation; protein kinase activity |
| **jg21433** | **jg5062** | **2.2800** | **4.63E-107** | | | **enzyme inhibitor activity; ionotropic glutamate receptor activity; serine-type endopeptidase inhibitor activity; ligand-gated monoatomic ion channel activity; cholesterol metabolic process; oxidoreductase activity** | **peptidase inhibitor activity; enzyme inhibitor activity; serine-type endopeptidase inhibitor activity** |
| jg21484 | jg5116 | 0.3302 | 1.86E-02 | | | serine-type endopeptidase activity; proteolysis | serine-type endopeptidase activity; proteolysis |
| jg21488 | jg5121 | 0.2369 | 5.21E-05 | | | damaged DNA binding; DNA repair | damaged DNA binding; DNA repair |
| jg21502 | jg5141 | 0.3069 | 7.39E-03 | | | cytosolic ribosome assembly; ribosome biogenesis | cytosolic ribosome assembly; ribosome biogenesis |
| jg21546 | jg5180 | 0.7296 | 9.91E-14 | | | calcium ion binding; transmembrane transport | transmembrane transport; calcium ion binding |
| **jg21553** | **jg5189** | **0.9609** | **1.60E-87** | | | **pseudouridine synthesis; RNA binding; RNA modification; pseudouridine synthase activity** | **pseudouridine synthesis; RNA binding; RNA modification; pseudouridine synthase activity; tRNA binding; aminoacyl-tRNA ligase activity; ATP binding; tRNA aminoacylation** |
| **jg21665** | **jg5288** | **0.4399** | **3.67E-09** | | | **hydrolase activity, acting on ester bonds; lipid metabolic process** | **lipid metabolic process** |

| | | | | | | | |
|---|---|---|---|---|---|---|---|
| jg21695 | jg5311 | 0.0083 | 4.61E-04 | | | helicase activity; nucleic acid binding; ATP binding | nucleic acid binding; ATP binding; helicase activity |
| jg21703 | jg5321 | 3.9077 | 9.18E-196 | | | glycylpeptide N-tetradecanoyltransferase activity; N-terminal protein myristoylation | glycylpeptide N-tetradecanoyltransferase activity; N-terminal protein myristoylation |
| **jg21709** | **jg5328** | **10.9303** | **/** | | | **nucleic acid binding; zinc ion binding** | |
| jg21725 | jg5340 | 1.9924 | 4.81E-02 | | | carbonate dehydratase activity; zinc ion binding | carbonate dehydratase activity; zinc ion binding |
| jg21780 | jg5400 | 5.2172 | 4.35E-05 | | | peptidase inhibitor activity | peptidase inhibitor activity |
| **jg21810** | **jg5425** | **3.4565** | **/** | | | **protein serine/threonine kinase activity; protein phosphorylation** | |
| **jg21829** | **jg6453** | **3.2361** | **8.45E-12** | | | **protein binding** | **sleep; regulation of synaptic transmission, cholinergic; positive regulation of voltage-gated potassium channel activity** |
| jg21847 | jg5469 | 4.8803 | 1.04E-37 | | | oxidoreductase activity; flavin adenine dinucleotide binding; protein dimerization activity; ubiquitin-protein transferase activity; protein ubiquitination | oxidoreductase activity; flavin adenine dinucleotide binding; protein dimerization activity |
| jg21922 | jg5532 | 0.3659 | 1.71E-05 | | | protein binding | protein binding |
| **jg21935** | **jg5548** | **3.1456** | **/** | | | **serine-type endopeptidase activity; proteolysis** | **olfactory receptor activity; odorant binding; sensory perception of smell; serine-type endopeptidase activity; proteolysis** |
| jg21939 | jg5553 | 0.2268 | 2.30E-02 | | | iron ion transport; intracellular iron ion homeostasis; ferric iron binding | iron ion transport; intracellular iron ion homeostasis; ferric iron binding |
| jg21960 | jg5577 | 1.6040 | 1.26E-16 | microsomal glutathione S-transferase 1-like isoform X2 [Lucilia cuprina] | microsomal glutathione S-transferase 1-like isoform X2 [Lucilia cuprina] | | |
| **jg21966** | **jg5586** | **3.3008** | **2.52E-25** | | | | **ATP binding; ATP-dependent protein folding chaperone** |
| jg21993 | jg5613 | -0.5214 | 3.98E-15 | | | zinc ion binding; ubiquitin protein ligase activity; ubiquitin-dependent protein catabolic process via the N-end rule pathway | zinc ion binding; ubiquitin protein ligase activity; ubiquitin-dependent protein catabolic process via the N-end rule pathway |

| | | | | | | | |
|---|---|---|---|---|---|---|---|
| jg21994 | jg5614 | 0.3313 | 1.90E-02 | | | copper ion binding; copper chaperone activity | copper ion binding; copper chaperone activity |
| jg21995 | jg5615 | 0.9584 | 1.32E-27 | | | NADH dehydrogenase (ubiquinone) activity; ATP synthesis coupled electron transport; oxidoreductase activity; iron-sulfur cluster binding; oxidoreductase activity, acting on NAD(P)H | oxidoreductase activity; NADH dehydrogenase (ubiquinone) activity; ATP synthesis coupled electron transport; iron-sulfur cluster binding; oxidoreductase activity, acting on NAD(P)H |
| jg22056 | jg5675 | 0.8366 | 1.76E-09 | | | protein binding | protein binding |
| **jg22071** | **jg5692** | **-1.9167** | **1.01E-07** | | | | **ATP binding; hydrolase activity** |
| jg22211 | jg5844 | 0.2306 | 2.79E-02 | | | protein metabolic process; manganese ion binding; metalloaminopeptidase activity; proteolysis | proteolysis; metalloaminopeptidase activity; protein metabolic process; manganese ion binding |
| jg22233 | jg5863 | 0.3215 | 7.90E-06 | | | hydrolase activity, acting on carbon-nitrogen (but not peptide) bonds; guanine catabolic process; zinc ion binding; guanine deaminase activity; hydrolase activity | hydrolase activity, acting on carbon-nitrogen (but not peptide) bonds; hydrolase activity; guanine catabolic process; zinc ion binding; guanine deaminase activity |
| jg22260 | jg5889 | 1.8744 | 9.20E-26 | | | protein binding | protein binding |
| jg22273 | jg5899 | 9.2182 | 4.62E-27 | | | cilium movement; outer dynein arm assembly | cilium movement; outer dynein arm assembly |
| jg22282 | jg5911 | 0.5221 | 6.07E-05 | | | protein binding; structural molecule activity; retrograde vesicle-mediated transport, Golgi to endoplasmic reticulum | protein binding; structural molecule activity; retrograde vesicle-mediated transport, Golgi to endoplasmic reticulum |
| **jg22288** | **jg5916** | **8.8483** | / | | **cytochrome P450 [Chironomus tentans]** | **monooxygenase activity; iron ion binding; oxidoreductase activity, acting on paired donors, with incorporation or reduction of molecular oxygen; heme binding; calcium ion binding; enzyme regulator activity** | **monooxygenase activity; iron ion binding; oxidoreductase activity, acting on paired donors, with incorporation or reduction of molecular oxygen; heme binding** |
| jg22319 | jg5949 | 3.8207 | / | | | metalloendopeptidase activity; proteolysis; metallopeptidase activity; zinc ion binding | metalloendopeptidase activity; proteolysis; metallopeptidase activity; zinc ion binding |

| | | | | | | | |
|---|---|---|---|---|---|---|---|
| jg22377 | jg6019 | 0.1128 | 2.98E-05 | | | calcium ion binding; mitochondrial calcium ion transmembrane transport | calcium ion binding; mitochondrial calcium ion transmembrane transport |
| jg22394 | jg6032 | 1.2685 | 6.70E-36 | | | peptidyl-prolyl cis-trans isomerase activity | peptidyl-prolyl cis-trans isomerase activity |
| jg22464 | jg6098 | 0.4078 | 5.85E-06 | | | proteolysis; metalloexopeptidase activity | proteolysis; metalloexopeptidase activity |
| jg22468 | jg6102 | 0.1892 | 3.84E-03 | | | transcription coregulator activity; regulation of transcription by RNA polymerase II | transcription coregulator activity; regulation of transcription by RNA polymerase II |
| jg22477 | jg6111 | 1.1215 | 1.88E-23 | | | metallopeptidase activity; zinc ion binding; proteolysis | proteolysis; metallopeptidase activity; zinc ion binding |
| **jg22505** | **jg6138** | **2.5923** | **2.50E-116** | | | **transaminase activity; pyridoxal phosphate binding; catalytic activity** | |
| jg22507 | jg6139 | 4.8858 | / | | | calcium ion binding | calcium ion binding |
| jg22531 | jg6164 | 0.6690 | 4.98E-10 | | | catalytic activity | catalytic activity |
| **jg22540** | **jg6170** | **7.6434** | **/** | **odorant receptor 57 [Bradysia odoriphaga]** | **odorant receptor 57 [Bradysia odoriphaga]** | | **olfactory receptor activity; odorant binding; sensory perception of smell** |
| **jg22733** | **jg6343** | **10.6978** | **9.80E-18** | | | **magnesium ion binding; tRNA modification; tRNA guanylyltransferase activity** | |
| jg22756 | jg6367 | 0.3179 | 2.31E-03 | | | protein binding | protein binding |
| jg22775 | jg6384 | 0.2328 | 2.64E-04 | | | protein binding | protein binding |
| jg22782 | jg6390 | 2.0075 | 1.08E-03 | | | oxidoreductase activity, acting on the CH-CH group of donors; acyl-CoA oxidase activity; fatty acid metabolic process; FAD binding | acyl-CoA oxidase activity; fatty acid metabolic process; FAD binding; oxidoreductase activity, acting on the CH-CH group of donors; fatty acid beta-oxidation |
| **jg22783** | **jg6391** | **0.9187** | **7.01E-237** | **neuroglian-like isoform X2 [Contarinia nasturtii]** | | | |
| jg22787 | jg6394 | 0.6224 | 4.53E-10 | | | heat shock protein binding; unfolded protein binding; protein folding; Hsp70 protein binding | heat shock protein binding; unfolded protein binding; protein folding; Hsp70 protein binding |
| jg22876 | jg6496 | 0.3467 | 2.21E-06 | | | catalytic activity | catalytic activity |

| | | | | | | | |
|---|---|---|---|---|---|---|---|
| jg22919 | jg6541 | 1.4360 | 1.46E-63 | | | kinase activity; NAD+ kinase activity | kinase activity; NAD+ kinase activity |
| jg22925 | jg6550 | 0.2913 | 3.26E-02 | | | small GTPase binding | small GTPase binding |
| jg22927 | jg6551 | 1.3282 | 3.26E-09 | | | protein binding | protein binding |
| jg22946 | jg6560 | 3.2172 | 4.97E-07 | | | GTPase activity; GTP binding | GTPase activity; GTP binding |
| jg22972 | jg6590 | 3.1739 | / | | | carbohydrate metabolic process; hydrolase activity, acting on carbon-nitrogen (but not peptide) bonds | carbohydrate metabolic process; hydrolase activity, acting on carbon-nitrogen (but not peptide) bonds |
| jg23000 | jg6620 | 0.4446 | 1.08E-07 | | | metallopeptidase activity; metalloendopeptidase activity; proteolysis | metalloendopeptidase activity; proteolysis; metallopeptidase activity |
| jg23002 | jg6622 | 0.0084 | 4.27E-02 | | | metallopeptidase activity; metalloendopeptidase activity; proteolysis | metallopeptidase activity; metalloendopeptidase activity; proteolysis |
| jg23004 | jg6624 | 0.8735 | 3.41E-03 | | | protein binding | protein binding |
| jg23013 | jg6633 | 0.2698 | 5.73E-03 | cytochrome P450 4V2 isoform X1 [Pipistrellus kuhlii] | cytochrome P450 4V2 isoform X1 [Pipistrellus kuhlii] | iron ion binding; oxidoreductase activity, acting on paired donors, with incorporation or reduction of molecular oxygen; monooxygenase activity; heme binding | iron ion binding; oxidoreductase activity, acting on paired donors, with incorporation or reduction of molecular oxygen; monooxygenase activity; heme binding |
| jg23050 | jg6668 | 0.5079 | 3.65E-03 | zinc finger protein OZF-like isoform X1 [Photinus pyralis] | zinc finger protein 879 isoform X1 [Drosophila santomea] | zinc ion binding | zinc ion binding |
| jg23071 | jg6687 | -0.0578 | 1.30E-06 | | | acetyltransferase activity; N-acetyltransferase activity | acetyltransferase activity; N-acetyltransferase activity |
| jg23106 | jg6715 | 0.2129 | 2.59E-03 | | | protein binding | protein binding |
| jg23157 | jg6760 | 0.2897 | 1.85E-05 | | | ATP binding; protein kinase activity; protein phosphorylation | protein kinase activity; protein phosphorylation; ATP binding |
| jg23165 | jg6770 | 0.9609 | 5.83E-21 | | | hydrolase activity, acting on acid anhydrides, in phosphorus-containing anhydrides; metal ion binding | hydrolase activity, acting on acid anhydrides, in phosphorus-containing anhydrides; metal ion binding |
| jg23186 | jg6791 | 5.0061 | / | | | chitin binding | chitin binding |
| **jg23200** | **jg6804** | **-1.3270** | **5.28E-05** | | | **signal transduction; toll-like receptor signaling pathway; transmembrane signaling** | **signal transduction; protein binding** |

| | | | | | | | |
|---|---|---|---|---|---|---|---|
| | | | | | | **receptor activity; immune response** | |
| jg23204 | jg6808 | 2.0329 | 1.48E-11 | | | glucosylceramidase activity; sphingolipid metabolic process | glucosylceramidase activity; sphingolipid metabolic process |
| jg23210 | jg6814 | 1.9217 | 1.36E-11 | | | protein binding; calcium channel activity; calcium ion transmembrane transport; monoatomic ion channel activity; monoatomic ion transport; transmembrane transport | protein binding; calcium channel activity; calcium ion transmembrane transport; monoatomic ion channel activity; monoatomic ion transport; transmembrane transport |
| jg23213 | jg6816 | 0.1950 | 4.30E-04 | | | magnesium ion binding; ribose phosphate diphosphokinase activity; nucleotide biosynthetic process | magnesium ion binding; ribose phosphate diphosphokinase activity; nucleotide biosynthetic process |
| jg23228 | jg6830 | 9.6003 | / | | | ATP binding; transmembrane transport; ABC-type transporter activity | ABC-type transporter activity; ATP binding; transmembrane transport |
| **jg23237** | **jg6837** | **1.2584** | **4.30E-10** | | | **serine-type endopeptidase activity; proteolysis** | |
| jg23324 | jg6911 | 1.9874 | 1.95E-04 | | | transmembrane transporter activity; transmembrane transport | transmembrane transporter activity; transmembrane transport |
| **jg23325** | **jg6912** | **-0.0731** | **1.30E-02** | | | | **zinc ion binding; Notch signaling pathway; protein ubiquitination** |
| jg23329 | jg6917 | 2.5045 | 1.89E-03 | | | alkylmercury lyase activity; organomercury catabolic process; iron ion binding; lipid biosynthetic process; oxidoreductase activity | iron ion binding; lipid biosynthetic process; oxidoreductase activity |
| jg23338 | jg6926 | 0.2822 | 5.19E-04 | | | mannosyl-glycoprotein endo-beta-N-acetylglucosaminidase activity | mannosyl-glycoprotein endo-beta-N-acetylglucosaminidase activity |
| jg23401 | jg6995 | 0.1808 | 3.42E-02 | | | protein-containing complex assembly | protein-containing complex assembly |
| jg23410 | jg7006 | 1.1830 | 2.68E-58 | | | RNA binding; nucleic acid binding | RNA binding; nucleic acid binding |
| jg23418 | jg7013 | 2.2558 | 3.29E-28 | | | catalytic activity | catalytic activity |
| jg23438 | jg7037 | 4.4044 | 3.60E-05 | | | protein binding; regulation of kainate selective glutamate receptor activity | protein binding; regulation of kainate selective glutamate receptor activity |
| **jg23443** | **jg7043** | **11.6049** | **/** | | | **hydrolase activity, hydrolyzing O-glycosyl compounds;** | |

| | | | | | | | |
|---|---|---|---|---|---|---|---|
| | | | | | | **carbohydrate metabolic process; protein binding** | |
| jg23448 | jg7055 | 1.9524 | 1.89E-55 | | | protein binding; transcription elongation by RNA polymerase II | transcription elongation by RNA polymerase II; protein binding |
| jg23462 | jg7075 | 11.7515 | / | DNA damage-binding protein 1 [Culex pipiens pallens] | PREDICTED: echinoderm microtubule-associated protein-like CG42247 [Bactrocera latifrons] | intracellular signal transduction; nucleic acid binding; protein binding; DNA repair | intracellular signal transduction |
| jg23486 | jg7096 | 2.8465 | 4.55E-85 | | | protein binding | protein binding |
| jg23490 | jg7100 | 1.2834 | 3.12E-33 | PREDICTED: delta(24)-sterol reductase-like [Branchiostoma belcheri] | PREDICTED: delta(24)-sterol reductase-like [Branchiostoma belcheri] | FAD binding; flavin adenine dinucleotide binding | flavin adenine dinucleotide binding; FAD binding |
| **jg23498** | **jg7108** | **0.1057** | **1.57E-43** | | | | **oxidoreductase activity** |
| jg23511 | jg7119 | -0.7593 | 9.68E-03 | | | carbonate dehydratase activity; zinc ion binding | carbonate dehydratase activity; zinc ion binding |
| jg23528 | jg7140 | 12.2831 | / | diacylglycerol kinase theta isoform X6 [Contarinia nasturtii] | diacylglycerol kinase theta isoforms [Contarinia nasturtii] | ATP-dependent diacylglycerol kinase activity; signal transduction; protein kinase C-activating G protein-coupled receptor signaling pathway; kinase activity; RNA binding; intracellular protein transport; vesicle-mediated transport; NAD+ kinase activity; clathrin adaptor activity; nucleic acid binding | nucleic acid binding; ATP-dependent diacylglycerol kinase activity; signal transduction; RNA binding; NAD+ kinase activity; kinase activity; protein kinase C-activating G protein-coupled receptor signaling pathway |
| jg23535 | jg7154 | 1.1866 | 1.28E-09 | | | DNA repair | DNA repair |
| jg23555 | jg7179 | -0.1744 | 5.83E-28 | | | DNA binding; DNA-binding transcription factor activity, RNA polymerase II-specific; regulation of DNA-templated transcription | DNA binding; DNA-binding transcription factor activity, RNA polymerase II-specific; regulation of DNA-templated transcription |
| jg23556 | jg7181 | 4.3344 | / | | | nucleic acid binding; zinc ion binding; monooxygenase activity; iron ion binding; oxidoreductase activity, acting on paired donors, with incorporation or reduction of molecular oxygen; heme binding | monooxygenase activity; iron ion binding; oxidoreductase activity, acting on paired donors, with incorporation or reduction of molecular oxygen; heme binding |

| | | | | | | | |
|---|---|---|---|---|---|---|---|
| **jg23569** | **jg7191** | **-1.2357** | **1.06E-40** | | | **DNA-binding transcription factor activity; regulation of DNA-templated transcription; sequence-specific DNA binding** | **DNA-binding transcription factor activity; regulation of DNA-templated transcription; sequence-specific DNA binding; protein binding; intracellular signal transduction** |
| jg23610 | jg6776 | 10.1987 | / | deleted in malignant brain tumors 1 protein [Hermetia illucens] | deleted in malignant brain tumors 1 protein [Hermetia illucens] | | |
| jg23620 | jg7238 | 0.5563 | 6.44E-03 | | | RNA binding; GTPase activity; GTP binding; intracellular signal transduction; nucleic acid binding | nucleic acid binding; RNA binding |
| jg23633 | jg7252 | 0.5176 | 1.80E-09 | | | calcium ion binding | calcium ion binding |
| jg23658 | jg7273 | 4.2172 | / | 3-hydroxyisobutyryl-CoA hydrolase, mitochondrial-like [Anopheles albimanus] | 3-hydroxyisobutyryl-CoA hydrolase [Tropilaelaps mercedesae] | | |
| jg23690 | jg7309 | 4.9852 | / | | | monooxygenase activity; iron ion binding; oxidoreductase activity, acting on paired donors, with incorporation or reduction of molecular oxygen; heme binding | monooxygenase activity; iron ion binding; oxidoreductase activity, acting on paired donors, with incorporation or reduction of molecular oxygen; heme binding |
| jg23700 | jg7323 | -0.3304 | 7.60E-12 | | | regulation of DNA-templated transcription | regulation of DNA-templated transcription |
| jg23708 | jg7333 | 4.1960 | 4.34E-50 | | | nucleic acid binding | nucleic acid binding |
| **jg23710** | **jg7335** | **4.4212** | **2.60E-104** | | | **transcription elongation by RNA polymerase II** | |
| jg23718 | jg7341 | 0.7857 | 8.89E-03 | | | protein binding; monoatomic ion channel activity; monoatomic ion transport; transmembrane transport | protein binding; monoatomic ion channel activity; monoatomic ion transport; transmembrane transport |
| jg23760 | jg7397 | 0.3608 | 1.32E-05 | | | calcium ion binding | calcium ion binding |
| **jg23761** | **jg7398** | **0.2423** | **1.09E-07** | | | | **protein binding** |
| jg23774 | jg7413 | 0.7765 | 3.23E-88 | non-structural maintenance of chromosomes element 3 homolog [Hermetia illucens] | non-structural maintenance of chromosomes element 3 homolog [Hermetia illucens] | | |

| | | | | | | | |
|---|---|---|---|---|---|---|---|
| jg23812 | jg7459 | 0.2503 | 3.65E-03 | | | RNA methylation; methyltransferase activity; 7-methylguanosine RNA capping | RNA methylation; methyltransferase activity; 7-methylguanosine RNA capping |
| jg23813 | jg7460 | 0.4410 | 7.23E-05 | | | mRNA splicing, via spliceosome; spliceosomal snRNP assembly | mRNA splicing, via spliceosome; spliceosomal snRNP assembly |
| jg23818 | jg7465 | 0.2635 | 2.44E-03 | | | thiol oxidase activity; flavin-dependent sulfhydryl oxidase activity | thiol oxidase activity; flavin-dependent sulfhydryl oxidase activity |
| jg23833 | jg7479 | 6.2154 | 2.29E-116 | L-dopachrome tautomerase yellow-f2 [Ceratitis capitata] | L-dopachrome tautomerase yellow-f2 [Ceratitis capitata] | | |
| jg23916 | jg7560 | 0.2160 | 8.90E-03 | | | glucosamine 6-phosphate N-acetyltransferase activity; UDP-N-acetylglucosamine biosynthetic process; acetyltransferase activity | acetyltransferase activity; glucosamine 6-phosphate N-acetyltransferase activity; UDP-N-acetylglucosamine biosynthetic process |
| jg23954 | jg7598 | 0.3103 | 4.45E-05 | | | beta-1,4-mannosyltransferase activity | beta-1,4-mannosyltransferase activity |
| jg23959 | jg7603 | 0.5370 | 1.21E-07 | | | RNA binding; nucleic acid binding | RNA binding; nucleic acid binding |
| jg23976 | jg7618 | 0.5743 | 1.98E-06 | | | NAD+ binding | NAD+ binding |
| jg24003 | jg7641 | 0.3966 | 3.15E-06 | | | hydrolase activity, hydrolyzing O-glycosyl compounds; carbohydrate metabolic process; beta-N-acetylhexosaminidase activity | hydrolase activity, hydrolyzing O-glycosyl compounds; carbohydrate metabolic process; beta-N-acetylhexosaminidase activity |
| **jg24045** | **jg7693** | **8.7896** | **/** | **nucleoporin NUP145 [Lingula anatina]** | | | |
| **jg24120** | **jg7765** | **4.4213** | **8.22E-208** | | | **odorant binding** | **serine-type endopeptidase activity; odorant binding** |
| **jg24352** | **jg8129** | **4.8581** | **2.53E-03** | | | **protein binding; GTP binding** | |
| jg24354 | jg8136 | 1.2979 | 3.80E-03 | | | protein binding | protein binding |
| **jg24355** | **jg8138** | **2.3219** | **2.55E-02** | | **unconventional myosin-XVIIIa-like [Culex pipiens pallens]** | **multivesicular body sorting pathway** | |

**Table S10.** List of autosomal genes found to be significantly differentially deposited into the eggs of gynogenic and androgenic females, and the chromosome it's found on. Log fold change (logFC) and the false discovery rate (i.e. adjusted p-value) is given to 4 decimal places. Column 5 contains BLAST results for the protein to the entire Non-redundant (NR) Protein database, excluding *Bradysia coprophila*. Column 6 contains GO terms that result from a comprehensive InterProScan to obtain protein domain-level predictions and other protein/peptide/domain motif analyses to predict functional information. Many hits to the NCBI database were to hypothetical/uncharacterised proteins, and these were left out. Any genes there are significantly differentially expressed but have no hits to the NCBI database/hits only to hypothetical proteins, and no GO term results from the InterProScan, are not shown in this table.

| Gene ID | Chromo-some | logFC (4 d.p.) | FDR (4 d.p.) | NCBI best-match protein | InterProScan GO Term function |
|---|---|---|---|---|---|
| jg166 | II | 3.4318 | 0.0359 | piggyBac transposable element-derived protein 3-like [Portunus trituberculatus] | |
| jg336 | II | 1.0786 | 0.0260 | | ATP binding; ATP hydrolysis activity; microtubule binding; microtubule severing ATPase activity; microtubule severing |
| jg1130 | II | 0.5153 | 0.0204 | | monooxygenase activity; iron ion binding; oxidoreductase activity, acting on paired donors, with incorporation or reduction of molecular oxygen; heme binding |
| jg1899 | II | 0.9602 | 0.0033 | | acyltransferase activity; transferase activity; oxidoreductase activity |
| jg2861 | II | 3.9020 | 0.0000 | | monooxygenase activity; iron ion binding; oxidoreductase activity, acting on paired donors, with incorporation or reduction of molecular oxygen; heme binding |
| jg3662 | II | -0.4880 | 0.0222 | 1,2-dihydroxy-3-keto-5-methylthiopentene dioxygenase [Culex pipiens pallens] | acireductone dioxygenase [iron(II)-requiring] activity |
| jg8652 | IV | 1.9927 | 0.0004 | | intraciliary transport |
| jg9168 | IV | -0.8711 | 0.0169 | | acyltransferase activity; acyltransferase activity, transferring groups other than amino-acyl groups |
| jg10423 | IV | 0.6107 | 0.0375 | protein spartin [Cryptotermes secundus] | |
| jg10711 | IV | 1.7972 | 0.0490 | | protein binding |
| jg11334 | IV | 3.3406 | 0.0006 | | nucleic acid binding |
| jg11564 | IV | 3.5783 | 0.0000 | | catalytic activity; iron-sulfur cluster binding |
| jg11827 | IV | 3.8000 | 0.0000 | | antiporter activity; xenobiotic transmembrane transporter activity; transmembrane transport |
| jg12773 | IV | -0.6973 | 0.0234 | | nicotinate-nucleotide diphosphorylase (carboxylating) activity; NAD biosynthetic process; pentosyltransferase activity |
| jg13006 | IV | 1.0207 | 0.0026 | | protein binding; calcium ion binding |
| jg15917 | III | 1.4607 | 0.0004 | | ribonuclease III activity; RNA processing |

| | | | | | |
|---|---|---|---|---|---|
| jg16549 | III | 1.9684 | 0.0375 | | endopeptidase inhibitor activity |
| jg18809 | III | 1.2585 | 0.0004 | | nucleotidyltransferase activity |
| jg19292 | III | 6.7064 | 0.0000 | | RNA binding; adenosine deaminase activity; RNA processing; protein binding |
| jg19650 | III | 0.5175 | 0.0154 | | protein binding |
| jg20813 | III | 0.5289 | 0.0096 | | metal ion binding |
| jg20850 | III | 2.3942 | 0.0134 | | protein binding |
